# Complexes between Netrin G Ligands and Chiral Nanoparticles Promote Axons Regeneration under Near-Infrared Illumination

**DOI:** 10.1101/2025.07.18.665327

**Authors:** Aihua Qu, Maozhong Sun, Xinxin Xu, Si Li, Felippe Mariano Colombari, Wenxiong Shi, Changlong Hao, Baimei Shi, Jingqi Dong, André F. de Moura, Michael Veksler, Chuanlai Xu, Liguang Xu, Nicholas A. Kotov, Hua Kuang

## Abstract

Chiral nanoparticles combine functionalities of inorganic materials and large biomacromolecules enabling stimulation of neurons. However, multiple challenges remain in their use for central nervous system including identifying suitable cell signaling pathways and traversing the blood-brain barrier. In this study, we show that glutathione-coated Ni(OH)₂ nanoparticles form nanoscale complexes with netrin-G1 ligands (NGL-1), critical for neuronal regeneration. NGL-1 features a semispherical pocket with a 2.5 nm radius of curvature, fitting well with nanoparticles sized 3 ± 1.2 nm. *D*-NPs with *D*-glutathione surface ligands activate *D*-glutathione receptors on epithelial cells, facilitating their transport into the brain. When illuminated with 980 nm light, the nanoparticle-protein complex stimulates axon regeneration through localized IGF-1 production. This approach successfully regenerated (a) hippocampal neurons in Alzheimer’s disease mice and (b) dorsal root ganglion neurons in spinal cord-injured mice. The nanoparticles were thoroughly tested for safety and excreted intact. The local, rather than systemic, IGF-1 upregulation minimizes its side effects.

---

Systemic delivery of large biomacromolecules with nanoscale dimensions into the central nervous system is impeded by their enzymatic degradation in the bloodstream. Furthermore, these molecules must traverse the tight intercellular junctions of the blood-brain barrier (BBB) and blood-spinal cord barrier (BSCB), which are highly restrictive for large biomolecules.^1–5^ Specially engineered liposomes, exosomes, and polymeric and lipid nanoparticles (NPs) can cross these barriers, carrying biomolecular cargo while protecting it from degradation.^6–8^ For example, in Alzheimer’s disease (AD) treatments using anti-Aβ antibodies, encapsulation can lead to conformational changes that compromise these large biomacromolecules’ ability to recognize amyloid plaques.^9^ The lipid-based carriers are also known to release cargo prematurely before crossing the BBB.^10,11^ The challenges posed by often contrarian requirements for systemic stability, efficient transport, and biological activity of nanoscale biomolecules have prompted the search for alternatives to existing drug delivery concepts for the central nervous system.^12,13^

Inorganic NPs can interact strongly with biomolecules, forming both specific^14^ and non-specific complexes.^15^ The latter, also known as protein coronas, may contain multiple loosely bound biomolecules and are chemically robust enough to endure transport through blood, lymph, or cerebrospinal fluid.^16,17^ Inorganic NPs can also absorb or emit light in the red and near infrared (NIR) spectrum, facilitating trans-BBB transport and neuronal stimulation via biologically safe NIR photons^18–24^ and other electromagnetic fields.^25–28^ Although limited, evidence suggests that inorganic NPs can directly traverse the tight junctions of the BBB and BSCB.^6,7,29^ Despite these advantages, typical inorganic NPs lack the bioactivity and specificity of biomacromolecules, resulting in poor tissue-targeting ability^8^ as they do not form nanoscale complexes with specific protein targets.^30^ Attaching antibodies, viral peptides, and other biomacromolecules to NPs can enhance biospecificity,^29,31,32^ but reintroduces the enzymatic vulnerability characteristic of large biomacromolecules.

Bioinspired chiral NPs offer a new approach to address these biomedical challenges^33^ by fine-tuning their biological interactions with amino acids,^23,34,35^ peptide assemblies,^29,36,37^ and cells.^38–40^ The tight coating of short amino acids^41,42^ or dipeptides^38,43^ is likely to be more enzymatically stable than large biomacromolecules, while enabling collective supramolecular interactions with specific proteins. However, many uncertainties remain, including circulation time, systemic biocompatibility, and potential side effects^44^. Additionally, the receptors and mechanisms through which NPs stimulate neural activity are still unknown.

Netrin-G1 ligand (NGL-1) has recently attracted considerable attention as a component of the cell signaling pathway that regulates axon growth.^45–47^ NGL-1, a transmembrane protein abundant in neuronal dendrites, features an arc-shaped leucine-rich repeat (LRR) region with a geometry conducive to docking spheroidal or elliptical NPs **(Figure S1)**. However, geometric fit alone is insufficient for precise targeting as many receptors in neurons and other cells also possess semispherical pockets and LRR segments of varying lengths.^48–50^ Chiral NPs that combine nanoscale, supramolecular, and molecular features may enhance specificity, offering a desirable combination of therapeutic benefits.

## Synthesis and Characterization of Chiral Nanoparticles

When purified from nickel ions, surfactants, and other admixtures, Ni(OH)_2_ NPs become biocompatible.^51,52^ They also exhibit high colloidal stability in aqueous dispersions,^53^ which made them suitable for this study. Ni(OH)_2_ NPs with a diameter of 3 ± 1.2 nm were synthesized in the presence of glutathione (GSH) **(Figure S2)**.^54^ These NPs will be referred to as *D*-NPs, *L*-NPs, and *rac*-NPs when *D*-, *L*-, and *rac*-GSH were used as surface ligands, respectively.

Molecular dynamics (MD) simulations of the growth mechanism revealed that Ni(OH)_2_ units form a [Ni(OH)_2_]_48_ cluster with all GSH molecules positioned on its surface (**Figure S3, S4**), rather than being intercalated between Ni(OH)_2_ nanosheets. Liquid chromatography-mass spectrometry analysis revealed that approximately 150 GSH molecules are attached to the surface of each NP (**Figure S5**). Chirality is transferred from the GSH surface ligands, which have multiple attachment points to the inorganic core. This is evidenced by strong, broadened peaks in vibrational circular dichroism (VCD) at approximately 1640 and 1590 cm^−1^, corresponding to the C=O stretching vibration of secondary amide groups and the asymmetric stretching of the carboxylate anions in GSH, respectively (**Figure S6**). MD models confirm the transfer of chirality from ligand to core (**Figure S7**), which can be tracked using the average values of the Osipov-Pickup-Dunmur (OPD) chirality index (**Table S1**).

The *D*-, *L*-, and *rac-*NPs displayed nearly identical sizes and size distributions, UV-Vis absorption spectra, zeta-potentials, and small angle X-ray scattering (SAXS) spectra **(Figure S8)**. X-ray photoelectron spectra (XPS) revealed two prominent binding energy peaks for Ni 2p at 870 eV and 852.38 eV, corresponding to Ni^2+^ 2p_1/2_ and Ni^2+^ 2p_3/2_, respectively. Transmission electron microscopy (TEM) images showed lattice spacings of 0.269 nm corresponding to the (100) facets of β-Ni(OH)_2_ **(Figure 1a-1f)**. Energy-dispersive X-ray (EDX) spectroscopy element mapping confirmed the composition of NPs **(Figure 1g)**. The X-ray diffraction (XRD) peaks of the NPs are consistent with β-Ni(OH)_2_ **(Figure 1h** and **Figure S9)** with expected broadening of the 001 peaks at ∼19.1 degree. Importantly, additional reflections for *L-* or *D*-NPs are observed at 38.3 degree that correspond to twist distortions of the inorganic lattice that are obviously smaller for *rac-*NPs and for *D-* and *L-*NPs.^55^

**Figure 1.**
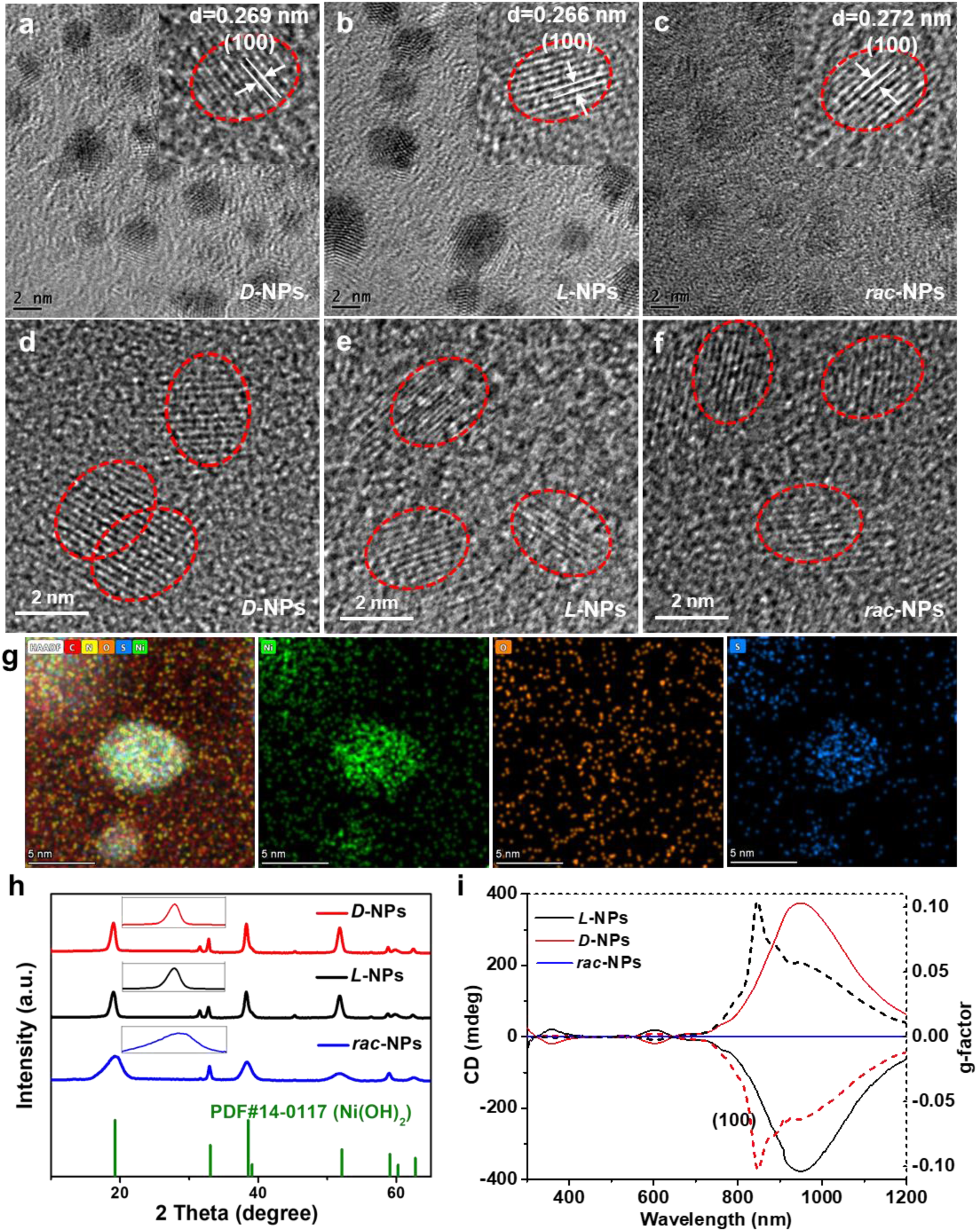
Characterization of chiral GSH-modified NPs of Ni(OH)_2_. **(a-c)** TEM images of *D*-NPs **(a)**, *L*-NPs **(b)**, and *rac*-NPs **(c)**; high resolution TEM images are in the inserts. **(d-f)** Aberration-corrected TEM images of *D*-NPs **(d)**, *L*-NPs **(e)**, and *rac*-NPs **(f)**. **(g)** Energy-dispersive X-ray spectroscopy (EDX) mapping of *D*-NPs. **(h)** XRD spectra of *D*-, *L*-, and *rac*-NPs; Joint Committee on Powder Diffraction Standards, JCPDS, No. 14-0117. **(i)** CD (full lines) and *g*-factor spectra (dashed lines) of *D*-, *L*-, and *rac*-NPs.

The circular dichroism (CD) spectra of *D*-NPs and *L*-NPs displayed mirror-symmetry with strong peaks in the NIR region from 800 nm to 1100 nm, and maximum *g-*factors reaching 0.1 at ∼820 nm–an unusual feature for NPs in the NIR range. As expected, *rac*-NPs were chiroptically silent **(Figure 1i** and **Figure S10)**.

The CD spectra of *D*-NPs showed no significant changes after incubation with PBS, cell culture medium, or plasma, indicating that the surface ligands and NPs are stable against enzymatic degradation both *in vitro* and *in vivo* **(Figure S11**) – a crucial factor for NP delivery to neural tissues.

## Interactions Between Nanoparticles and Netrin G1 Ligands

We found that the binding affinity, *K_a_*, between NPs and NGL-1 is strongly dependent on NP chirality. NGL-1 binds to *D*-NPs with a *K_a_* = 3.48 × 10^7^ M^−1^, which is 34.1-fold and 6.9-fold higher than for *L*-NPs and *rac*-NPs, respectively **(Figure S12)**. The *K_a_* of *D*-NPs is also comparable to that of the NGL-1 and Netrin G1 complex (*K_a_* =0.65×10^7^ M^−1^).^47,56^ In contrast, the *K_a_* between *D*-NPs and Netrin G1 was 9.16 × 10^4^ M^−1^, which is more than 100-fold lower than that for *D*-NPs and NGL-1 **(Figure S13)**.

The strong *K_a_* of *D-*NPs to NGL-1, as well as differences in binding between NPs of different chirality, were also verified by confocal images in living cells (**Figure 2a, b)**. To gain a deeper understanding of the mechanism of chiral effects in NP-NGL-1 interactions, we ran three independent MD simulations for the *L*-NPs, *D*-NPs, and *rac*-NPs interacting with the LRR portion of NGL-1. The average affinity energies obtained from 60-200 ns sections of MD trajectories were –19200 ± 107 kJ/mol for the *rac*-GSH cluster, –19400 ± 128 kJ/mol for the *L*-GSH cluster, and –19500 ± 133 kJ/mol for the *D*-GSH cluster **(Figure 2c-2e** and **Figure S14-S19)**, which is commensurate with the experimental data.

**Figure 2.**
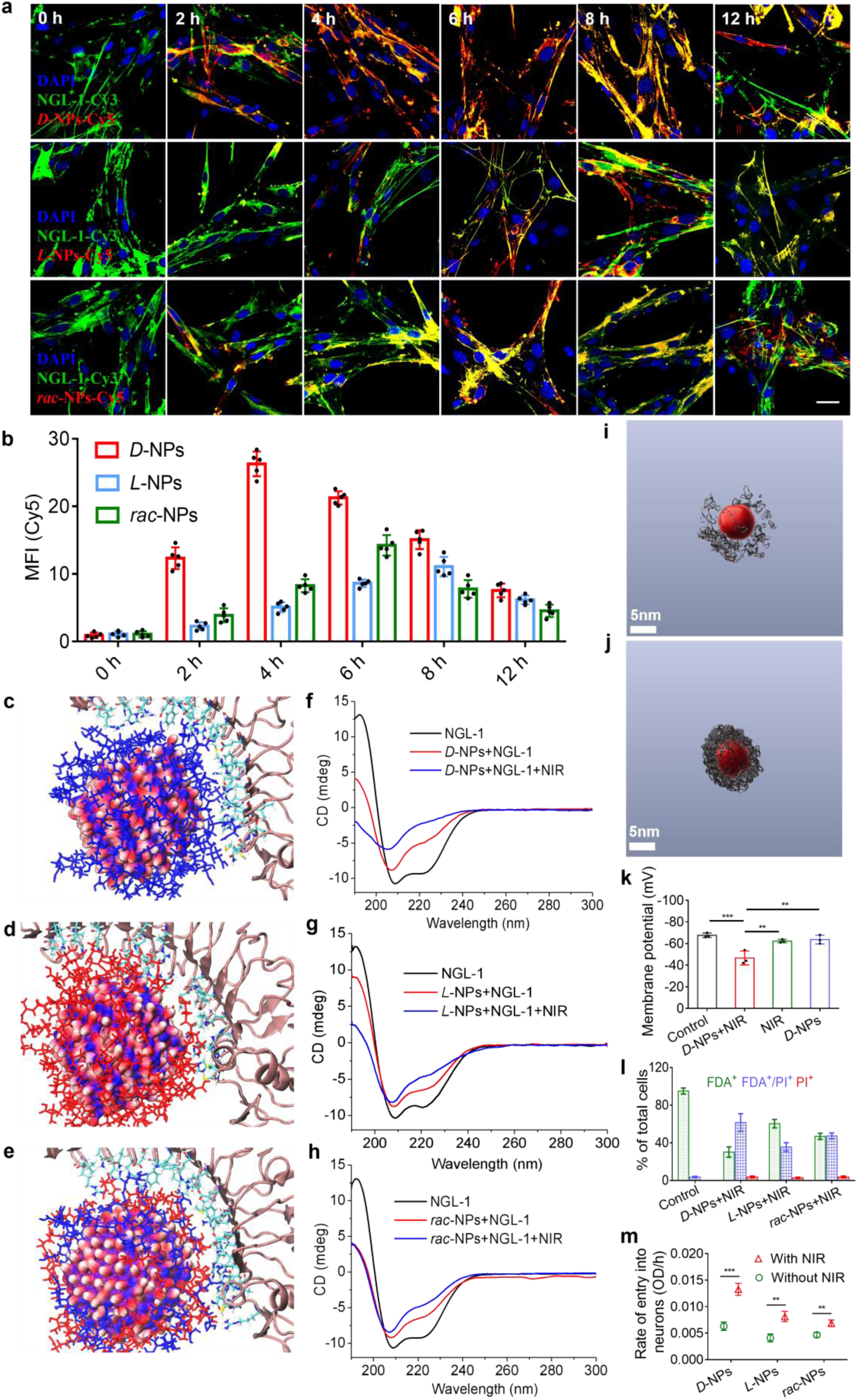
Interactions between NPs and NGL-1. **(a)** Confocal imaging of hippocampus neurons incubated with *D*-NPs, *L*-NPs, and *rac*-NPs under NIR light illumination for different times. Blue: DAPI, Red: *D*-/*L*-/*rac*-NPs-Cy5, Green: NGL-1-Cy3. Scale bar, 20 μm. 500-1000 cells per well for independent observations and randomly select 10-50 cells from any region for imaging and fluorescence statistics. Each experiment was repeated five times. **(b)** Mean fluorescence intensity of Cy5 for hippocampus neurons incubation with *D*-NPs, *L*-NPs, or *rac*-NPs under NIR light illumination of Figure 2a. n = 5 independent experiments. **(c-e)** Protein-NP structures obtained after 900 ns of MD simulation for the **(c)** *D*-NPs, **(d)** *L*-NPs, or **(e)** *rac*-NPs system. Protein chain is presented as the pink cartoon and the residues in closer contact (0.4 nm) with the NP ligands are presented as licorice molecules with default color scheme (carbon atoms are cyan, oxygen atoms are red, nitrogen atoms are blue, and hydrogen atoms are white). The NP is represented as the red (oxygen), blue (nickel), and white (hydrogen) surface with the *D*-GSH ligands represented as blue licorice molecules and the *L*-GSH ligands represented as red licorice molecules. **(f)** CD spectra from 190 nm to 300 nm of NGL-1, NGL-1 with NIR illumination, *D*-NPs with NIR illumination, or *D*-NPs incubated with NGL-1 at a concentration ratio of 500:1 with or without NIR illumination. **(g)** CD spectra from 190 nm to 300 nm of NGL-1, NGL-1 with NIR illumination, *L*-NPs with NIR illumination, or *L*-NPs incubated with NGL-1 at a concentration ratio of 500:1 with or without NIR illumination. **(h)** CD spectra from 190 nm to 300 nm of NGL-1, NGL-1 with NIR illumination, *rac*-NPs with NIR illumination, or *rac*-NPs incubated with NGL-1 at a concentration ratio of 500:1 with or without NIR illumination. **(i)** Cryo-TEM tomography image of *D*-NPs incubated with NGL-1 without NIR illumination. **(j)** Cryo-TEM tomography image of *D*-NPs incubated with NGL-1 with NIR illumination (980 nm, 400 mW/cm^2^, 10 min). **(k)** Membrane potential of hippocampal neurons incubated with *D*-NPs under NIR light (980 nm laser, 400 mW/cm^2^, time of incubation), incubated with *D*-NPs in dark, or only with NIR light. **(l)** Flow cytometry of hippocampal neurons incubation with *D*-/*L*-/*rac*-NPs with NIR light illumination and subsequently stained with PI and FDA. Control represents hippocampal neurons cells without NPs under NIR treatment. FDA^+^, FDA^+^/PI^+^, and PI^+^ denotes the subpopulations of non-permeabilized cells, permeabilized-resealed cells, and dead cells, respectively. **(m)** The rate of *D*-NPs, *L*-NPs, or *rac*-NPs entry into the hippocampal neurons with or without NIR treatment. Data are presented as mean ± s.d. *P < 0.05, **P < 0.01, ***P < 0.001.

In addition to the geometric match between the particle diameter and LRR segment, MD simulations reveal that the binding between NPs and the receptor are due to the collective effect of multiple hydrogen bonds between them. These bonds are dynamic, constantly forming and breaking, but they result in the large difference in affinity between the *L*- and *D*-GSH-NP enantiomers and the NGL-1 (**Figure S20-S23**). The relative probabilities of these interactions may be quantified by measuring hydrogen bond occupancy between NGL-1 and *D*-GSH-NP (**Table S2**), *L*-GSH-NP (**Table S3**) and *rac*-GSH-NP (**Table S4**). Importantly, *D*-GSH-NP demonstrated a more stable hydrogen bond network (**Table S2**) compared to other GSH enantiomers, forming optimal H-bonds as donors to the same amino acids–a stability not observed with *L*-GSH-NP and *rac*-GSH-NP (**Tables S3** and **S4**). The latter showed more dynamic and weaker interactions. For example, the GLY residue of *L*-GSH within *L-*NPs interacted as a donor with GLU181 and as an acceptor with ARG156 in the LRR region (**Table S4**).

Ni(OH)_2_ NPs are spheroidal but lack spherical symmetry due to the inhomogeneous distribution of GSH ligands on their surfaces, which becomes clear when the NP cores are aligned (**Figure S24-Figure S29**). Notably, the *L-* and *D-* enantiomers bind to the protein from the opposite sides, while the *rac-*NP binds through an intermediate region between these opposite sides.

The formation of NP-NGL-1 complexes is also evident from their CD spectra. Protein peaks between 190 nm and 300 nm show clear conformational changes when NGL-1 interacts with *D*-NPs. Similar interactions occur with *L*-NPs and *rac*-NPs, though the peak shifts are smaller than those observed for *D*-NPs **(Figure 2f-2h)**. Minor red shift was also observed for the CD peak of NPs in the 980 nm band, attributed to a protein-induced change in the dielectric constant **(Figure S30)**. However, no significant changes in the CD spectra were observed for either *D*-NPs or NGL-1 upon NIR illumination.

## Effect of Near Infrared Illumination at 980 nm

The CD spectra of NP-NGL-1 complexes showed considerable changes after light exposure **(Figure 2f-2h)**. Since the NPs display a CD peak between 950 and 990 nm, we tested illumination at 980 nm (400 mW/cm^2^) and observed a strong spectral shift after 10 min of illumination **(Figure S30).** XPS spectra showed that the binding energy peaks of chiral NPs remained largely unchanged, suggesting that the valence state of nickel was unaffected by NIR photons **(Figures S8** and **S31)**. The reduction in intensity of the CD bands at 200-240 nm indicates conformational changes in NGL-1 and a tightening of the NP-NGL-1 complex. NIR-induced conformational changes were further confirmed by ^1^H-NMR spectroscopy; spectra recorded before and after illumination of the same amount of NP-NGL-1 complex resulted in additional broadening and decreased intensity of NMR peaks **(Figure S32**), which indicated an increase of the relaxation time is associated with tighter NP-protein binding.

Negatively stained TEM images of a mixture of *D*-NPs and NGL-1 showed increased agglomeration following NIR illumination **(Figure S33)**. Cryo-TEM tomography indicated that the particle size of *D*-NPs remained nearly constant at around 6 nm before and after NIR illumination. The outer layer of the *D*-NPs, representing NGL-1, showed tighter wrapping of the *D*-NPs by NGL-1 after NIR illumination compared to before **(Figure 2i, j, Supplementary Videos 1 and 2)**.

Following prior studies,^57,58^ we also used atomic force microscopy (AFM) to assess the NP interactions with NGL-1 receptors on neurons.^59,60^ *D*-NPs have stronger close-range interactions with cell membranes than *L*-NPs or *rac*-NPs. The interaction force between chiral NPs and neurons under NIR illumination increased by approximately two times **(Figure S34)**. Cell membrane potential **(Figure 2k)** and cell entry efficiency of NPs **(Figure 2l, 2m**; **Figure S35**, and **Figure S36)** also increased under illumination at 980 nm, confirming NIR-induced biological response mediated by *D*-NPs.

To explain the effect of NIR illumination we calculated the energy states of this complex system and determined found that the chiral excitonic level of the NP core and energy levels in the protein are energetically aligned and symmetrically compatible. This leads to local electronic resonances that dissipate the energy of photons into vibrational states of LRR. Therefore, NIR illumination provides kinetic energy for LRR fragments to induce post-docking conformational changes that result in tighter binding **(Figure S37** and **Table S5)**.

## NIR Stimulation of Hippocampal Primary Neurons *in vitro*

Given the conformational changes in NGL-1, we investigated the stimulation of hippocampal primary neurons from wild-type (WT) and 3xTg-AD mice using NPs of different chirality, both with and without NIR illumination. Under identical culture conditions, neurons from 3xTg-AD mice displayed much shorter axons, 37.8 ± 16.4 μm, than those from wild type (WT) mice, 191.2 ± 28.9 μm **(Figure 3a, c)**. We incubated neurons from 3xTg-AD mice with chiral NPs under NIR illumination (10 minutes every 2 hours for the first 12 hours on days 1, 3, and 5) and observed a significant increase in axon length with *D*-NPs (211.8 ± 52.1 μm) compared to *L*-NPs (75.6 ± 14.5 μm) and *rac*-NPs (117.2 ± 22.9 μm). The axon lengths in hippocampal neurons from Alzheimer’s disease (AD) mice cultured under similar conditions but without NPs or without illumination were 47.0 ± 10.1 μm and 48.0 ± 11.6 μm, respectively **(Figure 3a, c** and **Figure S38)**. The efficient axon regeneration stimulated by *D*-NPs under NIR light was confirmed through confocal imaging of growth-associated protein 43 (GAP43) expression, as well as RT-PCR and western blot analysis **(Figure 3b, 3d-3f)**. No axon regeneration was observed when neurons were incubated with *D*-NPs without NIR illumination or exposed to NIR illumination without *D*-NPs **(Figure 3a-3e)**. Importantly, hippocampal neurons treated with *D*-, *L*-, or *rac*-NPs under NIR illumination secreted 5-hydroxytryptamine (5-HT), which likely contributed to the improvement of synaptic plasticity and to maintaining stability in the nervous system.^61^ The levels of other neurotransmitters, such as gamma-aminobutyric acid (GABA), dopamine, and glutamate released from repaired neurons approached those found in normal neurons **(Figure 3g** and **Figures S39)**.

**Figure 3.**
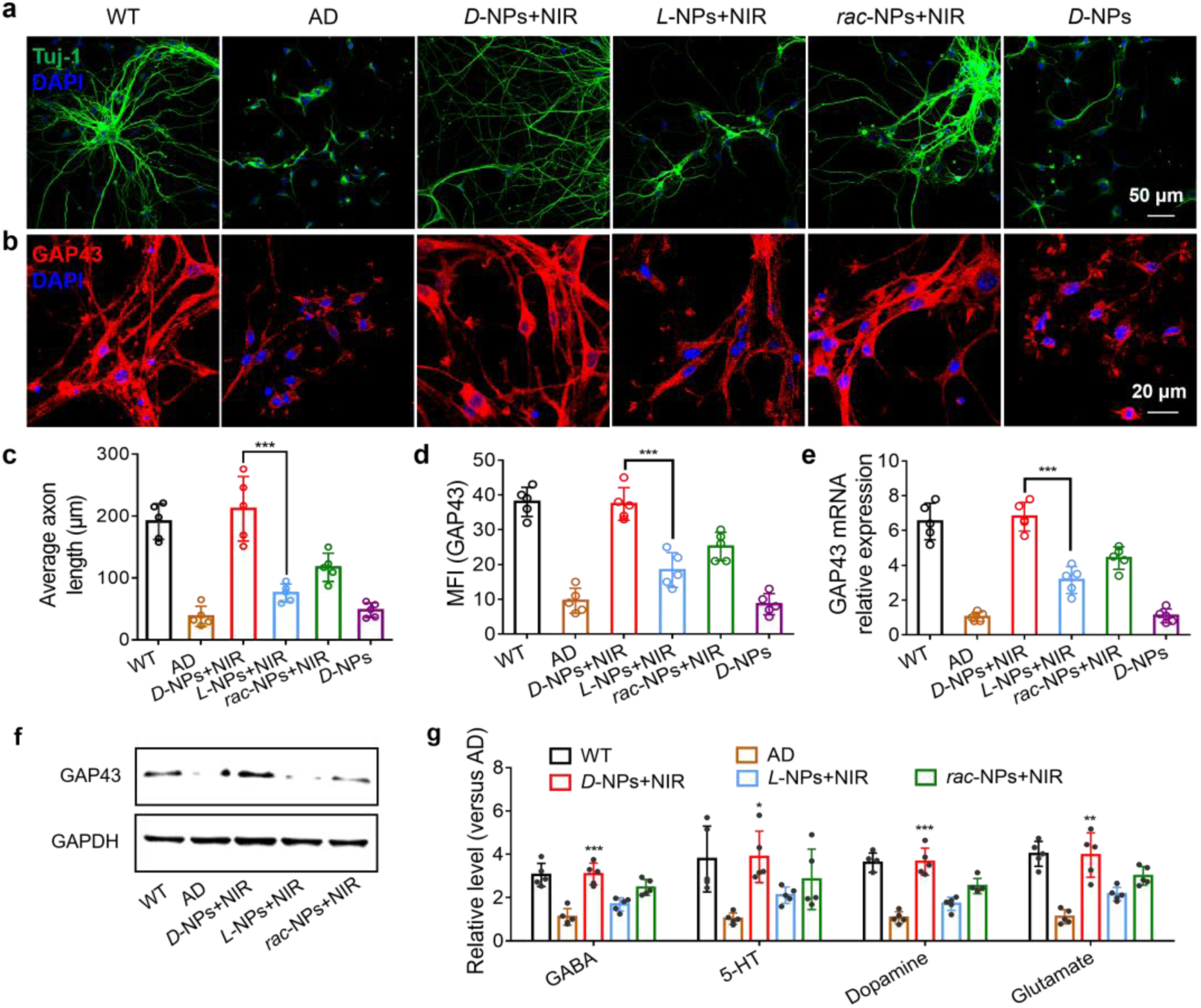
Promoting axon regeneration of primary hippocampal neurons incubated with chiral Ni(OH)_2_ NPs under NIR illumination (980 nm laser; 400 mw/cm^2^) in *vitro*. **(a, b)** Confocal images of hippocampal neurons extracted from WT or AD mice with different treatments. Tuj-1 (green) to label neurons, GAP43 (red) to label axon, and DAPI (blue) to stain nuclei. About 10-50 cells from any region of each well were used for average axon length and fluorescence intensity statistics. Each experiment was repeated five times. **(c)** Quantification of the axonal length of hippocampal neurons (Tuj-1) extracted from WT or AD mice with different treatments from Figure 3a. Mean fluorescence intensity **(d)** and RT-PCR **(e)** analysis of GAP43 in hippocampal neurons extracted from WT or AD mice with different treatments. **(f)** Western blot analysis for GAP43 protein in hippocampal neurons extracted from WT or different months of AD mice with different treatments; GAPDH acts as a control. **(g)** Neurotransmitter release of hippocampal neurons extracted from WT or AD mice with different treatments by ELISA. Data are presented as mean ± s.d. *P < 0.05, **P < 0.01, ***P < 0.001.

To benchmark these findings against possible biological effects of the chiral NPs with different chemistries, we varied their surface ligands and core materials. Aspartic acid (Asp)-Ni(OH)_2_ NPs, histidine (His)-Ni(OH)_2_ NPs, glutathione (GSH)-Cu(OH)_2_ NPs, and Asp-Co(OH)_2_ NPs were synthesized and incubated with neurons under 980 nm illumination. The data in **Figures S40-S44** clearly indicate that *D-*GSH-Ni(OH)_2_ NPs are optimal for axon regeneration. Additionally, the specificity of different chiral NPs for NGL-1 was investigated, further demonstrating that *D-*GSH-Ni(OH)_2_ NPs more efficiently bind with NGL-1 **(Figures 2a, b, and Figure S45)**, thereby activating the downstream signaling pathway and promoting IGF-1 production, which supports axon regeneration. In addition, no increase of reactive oxygen species (ROS) was detected in living cells under NIR with and without NPs **(Figure S46)**.

## *In vitro* Studies of Biological Effects of *D-*NPs

Confocal microscopy images of cell cultures showed that *D*-NPs under 980 nm illumination promote neuronal axon regeneration. Tagging Netrin G1, the receptor for NGL-1, it also became evident that NPs had higher binding efficiency with NGL-1 than with Netrin G1 **(Figure S47)**. Blocking NGL-1 eliminates the effect of NPs and NIR illumination on axonal regeneration **(Figure 4a-4d)**.

**Figure 4.**
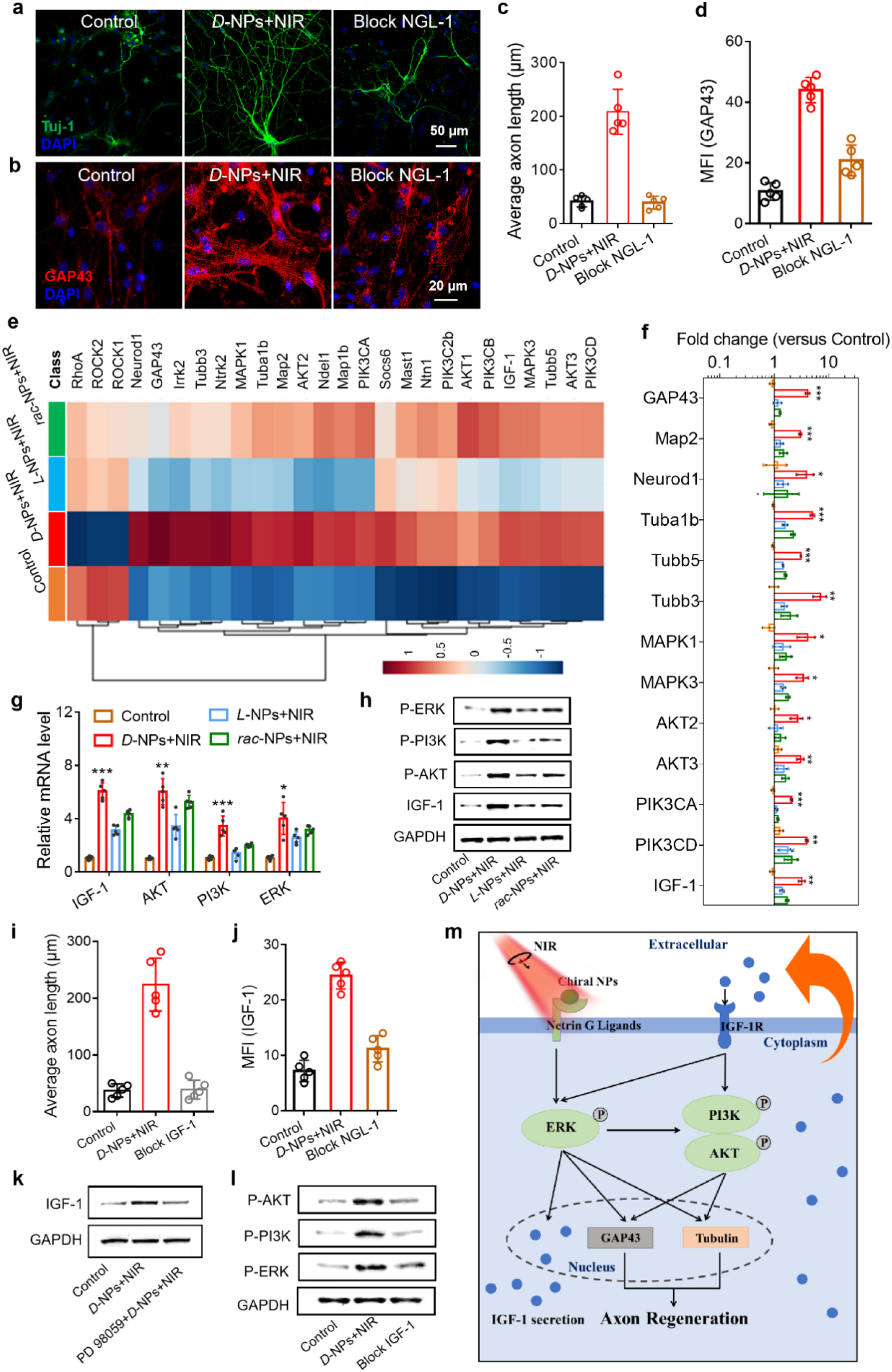
Mechanism of NIR-induced chiral NPs promoting axon regeneration. **(a, b)** Confocal images of hippocampus neurons pretreated with siNGL-1, and then incubated with *D*-NPs under NIR illumination. Tuj-1 (green) to label neurons, GAP43 (red) to label axon, and DAPI (blue) to stain nuclei. About 10-50 cells from any region of each well were used for average axon length and fluorescence intensity statistics. Each experiment was repeated five times. **(c)** Quantification the hippocampal neurons (Tuj-1) axon length pretreated with siNGL-1, and then incubated with *D*-NPs under NIR illumination from Figure 4a. **(d)** Mean fluorescence intensity of GAP43 in hippocampus neurons pretreated with siNGL-1, and then incubated with *D*-NPs under NIR illumination from Figure 4b. **(e)** Global gene-expression pattern of transcription factors in hippocampus neurons with different treatments, the hippocampus neurons extracted from AD mice without any treatment were used as the control group. **(f)** The differentially expressed genes from the heatmap results are expressed as fold change compared with levels in hippocampus neurons with different treatments. **(g)** RT-PCR analysis for IGF-1, P-AKT, P-PI3K, and P-ERK expression in hippocampus neurons with different treatments. **(h)** Western blot analysis for IGF-1, AKT, PI3K, and ERK expression in hippocampus neurons with different treatments. **(i)** Quantification of the hippocampal neurons axon length pretreated with siIGF-1, and then incubated with *D*-NPs under NIR illumination. **(j)** Mean fluorescence intensity of IGF-1 in hippocampus neurons pretreated with siNGL-1, and then incubated with *D*-NPs under NIR illumination. **(k)** Western blot analysis for IGF-1 expression in hippocampus neurons pretreated with PD 98059 (ERK pathway inhibitor), and then incubated with *D*-NPs under NIR illumination. **(l)** Western blot analysis for P-AKT, P-PI3K, and P-ERK expression in hippocampus neurons pretreated with siIGF-1, and then incubated with *D*-NPs under NIR illumination. **(m)** A diagram illustrating the mechanism of neurons axon regeneration via IGF-1 secretion. Data are presented as mean ± s.d. *P < 0.05, **P < 0.01, ***P < 0.001.

We then investigated transcriptome sequencing (RNA-Seq) of living primary hippocampal neurons incubated with *D*-, *L*-, and *rac*-NPs under NIR illumination. The upregulation of genes involved in the phosphatidylinositol 4,5-bisphosphate 3-kinase (PI3K)/protein kinase B (AKT) signaling pathway, the extracellular regulated protein kinase (ERK) signaling pathway, and neuron-associated markers, including Map2, neuronal differentiation 1 (Neurod1), GAP43, and β3-tubulin (Tubb3), was observed. Genes responsible for production of insulin-like growth factor-1 (IGF-1) displayed a significant upregulation as well **(Figure 4e, f)**, which was confirmed by RT-PCR and western blot **(Figure 4g, h)**. IGF-1 promotes synaptogenesis,^62–64^ which emerged as the primary hypothesis for the mechanism of neuronal regeneration by *D-*NPs. To test this hypothesis, we blocked IGF-1 biosynthesis using siRNA, which eliminated the regenerative effect **(Figure 4i** and **Figure S48)**. Similarly, blocking NGL-1, which serves as a cell signaling receptor for NPs, produced the same effect **(Figure 4j)**. The inhibition of the extracellular-signal-regulated kinase (ERK) signaling pathway with PD 98059, through its binding with upstream mitogen-activated protein kinase (MEK), also hindered IGF-1 production **(Figure 4k)**. Thus, the activation of the ERK and PI3K/Akt pathways under NIR illumination led to IGF-1 expression. This conclusion was further confirmed by western blot analysis **(Figure 4l, m)**. We also tested whether the other two Netrin G ligands, that is, NGL-2 and NGL-3, could be involved in axonal regeneration, but blocking NGL-2 and NGL-3 resulted in much smaller biological effects **(Figure S49)**. That is because the LRR segments in NGL-2 and NGL-3 have different 3D conformations, radii of curvature, and amino acid sequences^47^ **(Figure S1** and **Figure S50**) compared to NGL-1, which leads to suboptimal binding of *D-*NPs to these proteins.

## NIR Illumination of *D-*NP-NGL-1 Complexes Promotes Axon Regrowth and Functional Recovery of AD Mice

Neuron loss and dysfunction are associated with typical symptoms of AD.^65,66^ The ability of NPs to increase neuronal connection density in the brain depends on their ability to penetrate the BBB.^4,8^ We compared BBB transport of NPs with various sizes and compositions, as well as solutions of nickel (II) chloride, by intravenously injecting AD mice with the same Ni dose of 0.1 mg/kg. Their biodistribution in brain tissue was detected by inductively coupled plasma mass spectrometry (ICP-MS, **Figure 5a)**. *D*-NPs successfully crossed the BBB, reaching peak concentration in the brain 6 hours after injection. As expected, larger Ni(OH)_2_ NPs with sizes of 80 ± 4.3 nm were unable to cross the BBB.^8^ The transport of 3 ± 1.2 nm *D-*NPs across the BBB was also detected by fluorescence imaging **(Figure 5b).** The retention time of *D-*NPs in the brain was 5 days, significantly longer than the 48-hour retention observed for nickel (II) chloride solution. Additionally, we found that *D-*NPs were cleared from the brain in particle form rather than as Ni^2+^ ions **(Figure S51, S52)**, which is essential for minimizing side effects. Notably the chiral NPs were synthesized in ultrapure water rather than organic solvents to further reduce toxicity. In addition, successive centrifugations were performed to remove unreacted nickel ions, as the carcinogenic potential of Ni is directly related to the bioavailability of Ni^2+^ ions rather than to nickel particles.^67–69^ GSH was used as the chiral ligand because of its high biocompatibility and biosafety, as confirmed by the toxicity test and dosage effect trials. NPs were eventually excreted from the body intact through the kidneys, and to a smaller extent, through the liver, in agreement with previous studies of NPs of similar sizes.^70,71^

**Figure 5.**
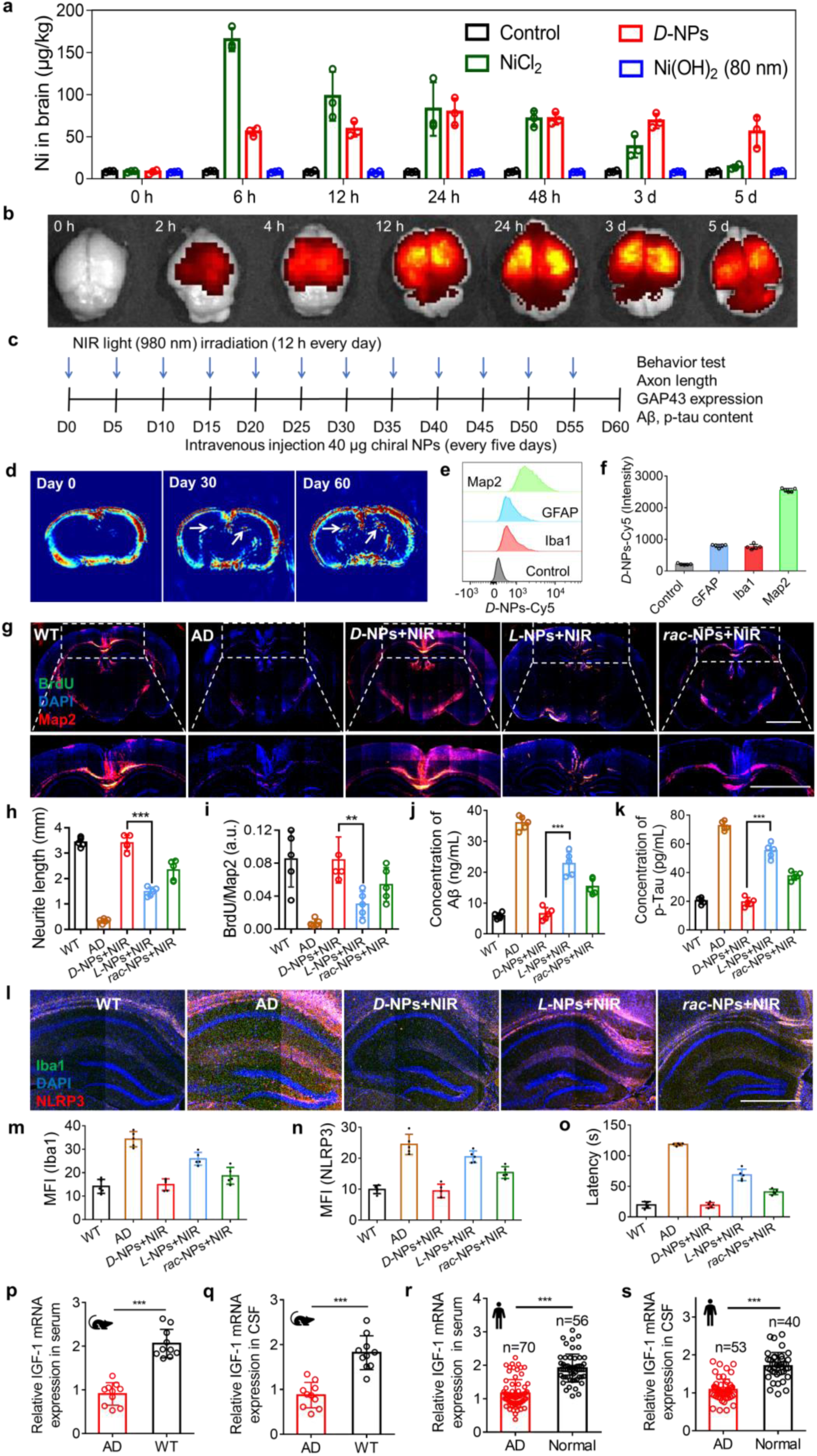
Intravenous injection with chiral Ni(OH)_2_ NPs under NIR illumination (980 nm laser; 600 mw/cm^2^) promotes axon regrowth, ameliorates neuroinflammation *in vivo*, and enhances functional recovery of AD mice. **(a)** ICP-MS analysis the Ni element in the mouse brain after intravenous injection with NiCl_2_, *D*-NPs, or 80 nm Ni(OH)_2_ (at the same Ni dose, 0.1 mg/kg). **(b)** Brain tissue fluorescence imaging of AD mice intravenously injected with *D*-NPs-Cy5 for different times. **(c)** Schematic illustration of injection coordinates of chiral NPs (2 mg/kg, corresponding to Ni dose of 0.1 mg/kg) for intravenous injection in AD mice once every five days for 60 days. **(d)** Photoacoustic images of brain through intravenous injection with *D*-NPs for different times (0, 30, 60 days), arrows represent *D*-NPs material signals. **(e)** Flow cytometry analysis the distribution of *D*-NPs-Cy5 in the neurons (Map2), microglia (Iba1) and astrocytes (GFAP) of the brain tissues though intravenous injection. **(f)** The fluorescence intensity of *D*-NPs-Cy5 in the neurons, microglia and astrocytes of the brain. **(g)** Representative brain section images of WT mice or AD mice with different treatments, immunostained for Map2 (red) to label neuronal dendrites, BrdU (green) to label dividing cells and DAPI (blue) to stain nuclei. Scale bar, 2 mm. **(h)** Quantification the hippocampal neurons (Map2) neurite length of WT mice or AD mice with different treatments from Figure 5g. **(i)** Quantification the ratio of newborn hippocampal neurons of WT mice or AD mice with different treatments from Figure 5g. **(j)** Aβ and **(k)** p-Tau concentrations in cerebrospinal fluid of WT and AD mice with different treatments. **(l)** Representative brain section images of WT mice or AD mice with different treatments, immunostained for NLRP3 (red), Iba1 (green), and DAPI (blue). Scale bar, 1 mm. **(m, n)** Mean fluorescence intensity of Iba1 **(m)** and NLRP3 **(n)** of brain section from Figure 5l. **(o)** The latent period in water maze of the WT mice and AD mice to find the target quadrant after different treatments. **(p, q)** RT-PCR analysis for IGF-1 expression in serum **(p)** and cerebrospinal fluid **(q)** of AD or WT mice. Data are presented as mean ± s.d. (n = 10). **(r, s)** Clinical analysis of relative IGF-1 mRNA expression through RT-PCR in serum **(r)** and cerebrospinal fluid **(s)** of AD patients and healthy humans. Data are presented as mean ± s.d. *P < 0.05, **P < 0.01, ***P < 0.001.

The successful crossing of the BBB by *D-*NPs is attributed to their small size and the activation of GSH transporters **(Figure S53)**.^29^ An *in vitro* BBB model was constructed using human brain endothelial hCMEC/D3 cell lines. This model unequivocally demonstrated that BBB-crossing of *D*-NPs is related to *D*-GSH receptors, because their blocking clearly inhibited NP transport **(Figure S54)**.

The outstanding performance on cross-BBB transport of NPs prompted us to investigate functional recovery in AD mice. 3xTg AD mice were intravenously administered 40 μg of *D*-, *L*-, or *rac*-NPs once every five days up to 60 days, with NIR illumination (980 nm laser; 600 mw/cm^2^) for 12 hours daily **(Figure 5c)**. Treatment was confirmed by photoacoustic (PA) imaging and ICP-MS analysis **(Figure 5d** and **Figure S51)**. In addition, the chiral NPs were completely excreted from the brain within 30 days. Flow cytometry analysis of brain tissue showed that most NPs entered the neurons **(Figure 5e, f)**. Immunofluorescence imaging visualized hippocampal neurons, and neurite length quantification indicated that the neurite length in AD mice treated with D-NPs under NIR illumination (3.4 ± 0.3 mm) reached levels comparable to WT mice (3.43 ± 0.18 mm). In contrast, neurite lengths were 1.4 ± 0.18 mm in AD mice treated with *L*-NPs under NIR and 2.33 ± 0.39 mm with rac-NPs under NIR illumination. Bromodeoxyuridine (BrdU) labeling showed relatively low proportion of newly dividing cells (0.08 ± 0.03), indicating that axon network regeneration was primarily due to cell growth^72^ rather than cell differentiation **(Figure 5g-5i)**. Notably, pathological markers of hyperphosphorylated tau (p-Tau) and amyloid-β (Aβ) were reduced by 73.1% and 81.9%, respectively, in the *D*-NPs under NIR treated AD mice, reaching levels similar to WT mice **(Figure 5j, k)**, which were also validated by immunofluorescence and immunohistochemical analyses **(Figure S55)**, comparable with the previous report.^73^ Nissl staining results confirmed that the nuclei of neurons in the hippocampus were intact and the number of neurons increased after administration of *D*-NPs under NIR illumination **(Figure S55)**.

Excessive microglia activation and resulting neuroinflammation contribute to neuronal cell death.^74^ In AD mice treated with *D*-NPs under NIR illumination, the expression of the microglial activation marker Iba1 and the NLRP3 inflammasome decreased to levels observed in WT mice (**Figures 5l, n, m**). Memory and spatial cognition ability were evaluated by the Morris water maze (MWM) test, and the latency of 3xTg AD mice after *D*-NPs under NIR treatment in finding the platform was equivalent to those of WT mice **(Figure 5o, Supplementary Videos 3, 4,** and **5)**.

Concurrently, the relative expression of IGF-1 mRNA in serum and cerebrospinal fluid (CSF) of AD and WT mice was further evaluated by RT-PCR. IGF-1 levels in the serum and CSF of WT mice were 2.27-fold and 2.08-fold higher, respectively, than in AD mice **(Figure 5p-5s)**. Identical trends were observed in human clinical samples, with IGF-1 levels in serum and CSF higher in healthy individuals compared to AD patients, which further confirmed the developed mechanism.

## Recovery of spinal cord injury *in vitro* and *in vivo*

Axon regeneration is also relevant for peripheral neurons, such dorsal root ganglion (DRG).^75^ Treatment with NIR-excited *D*-NPs promoted axon regeneration and increased GAP43 expression in primary DRG neurons **(Figure 6a-6d** and **Figure S56**). Blocking IGF-1 halted axon regrowth in DRG neurons was consistent with findings in hippocampal neurons. The gene expression changes in DRG neurons mirrored those in hippocampal neurons, indicating a shared cell stimulation pathway **(Figure S57)**. Confocal imaging and western blot analysis confirmed that IGF-1 expression in DRG neurons was driven by NGL-1 activation **(Figure S58** and **Figure 6e)**. Blocking IGF-1 eliminated the biological effects and chiral selectivity between *D-, L-,* and *rac-*NPs **(Figure 6a-6d)**.

**Figure 6.**
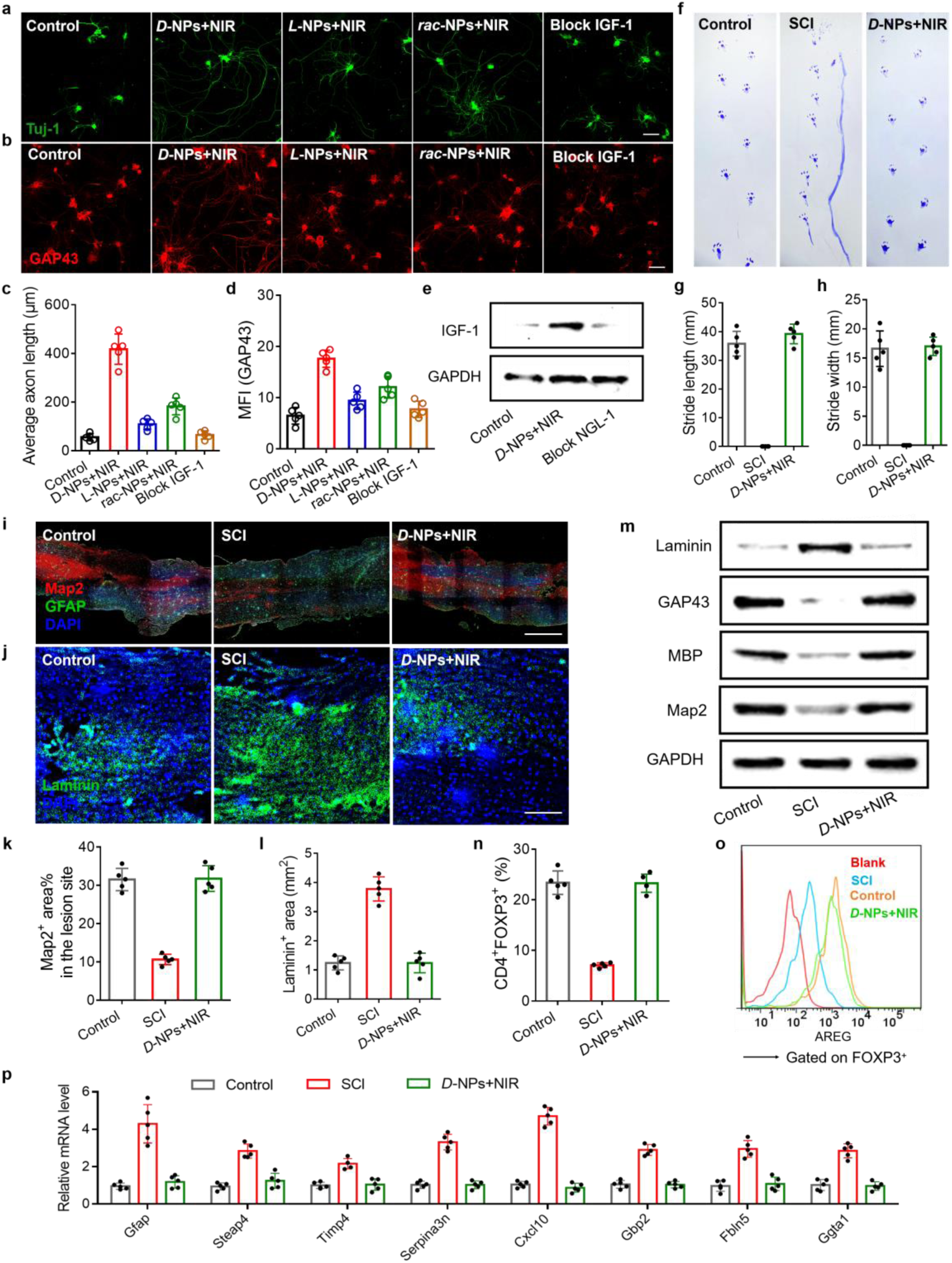
*In vitro* axon regeneration of DRG neurons incubated with *D-*NPs under NIR illumination (980 nm laser; 400 mw/cm^2^); and *in vivo* intravenous injection with chiral NPs under NIR illumination (980 nm laser; 600 mw/cm^2^) **(a, b)** Confocal microscopy images of DRG neurons cells incubated with *D-*NPs under NIR illumination, or pretreated with siIGF-1, and then incubated with *D*-NPs under NIR illumination. βIII-Tuj-1 (green) to label neurons, GAP43 (red) to label axon, and DAPI (blue) to stain nuclei. Control represents DRG neurons without treatment. Scale bar, 50 μm. **(c)** Quantification of the axonal length of DRG neurons (Tuj-1) with different treatments from Figure 6a. **(d)** Mean fluorescence intensity of GAP43 in DRG neurons after different treatments from Figure 6b. **(e)** Western blot analysis for IGF-1 expression in DRG neurons pretreated with siNGL-1, and then incubated with *D*-NPs under NIR illumination. **(f)** Representative footprints of healthy mice (Control), SCI mice, and SCI mice with *D*-NPs under NIR treatment (2 mg/kg, corresponding to Ni dose of 0.1mg/kg). **(g, h)** Bar graphs representing **(g)** stride length in mm and **(h)** stride width in mm of mice mentioned in Figure 6f. **(i)** Fluorescent micrographs of longitudinal spinal cord sections for mice with different treatments mentioned in Figure 6f. GFAP (green) to label astrocytes (green), Map2 (red) to label neuronal dendrites, and DAPI (blue) to stain nuclei. **(j)** Immunolabeling for laminin of longitudinal spinal cord sections in different groups mentioned in Figure 6f. **(k)** Map2-immunopositive^+^ area at the lesion site from Figure 6i. **(l)** Laminin-immunopositive^+^ area from Figure 6j. **(m)** Western blot results for Map2, MBP, laminin, and GAP43 expression in the spinal cord injury site from the healthy mice, SCI mice, and SCI mice with *D*-NPs and NIR treatment. **(n)** The numbers of CD4^+^FOXP3^+^ (Forkhead box protein P3) in the spinal cord injury tissue from healthy mice, SCI mice, and SCI mice with *D*-NPs under NIR treatment. **(o)** Flow cytometric analysis of amphiregulin (AREG) expression levels in Treg cells in conditions described in Figure 6f. **(p)** Quantitative PCR analysis of neurotoxic markers of astrocytes in conditions described in Figure 6f. Data are presented as mean ± s.d. *P < 0.05, **P < 0.01, ***P < 0.001.

Thoracic 10 (T10) spinal cord injury (SCI) mice with right hind limb paralysis were tested for functional regeneration.^76^ Healthy female mice, 4-6-weeks old, were anesthetized and positioned prone on the operating table. Following disinfection, a 2 cm incision was made along the midline of the back, centered over the T10 spinous process. The vertebrae were exposed, and the T10 lamina was removed to fully expose the spinal cord. The right T10 spinal cord was then completely removed with iris scissors, which led to paralysis of the ipsilateral hind limb, with no response to stimulation. The mice were intravenously administrated *D*-NPs (40 μg once every five days) under local NIR illumination using 980 nm laser at 600 mW/cm^2^ intensity. After the treatments for 60 days, the mice showed increased stride length and width. They also improved walking balance and coordination compared to injured control mice. These functions were eventually restored to the levels approaching those in healthy mice **(Figure 6f-6h** and **Figure S59)**. In addition, the spontaneous recovery of the SCI mice displayed increased stride length and width at 60 days after injury, but they did not recover to the normal level (**Figure S60**). Spinal cord tissues from damaged sites were immunolabeled using Map2 to visualize neuronal dendrites and laminin to mark fibrotic scarring **(Figure 6i, j)**. The axon regeneration area in the lesion site after treatment was approximately three-fold greater than that in the SCI model group **(Figure 6k)**. Fibrotic scar tissue, an axon growth inhibitor,^77^ was also reduced after treatment with *D-*NPs under NIR illumination **(Figure 6l)**. Myelin basic protein (MBP), the main component in the myelin sheath, was highly expressed in the lesion, due to the remyelination of neuronal axons following treatment with *D*-NPs under NIR illumination **(Figure 6m)**. Forkhead box protein P3 (FOXP3^+^) regulatory T (T_reg_) cells perform a vital role in the repair and regeneration of injured tissue.^78,79^ Flow cytometric analysis demonstrated that the numbers of CD4^+^FOXP3^+^ cells in SCI tissue increased to healthy tissue levels after treatment **(Figure 6n)**. T_reg_ cells expressed high levels of amphiregulin (AREG) and downregulated astrogliosis and promoted neurological recovery on day 60 with the regiment of *D*-NPs + NIR illumination **(Figure 6o)**. The overexpressed neurotoxic markers of astrocytes were decreased in the lesions **(Figure 6p)**. Magnetic resonance imaging (MRI) at 9.4 T (Bruker BioSpec 94/30 USR+PET insert small animal PET/MRI (Bruker, Germany), Jiangsu Institute of Nuclear Medicine) revealed pronounced damage at lesion sites in the controlled SCI mice group, whereas lesion sites in the treatment group were filled with regenerative tissue. In contrast lesion sites in SCI mice from the spontaneous recovery group remained undamaged **(Figure S61, S62)**. Collectively, these data suggest successful axonal regeneration and functional recovery in SCI mice.

In conclusion, this study demonstrates the use of chirality engineering to achieve strong, specific interactions with proteins relevant to BBB crossing and neuronal activation. *D-*NPs with strong optical activity at 900-1000 nm form a complex with the neuronal receptor NGL-1 via its LRR domain. Subsequent illumination with 980 nm photons induces dose-dependent axon growth in both central and peripheral nervous systems, opening the door for further exploration of NPs in neural regeneration. This work provides an alternative to neuron stimulation methods such as acoustic,^80^ auditory,^81^ electric,^82^ magnetic,^83^ and other electromagnetic field treatments.^21^ While valuable, these treatments primarily focus on generating action potentials via different biochemical mechanisms. Direct activation of the NGL-1 pathway enables a localized increase in IGF-1 within tissues rather than systemic elevation, reducing the risk of adverse side effects and improves treatment outcomes **(Figure S63)**.

## Supporting information

Supplementary methods, data, comments, references

Videos of mice before and after treatment with NP+NIR; WT included. In triplicate

TEM tomography of NP forming a ‘loose’ complex before NIR

TEM tomography of NP forming a ‘tight’ complex after NIR

## ACKNOWLEDGMENTS

This work is financially supported by the National Natural Science Foundation of China (21925402, 92156003, 32071400, 22207045). The N.A.K and M.V. are grateful for the financial support from National Science Foundation (NSF) and specifically for grant #2243104, Center for Complex Particle Systems (COMPASS); grant #2317423 *Lock-and-Key Interactions of Proteins and Chiral Nanoparticles*, and grant #2418861 CBET-EPSRC *Chiroptical Second-Harmonic Scattering of Nanostructures and Their Biocomplexes.* A.F.M. is grateful to National Council for Scientific and Technological Development (311353/2019-3) and The São Paulo Research Foundation (2013/07296-2) for financial support. We are grateful to the High-Performance Computing resources provided by the SDumont supercomputer at the National Laboratory for Scientific Computing (https://sdumont.lncc.br).

## DATA AVAILABILITY

The raw data and videos are available at *figshare* server: private link: https://figshare.com/s/bc7d2ae23668ccf5ad5b; DOI: https://doi.org/10.6084/m9.figshare.29578067;

## REFERENCES

1 Mahar, M., and Cavalli, V. (2018). Intrinsic mechanisms of neuronal axon regeneration. Nat Rev Neurosci 19, 323–337. 10.1038/s41583-018-0001-8.

2. Poplawski, G.H.D., Kawaguchi, R., Van Niekerk, E., Lu, P., Mehta, N., Canete, P., Lie, R., Dragatsis, I., Meves, J.M., Zheng, B., et al. (2020). Injured adult neurons regress to an embryonic transcriptional growth state. Nature 581, 77–82. 10.1038/s41586-020-2200-5.

3 Varadarajan, S.G., Hunyara, J.L., Hamilton, N.R., Kolodkin, A.L., and Huberman, A.D. (2022). Central nervous system regeneration. Cell 185, 77–94. 10.1016/j.cell.2021.10.029.

4 Nance, E., Pun, S.H., Saigal, R., and Sellers, D.L. (2022). Drug delivery to the central nervous system. Nat Rev Mater 7, 314–331. 10.1038/s41578-021-00394-w.

5 Riikonen, R. (2006). Insulin-Like Growth Factor Delivery Across the Blood-Brain Barrier. Chemotherapy 52, 279–281. 10.1159/000095957.

6 Zhou, Y., Peng, Z., Seven, E.S., and Leblanc, R.M. (2018). Crossing the blood-brain barrier with nanoparticles. J Control Release 270, 290–303. 10.1016/j.jconrel.2017.12.015.

7 Ding, S., Khan, A.I., Cai, X., Song, Y., Lyu, Z., Du, D., Dutta, P., and Lin, Y. (2020). Overcoming blood–brain barrier transport: Advances in nanoparticle-based drug delivery strategies. Mater Today 37, 112–125. 10.1016/j.mattod.2020.02.001.

8 Terstappen, G.C., Meyer, A.H., Bell, R.D., and Zhang, W. (2021). Strategies for delivering therapeutics across the blood-brain barrier. Nat Rev Drug Discov 20, 362–383. 10.1038/s41573-021-00139-y.

9. van Dyck, C.H. (2018). Anti-Amyloid-β Monoclonal Antibodies for Alzheimer’s Disease: Pitfalls and Promise. Biol Psychiatry 83, 311–319. 10.1016/j.biopsych.2017.08.010.

10 Lu, C.-T., Zhao, Y.-Z., Wong, H.L., Cai, J., Peng, L., and Tian, X.-Q. (2014). Current approaches to enhance CNS delivery of drugs across the brain barriers. Int J Nanomedicine 9, 2241–2257. 10.2147/ijn.S61288.

11 Liu, B., and Thayumanavan, S. (2019). Three-Component Sequential Reactions for Polymeric Nanoparticles with Tailorable Core and Surface Functionalities. Chem 5, 3166–3183. 10.1016/j.chempr.2019.09.001.

12 Gao, J., Xia, Z., Gunasekar, S., Jiang, C., Karp, J.M., and Joshi, N. (2024). Precision drug delivery to the central nervous system using engineered nanoparticles. Nat Rev Mater 9, 567–588. 10.1038/s41578-024-00695-w.

13 Wu, D., Chen, Q., Chen, X., Han, F., Chen, Z., and Wang, Y. (2023). The blood–brain barrier: Structure, regulation and drug delivery. Signal Transduct Target Ther 8, 217. 10.1038/s41392-023-01481-w.

14 Cha, M., Emre, E.S.T., Xiao, X., Kim, J.-Y., Bogdan, P., VanEpps, J.S., Violi, A., and Kotov, N.A. (2022). Unifying structural descriptors for biological and bioinspired nanoscale complexes. Nat Comput Sci 2, 243–252. 10.1038/s43588-022-00229-w.

15 Xu, M., Soliman, M.G., Sun, X., Pelaz, B., Feliu, N., Parak, W.J., and Liu, S. (2018). How Entanglement of Different Physicochemical Properties Complicates the Prediction of in Vitro and in Vivo Interactions of Gold Nanoparticles. ACS Nano 12, 10104–10113. 10.1021/acsnano.8b04906.

16 Janjua, T.I., Cao, Y., Ahmed-Cox, A., Raza, A., Moniruzzaman, M., Akhter, D.T., Fletcher, N.L., Kavallaris, M., Thurecht, K.J., and Popat, A. (2023). Efficient delivery of Temozolomide using ultrasmall large-pore silica nanoparticles for glioblastoma. J Control Release 357, 161–174. 10.1016/j.jconrel.2023.03.040.

17 Cai, Q., Li, X., Xiong, H., Fan, H., Gao, X., Vemireddy, V., Margolis, R., Li, J., Ge, X., Giannotta, M., et al. (2023). Optical blood-brain-tumor barrier modulation expands therapeutic options for glioblastoma treatment. Nat Commun 14, 4934. 10.1038/s41467-023-40579-1.

18 Cai, Y., Wei, Z., Song, C., Tang, C., Han, W., and Dong, X. (2019). Optical nano-agents in the second near-infrared window for biomedical applications. Chem Soc Rev 48, 22–37. 10.1039/c8cs00494c.

19 Qu, A., Sun, M., Kim, J.Y., Xu, L., Hao, C., Ma, W., Wu, X., Liu, X., Kuang, H., Kotov, N.A., et al. (2021). Stimulation of neural stem cell differentiation by circularly polarized light transduced by chiral nanoassemblies. Nat Biomed Eng 5, 103–113. 10.1038/s41551-020-00634-4.

20 Chen, S., Weitemier, A.Z., Zeng, X., He, L., Wang, X., Tao, Y., Huang, A.J.Y., Hashimotodani, Y., Kano, M., Iwasaki, H., et al. (2018). Near-infrared deep brain stimulation via upconversion nanoparticle mediated optogenetics. Science 359, 679–684. doi:10.1126/science.aaq1144.

21 Iaccarino, H.F., Singer, A.C., Martorell, A.J., Rudenko, A., Gao, F., Gillingham, T.Z., Mathys, H., Seo, J., Kritskiy, O., Abdurrob, F., et al. (2016). Gamma frequency entrainment attenuates amyloid load and modifies microglia. Nature 540, 230–235. 10.1038/nature20587.

22 Pappas, T.C., Wickramanyake, W.M.S., Jan, E., Motamedi, M., Brodwick, M., and Kotov, N.A. (2007). Nanoscale Engineering of a Cellular Interface with Semiconductor Nanoparticle Films for Photoelectric Stimulation of Neurons. Nano Lett 7, 513–519. 10.1021/nl062513v.

23 Cho, N.H., Guerrero-Martínez, A., Ma, J., Bals, S., Kotov, N.A., Liz-Marzán, L.M., and Nam, K.T. (2023). Bioinspired chiral inorganic nanomaterials. Nat Rev Bioeng 1, 88–106. 10.1038/s44222-022-00014-4.

24 Liu, X., Rao, S., Chen, W., Felix, K., Ni, J., Sahasrabudhe, A., Lin, S., Wang, Q., Liu, Y., He, Z., et al. (2023). Fatigue-resistant hydrogel optical fibers enable peripheral nerve optogenetics during locomotion. Nat Methods 20, 1802–1809. 10.1038/s41592-023-02020-9.

25 Romero, G., Park, J., Koehler, F., Pralle, A., and Anikeeva, P. (2022). Modulating cell signalling in vivo with magnetic nanotransducers. Nat Rev Methods Primers 2, 92. 10.1038/s43586-022-00170-2.

26 al., A.M.e. Piezoelectric Nanoparticle-Assisted Wireless Neuronal Stimulation. ACS Nano 9, 7678–7689.

27 Yoo, J., Lee, E., Kim, H.Y., Youn, D.H., Jung, J., Kim, H., Chang, Y., Lee, W., Shin, J., Baek, S., et al. (2017). Electromagnetized gold nanoparticles mediate direct lineage reprogramming into induced dopamine neurons in vivo for Parkinson’s disease therapy. Nat Nanotechnol 12, 1006–1014. 10.1038/nnano.2017.133.

28 Kim, Y.J., Kent, N., Vargas Paniagua, E., Driscoll, N., Tabet, A., Koehler, F., Malkin, E., Frey, E., Manthey, M., Sahasrabudhe, A., et al. (2024). Magnetoelectric nanodiscs enable wireless transgene-free neuromodulation. Nat Nanotechnol. 10.1038/s41565-024-01798-9.

29 Hou, K., Zhao, J., Wang, H., Li, B., Li, K., Shi, X., Wan, K., Ai, J., Lv, J., Wang, D., et al. (2020). Chiral gold nanoparticles enantioselectively rescue memory deficits in a mouse model of Alzheimer’s disease. Nat Commun 11, 4790. 10.1038/s41467-020-18525-2.

30 Gonzalez-Carter, D., Liu, X., Tockary, T.A., Dirisala, A., Toh, K., Anraku, Y., and Kataoka, K. (2020). Targeting nanoparticles to the brain by exploiting the blood-brain barrier impermeability to selectively label the brain endothelium. Proc Natl Acad Sci U S A 117, 19141–19150. 10.1073/pnas.2002016117.

31 Liu, H., Han, Y., Wang, T., Zhang, H., Xu, Q., Yuan, J., and Li, Z. (2020). Targeting Microglia for Therapy of Parkinson’s Disease by Using Biomimetic Ultrasmall Nanoparticles. J Am Chem Soc 142, 21730–21742. 10.1021/jacs.0c09390.

32 Cao, M., Cai, R., Zhao, L., Guo, M., Wang, L., Wang, Y., Zhang, L., Wang, X., Yao, H., Xie, C., et al. (2021). Molybdenum derived from nanomaterials incorporates into molybdenum enzymes and affects their activities in vivo. Nat Nanotechnol 16, 708–716. 10.1038/s41565-021-00856-w.

33 Cho, N.H., Kim, H., Kim, J.W., Lim, Y.-C., Kim, R.M., Lee, Y.H., and Nam, K.T. (2024). Chiral inorganic nanomaterials for biomedical applications. Chem 10, 1052–1070. 10.1016/j.chempr.2023.12.016.

34 Sun, X., Kong, H., Zhou, Q., Tsunega, S., Liu, X., Yang, H., and Jin, R.H. (2020). Chiral Plasmonic Nanoparticle Assisted Raman Enantioselective Recognition. Anal Chem 92, 8015–8020. 10.1021/acs.analchem.0c01311.

35 Sun, M., Xu, L., Qu, A., Zhao, P., Hao, T., Ma, W., Hao, C., Wen, X., Colombari, F.M., de Moura, A.F., et al. (2018). Site-selective photoinduced cleavage and profiling of DNA by chiral semiconductor nanoparticles. Nat Chem 10, 821–830. 10.1038/s41557-018-0083-y.

36 Yoo, S.I., Yang, M., Brender, J.R., Subramanian, V., Sun, K., Joo, N.E., Jeong, S.H., Ramamoorthy, A., and Kotov, N.A. (2011). Inhibition of Amyloid Peptide Fibrillation by Inorganic Nanoparticles: Functional Similarities with Proteins. Angew Chem Int Ed 50, 5110–5115. 10.1002/anie.201007824.

37 Wang, X., Wang, M., Lei, R., Zhu, S.F., Zhao, Y., and Chen, C. (2017). Chiral Surface of Nanoparticles Determines the Orientation of Adsorbed Transferrin and Its Interaction with Receptors. ACS Nano 11, 4606–4616. 10.1021/acsnano.7b00200.

38 Xu, L., Wang, X., Wang, W., Sun, M., Choi, W.J., Kim, J.Y., Hao, C., Li, S., Qu, A., Lu, M., et al. (2022). Enantiomer-dependent immunological response to chiral nanoparticles. Nature 601, 366–373. 10.1038/s41586-021-04243-2.

39 Yan, H., Cacioppo, M., Megahed, S., Arcudi, F., Đorđević, L., Zhu, D., Schulz, F., Prato, M., Parak, W.J., and Feliu, N. (2021). Influence of the chirality of carbon nanodots on their interaction with proteins and cells. Nat Commun 12, 7208. 10.1038/s41467-021-27406-1.

40 Zhao, X., Zang, S.Q., and Chen, X. (2020). Stereospecific interactions between chiral inorganic nanomaterials and biological systems. Chem Soc Rev 49, 2481–2503. 10.1039/d0cs00093k.

41 Lee, H.E., Ahn, H.Y., Mun, J., Lee, Y.Y., Kim, M., Cho, N.H., Chang, K., Kim, W.S., Rho, J., and Nam, K.T. (2018). Amino-acid- and peptide-directed synthesis of chiral plasmonic gold nanoparticles. Nature 556, 360–365. 10.1038/s41586-018-0034-1.

42 Jiang, W.F., Qu, Z.B., Kumar, P., Vecchio, D., Wang, Y.F., Ma, Y., Bahng, J.H., Bernardino, K., Gomes, W.R., Colombari, F.M., et al. (2020). Emergence of complexity inhierarchically organized chiral particles. Science 368, 642–648. 10.1126/science.aaz7949.

43 Kim, H., Im, S.W., Cho, N.H., Seo, D.H., Kim, R.M., Lim, Y.C., Lee, H.E., Ahn, H.Y., and Nam, K.T. (2020). gamma-Glutamylcysteine- and Cysteinylglycine-Directed Growth of Chiral Gold Nanoparticles and their Crystallographic Analysis. Angew Chem Int Ed Engl 59, 12976–12983. 10.1002/anie.202003760.

44 Neus Feliu, and Parak, W.J. (2024). Developing future nanomedicines. Science 384, 385–386.

45 Fujita, Y., Nakanishi, T., Ueno, M., Itohara, S., and Yamashita, T. (2020). Netrin-G1 Regulates Microglial Accumulation along Axons and Supports the Survival of Layer V Neurons in the Postnatal Mouse Brain. Cell Rep 31, 107580. 10.1016/j.celrep.2020.107580.

46 Lin, J.C., Ho, W.H., Gurney, A., and Rosenthal, A. (2003). The netrin-G1 ligand NGL-1 promotes the outgrowth of thalamocortical axons. Nat Neurosci 6, 1270–1276. 10.1038/nn1148.

47 Nishimura-Akiyoshi, S., Niimi, K., Nakashiba, T., and Itohara, S. (2007). Axonal netrin-Gs transneuronally determine lamina-specific subdendritic segments. Proc Natl Acad Sci U S A 104, 14801–14806. doi:10.1073/pnas.0706919104.

48 Morlot, C., Thielens, N.M., Ravelli, R.B.G., Hemrika, W., Romijn, R.A., Gros, P., Cusack, S., and McCarthy, A.A. (2007). Structural insights into the Slit-Robo complex. Proc Natl Acad Sci U S A 104, 14923–14928. doi:10.1073/pnas.0705310104.

49 Ko, J., and Kim, E. (2007). Leucine-rich repeat proteins of synapses. J Neurosci Res 85, 2824–2832. 10.1002/jnr.21306.

50 Soler-Llavina, G.J., Arstikaitis, P., Morishita, W., Ahmad, M., Sudhof, T.C., and Malenka, R.C. (2013). Leucine-rich repeat transmembrane proteins are essential for maintenance of long-term potentiation. Neuron 79, 439–446. 10.1016/j.neuron.2013.06.007.

51 Yang, L., Zhu, X., Xu, T., Han, F., Liu, G., Bu, Y., Zhang, J., Zhang, F., Zhou, H., and Xie, Y. (2020). Defect-engineered transition metal hydroxide nanosheets realizing tumor-microenvironment-responsive multimodal-imaging-guided NIR-II photothermal therapy. J Mater Chem B 8, 8323–8336. 10.1039/d0tb01608j.

52 Lin, J., Dong, L., Liu, Y.-M., Hu, Y., Jiang, C., Liu, K., Liu, L., Song, Y.-H., Sun, M., Xiang, X.-C., et al. (2023). Nickle-cobalt alloy nanocrystals inhibit activation of inflammasomes. Natl Sci Rev 10, nwad179. 10.1093/nsr/nwad179.

53 Chen, Z., Nai, J., Ma, H., and Li, Z. (2014). Nickel hydroxide nanocrystals-modified glassy carbon electrodes for sensitive l-histidine detection. Electrochim Acta 116, 258–262. 10.1016/j.electacta.2013.10.153.

54 Li, C., Li, S., Zhao, J., Sun, M., Wang, W., Lu, M., Qu, A., Hao, C., Chen, C., Xu, C., et al. (2022). Ultrasmall Magneto-chiral Cobalt Hydroxide Nanoparticles Enable Dynamic Detection of Reactive Oxygen Species in Vivo. J Am Chem Soc 144, 1580–1588. 10.1021/jacs.1c09986.

55 Yeom, J., Santos, U.S., Chekini, M., Cha, M., de Moura, A.F., and Kotov, N.A. (2018). Chiromagnetic nanoparticles and gels. Science 359, 309–314. doi:10.1126/science.aao7172.

56 Brasch, J., Harrison, O.J., Ahlsen, G., Liu, Q., and Shapiro, L. (2011). Crystal structure of the ligand binding domain of netrin G2. J Mol Biol 414, 723–734. 10.1016/j.jmb.2011.10.030.

57 Schabert, F.A., Henn, C., and Engel, A. (1995). Native Escherichia coli OmpF porin surfaces probed by atomic force microscopy. Science 268, 92–94. 10.1126/science.7701347.

58 Yersin, A., Hirling, H., Steiner, P., Magnin, S., Regazzi, R., Hüni, B., Huguenot, P., De Los Rios, P., Dietler, G., Catsicas, S., et al. (2003). Interactions between synaptic vesicle fusion proteins explored by atomic force microscopy. Proc Natl Acad Sci U S A 100, 8736–8741. doi:10.1073/pnas.1533137100.

59 Vasir, J.K., and Labhasetwar, V. (2008). Quantification of the force of nanoparticle-cell membrane interactions and its influence on intracellular trafficking of nanoparticles. Biomaterials 29, 4244–4252. 10.1016/j.biomaterials.2008.07.020.

60 Hummer, G., and Szabo, A. (2001). Free energy reconstruction from nonequilibrium single-molecule pulling experiments. Proc Natl Acad Sci U S A 98, 3658–3661. 10.1073/pnas.071034098.

61 Zeng, J., Li, X., Zhang, R., Lv, M., Wang, Y., Tan, K., Xia, X., Wan, J., Jing, M., Zhang, X., et al. (2023). Local 5-HT signaling bi-directionally regulates the coincidence time window for associative learning. Neuron 111, 1–18. 10.1016/j.neuron.2022.12.034.

62 O’Kusky, J.R., Ye, P., and D’Ercole, A.J. (2000). Insulin-like growth factor-I promotes neurogenesis and synaptogenesis in the hippocampal dentate gyrus during postnatal development. J Neurosci 20, 8435–8442. 10.1523/jneurosci.20-22-08435.2000.

63 Ozdinler, P.H., and Macklis, J.D. (2006). IGF-I specifically enhances axon outgrowth of corticospinal motor neurons. Nat Neurosci 9, 1371–1381. 10.1038/nn1789.

64 Zhang, Y., Williams, P.R., Jacobi, A., Wang, C., Goel, A., Hirano, A.A., Brecha, N.C., Kerschensteiner, D., and He, Z. (2019). Elevating Growth Factor Responsiveness and Axon Regeneration by Modulating Presynaptic Inputs. Neuron 103, 39–51.e35. 10.1016/j.neuron.2019.04.033.

65 Richetin, K., Steullet, P., Pachoud, M., Perbet, R., Parietti, E., Maheswaran, M., Eddarkaoui, S., Begard, S., Pythoud, C., Rey, M., et al. (2020). Tau accumulation in astrocytes of the dentate gyrus induces neuronal dysfunction and memory deficits in Alzheimer’s disease. Nat Neurosci 23, 1567–1579. 10.1038/s41593-020-00728-x.

66 Wilson, D.M., 3rd, Cookson, M.R., Van Den Bosch, L., Zetterberg, H., Holtzman, D.M., and Dewachter, I. (2023). Hallmarks of neurodegenerative diseases. Cell 186, 693–714. 10.1016/j.cell.2022.12.032.

67 Mukherjee, A., Latvala, S., Hedberg, J., Di Bucchianico, S., Möller, L., Odnevall Wallinder, I., Elihn, K., and Karlsson, H.L. (2016). Nickel Release, ROS Generation and Toxicity of Ni and NiO Micro- and Nanoparticles. Plos One 11, e0159684. 10.1371/journal.pone.0159684.

68 Cambre, M.H., Holl, N.J., Wang, B., Harper, L., Lee, H.-J., Chusuei, C.C., Hou, F.Y.S., Williams, E.T., Argo, J.D., Pandey, R.R., et al. (2020). Cytotoxicity of NiO and Ni(OH)2 Nanoparticles Is Mediated by Oxidative Stress-Induced Cell Death and Suppression of Cell Proliferation. Int J Mol Sci 21, 2355. 10.3390/ijms21072355.

69 Wu, Y., and Kong, L. (2020). Advance on toxicity of metal nickel nanoparticles. Environ Geochem Heal 42, 2277–2286. 10.1007/s10653-019-00491-4.

70 Du, B., Yu, M., and Zheng, J. (2018). Transport and interactions of nanoparticles in the kidneys. Nat Rev Mater 3, 358–374. 10.1038/s41578-018-0038-3.

71 Soo Choi, H., Liu, W., Misra, P., Tanaka, E., Zimmer, J.P., Itty Ipe, B., Bawendi, M.G., and Frangioni, J.V. (2007). Renal clearance of quantum dots. Nat Biotechnol 25, 1165–1170. 10.1038/nbt1340.

72 Denoth-Lippuner, A., and Jessberger, S. (2021). Formation and integration of new neurons in the adult hippocampus. Nat Rev Neurosci 22, 223–236. 10.1038/s41583-021-00433-z.

73 Carro, E., Trejo, J.L., Gomez-Isla, T., LeRoith, D., and Torres-Aleman, I. (2002). Serum insulin-like growth factor I regulates brain amyloid-β levels. Nature Medicine 8, 1390–1397. 10.1038/nm793.

74 McAlpine, C.S., Park, J., Griciuc, A., Kim, E., Choi, S.H., Iwamoto, Y., Kiss, M.G., Christie, K.A., Vinegoni, C., Poller, W.C., et al. (2021). Astrocytic interleukin-3 programs microglia and limits Alzheimer’s disease. Nature 595, 701–706. 10.1038/s41586-021-03734-6.

75 Hervera, A., De Virgiliis, F., Palmisano, I., Zhou, L., Tantardini, E., Kong, G., Hutson, T., Danzi, M.C., Perry, R.B., Santos, C.X.C., et al. (2018). Reactive oxygen species regulate axonal regeneration through the release of exosomal NADPH oxidase 2 complexes into injured axons. Nat Cell Biol 20, 307–319. 10.1038/s41556-018-0039-x.

76 Anderson, M.A., O’Shea, T.M., Burda, J.E., Ao, Y., Barlatey, S.L., Bernstein, A.M., Kim, J.H., James, N.D., Rogers, A., Kato, B., et al. (2018). Required growth facilitators propel axon regeneration across complete spinal cord injury. Nature 561, 396–400. 10.1038/s41586-018-0467-6.

77 Álvarez, Z., Kolberg-Edelbrock, A.N., Sasselli, I.R., Ortega, J.A., Qiu, R., Syrgiannis, Z., Mirau, P.A., Chen, F., Chin, S.M., Weigand, S., et al. (2021). Bioactive scaffolds with enhanced supramolecular motion promote recovery from spinal cord injury. Science 374, 848–856. doi:10.1126/science.abh3602.

78 Ito, M., Komai, K., Mise-Omata, S., Iizuka-Koga, M., Noguchi, Y., Kondo, T., Sakai, R., Matsuo, K., Nakayama, T., Yoshie, O., et al. (2019). Brain regulatory T cells suppress astrogliosis and potentiate neurological recovery. Nature 565, 246–250. 10.1038/s41586-018-0824-5.

79 Munoz-Rojas, A.R., and Mathis, D. (2021). Tissue regulatory T cells: regulatory chameleons. Nat Rev Immunol 21, 597–611. 10.1038/s41577-021-00519-w.

80 Legon, W., Sato, T.F., Opitz, A., Mueller, J., Barbour, A., Williams, A., and Tyler, W.J. (2014). Transcranial focused ultrasound modulates the activity of primary somatosensory cortex in humans. Nat Neurosci 17, 322–329. 10.1038/nn.3620.

81 Martorell, A.J., Paulson, A.L., Suk, H.-J., Abdurrob, F., Drummond, G.T., Guan, W., Young, J.Z., Kim, D.N.-W., Kritskiy, O., Barker, S.J., et al. (2019). Multi-sensory Gamma Stimulation Ameliorates Alzheimer’s-Associated Pathology and Improves Cognition. Cell 177, 256–271.e222. 10.1016/j.cell.2019.02.014.

82 Dmochowski, J., and Bikson, M. (2017). Noninvasive Neuromodulation Goes Deep. Cell 169, 977–978. 10.1016/j.cell.2017.05.017.

83 Stanley, S.A., Kelly, L., Latcha, K.N., Schmidt, S.F., Yu, X., Nectow, A.R., Sauer, J., Dyke, J.P., Dordick, J.S., and Friedman, J.M. (2016). Bidirectional electromagnetic control of the hypothalamus regulates feeding and metabolism. Nature 531, 647–650. 10.1038/nature17183.

