## Supplementary methods, data, comments, references for "Complexes between Netrin G Ligands and Chiral Nanoparticles Promote Axons Regeneration under Near-Infrared Illumination"

**Supplementary Information**

**1. Materials**

All reagents and chemicals used in this study were purchased from Sigma-Aldrich (St. Louis, MO, USA) and were analytical grade, unless otherwise stated. The deionized (DI) water was obtained with a Milli-Q device (18.2 MΩ; Millipore, Molsheim, France). All glassware was cleaned with freshly prepared aqua regia and rinsed thoroughly with DI water before use. The western blotting kit was purchased from Sangon Biotech (Shanghai) Co., Ltd. Map2, IGF-1, NGL-1, βIII-tubulin, BrdU, Ibal, NLRP3, p-Tau, Aβ_1-42_, and GAPDH primary antibodies were purchased from Thermo Fisher Scientific. GAP43 primary antibody was purchased from Santa Cruz Biotechnology. The secondary antibodies (goat anti-mouse IgG [H+L] Highly Cross-Adsorbed Secondary Antibody, Alexa Fluor™ Plus 555, and goat anti-rabbit IgG [H+L] Cross-Adsorbed Secondary Antibody, Alexa Fluor™ 488) were purchased from Thermo Fisher Scientific. ELISA kits (GABA, 5-HT, dopamine, glutamate, p-Tau, Aβ_1-42_) were purchased from Jingkang bio, Shanghai. 4',6-diamidino-2-phenylindole (DAPI) was purchased from Beyotime Biotechnology, Haimen, Jiangsu. Cell culture medium and related cytokines were purchased from Thermo Fisher Scientific.

**2. Spectroscopic Characterization**

Circular dichroism (CD) spectra were obtained with a Chirascan CD spectrometer from Applied Photophysics Limited. Confocal imaging was performed with a Leica TCS SP8 confocal fluorescence microscope. Photoacoustic (PA) imaging was performed with a MSOT inSight/inVision 256 imaging system (iThera Medical, Munich, Germany). Polymerase chain reaction (PCR) was run on an Applied Biosystems 7900HT Fast Real-Time PCR System (Waltham, MA USA). The flow-cytometric analysis was performed on a BD FACSAria™. Raman scattering spectra were measured by a LabRam-HR800 Micro-Raman spectrometer with LabSpec 5.0 software. TEM images were obtained with a JEOL JEM-2100 transmission electron microscope at an acceleration voltage of 200 kV. The particle size distributions and zeta potential were measured with the Zetasizer Nano ZS system (Malvern Panalytical, Ltd.). Brain tissue fluorescence imaging was acquired with an *in* *vivo* imaging instrument (IVIS Lumina XRMS Series III). Inductively coupled plasma mass spectrometry (ICP-MS) results were obtained using an ICAP TQ (Thermo Fisher Scientific Inc., Germany). X-ray diffraction (XRD) patterns were obtained on a Bruker D8 Advance system (Bruker, Germany). X-ray photoelectron spectroscopy (XPS) spectra were obtained with an AXIS Supra by Kratos Analytical Inc.

**3. Experimental methods**

**3.1 Synthesis of chiral Ni(OH)_2_ Nanoparticles (NPs)**

Chiral Ni(OH)_2_ nanoparticles (NPs) were synthesized using an amino acid mediated one-step synthesis method. Briefly, 238 mg of nickel II chloride (NiCl_2_) and 750 mg of glutathione (*D*-GSH, *L*-GSH, or *rac*-GSH) were mixed and dissolved in 60 mL of ultrapure water. After mixing evenly, 5 mL of NaOH (1 mol/L, 1 M) were added and stirring was continued for more than 12 h until the color of the solution turned to light green (blue). The chiral Ni(OH)_2_ NPs were then washed with isopropanol three times and resuspended in ultrapure water for use.

**3.2 Computational Models**

All energy minimization and MD simulations were performed using the CUDA-enabled Gromacs 2023.1 package (*1-3*). Simulations were carried out using a cutoff of 1.2 nm for both Lennard-Jones and Coulomb potentials. For the former, a force-switch was applied from 1.0 nm to 1.2 nm, while the truncation errors for the latter were corrected using the Particle-Mesh Ewald (PME) algorithm (*4, 5*). The LINCS algorithm (*6*) was used to keep all bonds containing hydrogen atoms at their equilibrium geometry, allowing the use of an integration time step of 2 fs. Interaction energies were calculated by post-processing the trajectories using energy groups. Because the electrostatic energy decomposition is not possible when using PME, these contributions were recomputed using the reaction-field-zero method along with a dielectric value of 78.5 beyond the cutoff.

The VMD 1.9.3 software (*7*) was used to render structures and to calculate hydrogen bond occupancies.

**3.2.1 MD Simulations for the growth of Ni(OH)_2_ Nanoclusters**

The growth of nanoparticles takes place in macroscopic timescales and involve very large numbers of atoms, rendering the computer simulations of these processes unfeasible. Nonetheless, smaller model systems can be used to track the formation of small clusters, providing insight about the earlier stages of the growth of larger structures. The model system comprised 48 Ni(OH)_2_ units, 6 GSH molecules (thiolate form), 3600 water molecules, and 3 Ni²⁺ cations, which were needed to neutralize the GSH charges. All molecules were randomly placed in a cubic box with initial edge length of 5 nm. Energy minimization was performed using the steepest descent algorithm until forces below 500 kJ/mol/nm were reached. The growth process was monitored along 200 ns of MD simulations using the constant-NpT ensemble (T = 300 K, p = 1 bar). All the interactions were described using the Charmm36 force field (*8*), with atomic partial charges for Ni(OH)_2_ units described by the CM5 model, obtained using the GFN1-xTB Hamiltonian as implemented in the xTB 6.5.1 software (*9, 10*). These are the same parameters used for the MD simulations of the functionalized NPs, except for the Lennard-Jones parameters (s_ij_) for the Ni-O, Ni-S, and S-O atomic pairs, which were set to the distances between these atoms in the GSH-NP structure. With these modifications we were able to emulate the formation of “new bonds” (or very strong cohesion) between Ni(OH)_2_ units and also between Ni(OH)_2_ and GSH units. Simulations were performed for pure enantiomeric *D*-GSH or *L*-GSH systems and for the racemic one (3 *D*-GSH + 3 *L*-GSH), in addition to one system without ligands. The initial structure for the *L*-GSH system was obtained by mirroring all the Cartesian coordinates of the *D*-GSH model after the energy minimization step.

**3.2.2 GSH-Nanoparticle parameterization**

The MD simulations of NPs functionalized with glutathione (GSH) required the parameterization of a force field, since they typically lack proper parameters for ceramics. Firstly, the structure of our model Ni(OH)_2_ NP was drawn using the VESTA 3.4.4 software (*11*), by the replication of the unit cell (Crystallography Open Database (*12, 13*), 1548811 Identifier), cutting the supercell along the relevant crystallographic planes identified in TEM analyses to yield a nearly spherical NP containing 1285 atoms (257 Ni, 514 O, and 514 H).

Using a neighbor-search algorithm, the NP had 42 hydroxyl groups, which is likely to be a metastable configuration. These groups were deleted yielding a NP with a net charge of +42 *e*, which corresponded to the structure in which the thiolate ending of the GSH ligands should bind. A single-point energy calculation with the GFN1-xTB semiempirical Hamiltonian (*9*) – as implemented in the xtb 6.5.1 software (*10*) – was performed for this cationic NP to assign the atomic partial charges using the CM5 protocol. The vacant positions of these deleted oxygen atoms were then used as the initial positions of the terminal S atoms of the GSH ligands (in the thiolate form). Since each one of the 42 ligands presented a –1 *e* charge, the functionalization yielded a neutral system. Two model systems were built according to the nature of the ligands: i) one named *L*-NP bearing 42 *L*-GSH ligands and ii) another named *rac*-NP bearing 21 *L*-GSH and 21 *D*-GSH ligands, added in alternating fashion (**Figure S15**). Although our computational models were smaller than the experimental NPs, with surface areas of 33 nm² (model) and 113 nm² (experimental); the ligand surface density was 1.3 GSH/nm² in both cases.

The ligands were added with random orientations with respect to the NP and to each other, which required an initial relaxation of their structures while keeping the NP atoms frozen at their crystallographic positions. This initial relaxation was performed in vacuum as follows: i) energy minimization using the steepest descent algorithm; followed by ii) another energy minimization step using the conjugated-gradient algorithm; followed by iii) a 250 ps molecular dynamics simulation along a temperature-annealing ramp from 1 K to 2000 K using the stochastic-dynamics integrator with a 0.8 fs timestep; and followed by another iv) 250 ps temperature-annealing ramp from 2000 K to 1 K.

Because of the size of the system, the large number of ligands, and the intrinsic complexity of the NP structure, a full geometry optimization using quantum chemistry methods (even within the GFN1-xTB semiempirical level) was prohibitive due to the number of optimization cycles and/or molecular dynamics steps required to relax the whole structure. Thus, all parameterization steps were performed within the classical force field approximation, as follows: i) Lennard-Jones parameters for the NP atoms were taken from the July 2022 revision of the Charmm36 forcefield (*8*); ii) parameters for the thiolate-GSH were taken from the SwissParam server (*14*); and iv) missing parameters were estimated and refined self-consistently. The set of interatomic distances between the Ni, S, O, and H atoms of the GSH-NP was used to construct a neighbor list of pairs within a given distance, which were converted to bond interactions, as detailed in **Table S6**.

Using these initial parameters set, another equilibration round was performed, as follows: i) a full energy minimization using the steepest descent algorithm; followed by ii) another full energy minimization using the conjugated-gradient algorithm; followed by iii) a 5 ns molecular dynamics simulations temperature-annealing ramp from 1 to 300 K using the stochastic dynamics integrator with a 0.8 fs timestep; and followed by 295 ns stochastic dynamics with a 0.8 fs timestep at T = 300 K. The resulting NPs presented a significant number of chiral distorsions induced by the surface ligands. These distortions were mapped into the force field by adding extra bonding interactions between Ni atoms (**Table S7**), with different torsions observed for the *L*-NPs and the *rac*-NPs. Finally, the *D*-NP structure was obtained from the optimized *L*-NP structure by mirroring its coordinates.

**3.2.3 Protein model**

The full 640-residue structure of the leucine-rich repeat-containing protein 4C (LRC4C) was taken from the UniProt database (**Figure S16a**). This structure contains regions with low-confidence predicted results and/or no structural consensus, including the transmembrane domain (M523 to Y548, highlighted in purple) and the final 92-residues corresponding to the cytoplasmic region of the protein (highlighted in cyan) (*15*). Since the interaction with the NPs is expected to occur at the LRR region (LRR1 to LRR9, highlighted in orange and red), we have removed the low confidence regions, considering only the 426 residue fragment beginning at P25 and ending at P450 in our simulations (**Figure S16b**). Both 10-residue termini from this smaller structure (P25-Q34 and L441-P450) were restrained in all simulation steps to mimic the binding with the membrane, avoiding unphysical translations and rotations of the structure. The protonation state of the whole protein was attributed using the propKa 3.0 software (*16*), with a pH value of 7.0.

**3.2.4 Molecular dynamics (MD) simulations of GSH-NP –Protein interactions**

Three independent MD simulations were carried out for the *L*-NP, *D*-NP, and *rac*-NP interacting with the LRR portion of the protein to gather information regarding the enantioselectivity of these interactions. To avoid biased initial structures for the protein-NP complexes, a coarse-grid, rigid-body scan of translations and rotations of the NPs near the concave surface of the protein was performed using the development version of our lab-made software Themis (*17*), Software available at https://github.com/colombarifm/themis. We generated 30 translation points along the bisecting angle of the concave region of the protein, corresponding to 30 different distances along the reaction coordinate of approximation of the NPs with respect to the protein surface. In each translation point, we rotated the NPs using the two Euler angles defining the relative orientation between the protein and the NPs, which amounted to 42 x 120 = 5040 rotation moves, to yield a total of 150structures along the reaction coordinate. The interaction energy was computed for each protein-NP configuration using the Charmm36 forcefield. To decrease the calculation time, Themis skips configurations where interatomic distances fall below 0.15 nm (and a repulsive energy value is considered for them) and a cutoff radius of 1.2 nm were considered. The most probable structures for the *L*-NP, the *D*-NP, and the *rac*-NP complexes (**Figure S17**) were used as the starting structures for MD simulations as follows: i) the protein-NP complex was centered in a 14.3 nm x 14.4 nm x 9.5 nm periodic box; ii) a total of 60,000 TIP3P water molecules were added; and iii) four water molecules were randomly replaced by chloride ions to neutralize the system charge.

An energy minimization step with the steepest descent algorithm was carried out to remove repulsive contacts in the initial structure and resulting coordinates were equilibrated for 0.5 ns in the canonical ensemble (NVT) (T = 300 K, v-rescale thermostat) with a position restraint potential for all protein heavy atoms. Another 0.5 ns NVT simulation was carried out with the position restraint potential applied only on the first and last 10 residues of the protein to mimic the protein embedding in the biological membrane (this restriction will be applied for all simulations hereafter). After this step, the system was further equilibrated for another 50 ns in the isobaric-isothermal ensemble (NpT) (T = 300 K, v-rescale thermostat; p = 1 bar, Berendsen barostat) (*18*) to adjust the density. The data collection runs corresponded to 1000 ns simulations in the NpT ensemble (T = 300 K, v-rescale thermostat; p = 1 bar, Parrinello-Rahman barostat) (*19*).

**3.2.5 Quantum chemistry calculations of GSH-NP – Protein complexes**

Due to the large size of the systems, Quantum-Mechanical (QM) calculations – even at the semiempirical GFN2-xTB – are challenging. Thus, subset structures were cut out from the MD simulations for each enantiomer, each one contained the full GSH-NP, the protein fragment in closer contact with the nanomaterial (residues T128 to N331), and the water molecules within 0.3 nm of any atoms comprising the NP-protein interface. We considered the MD simulation frame at 900 ns since it presented the most favorable interaction energy between the *D*-GSH-NP and the protein. The backbone of each protein fragment was saturated by adding -NH_2_ or -COOH termini, resulting in model systems containing 6120, 6198, and 6195 atoms for *L*-GSH-NP, *D*-GSH-NP, and *rac*-GSH-NP, respectively. The electronic structure calculations were performed using the xtb4stda 1.0 software (*20*), with the GFN2-xTB (*21*) semiempirical Hamiltonian and the GBSA implicit solvation model for water. The ECD spectra were calculated using the sTDA 1.6 software (*20*), with an energy window of 4 eV. For comparison purposes, similar calculations were performed considering only the protein fragment from the *D*-GSH NP system (**Table S2**). For each system, Natural Transition Orbitals (NTOs) corresponding to the 10 lowest energy transition were calculated for further analysis.

**3.3 Cell lines and incubation conditions**

Primary hippocampal neurons were extracted from 10-month-old wild-type (WT) or Alzheimer’s disease AD mice (*22*). The mice were anesthetized by a procedure performed in accordance with ethical principals in a container with pentobarbital, and then the head was disinfected with 70% ethanol. The skin over the top of the skull was dissected to expose the skull, and the calvarium on both sides near to the front were carefully cut to avoid damage to the brain. The brain was exposed, gently extracted, and transferred to a dissection dish. The hippocampus regions were rapidly dissected, transferred to filter paper, and then cut into 0.5 mm slices. The extracted slices were digested with papain for 40 min in 15 mL tubes at 37 °C, and the cell suspensions were centrifuged for 2 min at 1,100 rpm. The supernatant was discarded, and the cells re-suspended in neurobasal A/B27 medium containing glutamax, growth factors, and gentamycin and placed in a cell incubator.

Primary mouse dorsal root ganglion (DRG) neurons were extracted from 4-6-week-old female mice (*23, 24*). Briefly, the surrounding muscle tissue was removed with ophthalmic scissors, and the spinal canal was cut with microscope scissors. The spinal cord was exposed, and then L4–L7 DRG pairs were dissected under the microscope using microscope tweezers. The extracted DRG pairs were digested by digestive enzymes for 1 h in an oven at 37 °C. Following digestion, the digestive enzymes were removed, and culture solution was added. Finally, the DRG neurons were cultured in neurobasal culture medium containing B27 supplement and *L*-glutamine at 37 °C and 5% CO_2_.

**3.4 Cell viability**

Primary hippocampal neurons (1.0 × 10^6^) were seeded in a 96-well plate, with three wells for each concentration of the chiral *D*-NPs (2.5 mg/mL, 1 mg/mL, 0.5 mg/mL, 0.25 mg/mL, 0.1 mg/mL, 50 μg/mL, 25 μg/mL, 5 μg/mL, 2.5 μg/mL, 0.5 μg/mL, and 0.25 μg/mL) and nickel chloride solution with the same concentration Ni. The cells were incubated at 37°C for 48 h in culture medium, which was then replaced with 100 μL of fresh Opti-MEM (Life Technologies). The cells were treated with 10 μL of Cell Counting Kit-8 (CCK-8, Beyotime) for 2 h. The absorbance (A) of each well was measured with a microplate reader at 450 nm, and the relative cell viability (%) was calculated as (A_test_/A_control_) × 100. The control group represented the cells without Ni treatment.

**3.5 NIR-induced chiral NPs promote axon regeneration**

Primary hippocampal neurons and DRG neurons were seeded on a culture well at a density of 1 × 10^6^ cells for 24 h. After the cells had adhered to the substrate, *D*-, *L*-, or *rac*-NPs were added at a final concentration of 100 ng/mL, and then illuminated with near-infrared radiation (NIR, 980 nm laser; 400 mw/cm^2^) for 10 min every 2 h, up to 12 h. Incubation continued for a further 36 h, and the neurons were then supplemented with fresh medium containing 100 ng/mL *D*-, *L*-, or *rac*-NPs and illuminated with NIR. The neurons were replenished with fresh medium containing 100 ng/mL NPs every two days under NIR illumination six times at intervals (10 min every 2 h), and this process continued for six days. On the seventh day, the cells were replenished with fresh medium containing 100 ng/mL NPs and under NIR illumination (980 nm laser; 400 mw/cm^2^) for 10 min every 2 h, up to 12 h. Incubation continued for another 12 h, and finally used for other experiments.

**3.6 Cell immunofluorescence staining**

The treated cells were washed three times with phosphate-buffered saline containing Tween 20 (PBST) and fixed with 4% paraformaldehyde for 20 min. The fixed cells were washed three times with PBST and then permeabilized with 0.2% Triton X-100 for 5 min. After washing three times with PBST, 1 mL of 1.5% (v/v) bovine serum albumin (BSA) in PBST was added for 2 h to block the cells. The cells were then washed three times with PBST, and incubated with primary antibody (Map2, GAP43, βIII-tubulin, IGF-1, NGL-1) for 2 h. After that, the cells were again washed three times with PBST (5 min each time), and then incubated with secondary antibodies (goat anti-mouse IgG [H+L] Highly Cross-Adsorbed Secondary Antibody, Alexa Fluor™ Plus 555, and goat anti-rabbit IgG [H+L] Cross-Adsorbed Secondary Antibody, Alexa Fluor™ 488) for 2 h. Finally, the cells were washed with PBST for 15 min and 4,6-diamidino-2-phenylindole (DAPI) was used to counterstain the nuclei for 10 min. The cells were observed with an inverted fluorescence microscope (Leica DMI8).

**3.7 Western blot analysis**

Hippocampal neurons (1.0 × 10^6^), DRG neurons (1.0 × 10^6^), brain tissue, and spinal cord tissue were collected for western blot analysis, and their proteins were extracted with RIPA Lysis Buffer IV (Beyotime). The protein lysates were separated by SDS-PAGE, and then transferred onto polyvinylidene difluoride (PVDF) membranes using the semi-dry method. Defatted milk powder (0.5 mg/mL) was added for 1.5 h to block the PVDF membranes. After washing three times with tris-buffered saline with 0.1% Tween 20 detergent (TBST) buffer, the membranes were incubated with primary antibodies directed against Map2, GAP43, βIII-tubulin, IGF-1, NGL-1, laminin, MBP, p-ERK, p-PI3K, and GAPDH for 2 h at room temperature. The membranes were washed three times with TBST for 10 min, and then incubated with horseradish-peroxidase-conjugated secondary antibody for 1 h. Lastly, the PVDF membranes were placed in the luminescence instrument to acquire images using the chemiluminescence chromogenic kit (Sangon Biotech, China).

**3.8 Flow cytometry**

To quantify FOXP3^+^ Treg cells expression and AREG levels in Treg cells at the SCI sites, SCI tissues were removed and ground to a cell suspension in PBS buffer. The cells were then incubated with target antibodies (CD4, FOXP3^+^, and AREG) for 40 min at room temperature in the dark. The cells were washed three times with PBS and resuspended in 500 μL of PBS for flow cytometry analysis.

To determine the distribution of chiral NPs in brain cells, chiral NPs carrying fluorescent molecule Cy5 were intravenously injected into AD mice. After 6 h, brain tissues were collected to obtain a cell suspension. The cells were fixed with 4% paraformaldehyde, permeabilized with 0.1% Triton X-100, blocked with 1.5% BSA, and then incubated with primary antibody (Map2, GFAP, Ibal) for 2 h. After washing three times with PBS, secondary fluorescent antibody (goat anti-mouse IgG [H+L] Highly Cross-Adsorbed Secondary Antibody, Alexa Fluor™ Plus 555 or goat anti-rabbit IgG [H+L] Cross-Adsorbed Secondary Antibody, Alexa Fluor™ 488) was added for 2 h. The flow cytometry data were analyzed with FlowJo 10.3 and GraphPad Prism software.

**3.9 Neurotransmitter detection**

The cell culture supernatant (Section 3.5) was collected to determine the expression of neurotransmitters, including GABA, 5-HT, dopamine, and glutamate with the ELISA kit (Jingkang bio, Shanghai), according to the manufacturer’s instructions. A reliable LC-MS/MS method was established for 5-HT detection (*25*).

**3.10 Global gene expression analysis**

Total RNA in each sample was extracted from hippocampal neurons and DRG neurons using total RNA extractor (Trizol, Sangon Biotech, China). Total RNA was accurately quantified with the Qubit RNA detection kit. The following steps were then performed in turn: mRNA purification and fragmentation, double-stranded cDNA synthesis, and double-stranded cDNA purification. Both ends of the purified double-stranded cDNA were repaired with End Prep Enzyme, and then a T-A ligation was added to both ends. Adaptor ligated DNA was then purified, and fragment sorting was performed. Each sample was then amplified by PCR and the products were purified using Hieff NGS™ DNA Selection Beads. All purified products were detected by gel electrophoresis, and the main fragment between 300-500 bp was sequenced. Finally, the Qubit DNA detection kit was used to accurately quantify the recovered DNA, to facilitate sequencing after equal mixing in a 1:1 ratio.

**3.11 Membrane potential detection in neurons**

Membrane potential analysis of hippocampal neurons was recorded by a whole cell current clamp (*26*), which was conducted using a patch clamp amplifier at room temperature of 20-25 °C. The experimental parameter settings, data acquisition, and stimulation mode application were controlled by Pulse software, through 4 kHz filtering and 20 kHz data sampling. The glass microelectrode was drawn by the PP-830 electrode drawing instrument in two steps. After filling the electrode with liquid, its impedance was 2~4 MΩ. When the cell membrane and the electrode reached GΩ sealing, the membrane broke to form the whole cell state, which was stable for 3-5 min, and then switched to the current clamp mode. The voltage signal was recorded in current clamp mode, and the membrane current was set to zero to record the resting membrane potential.

**3.12 Animal models**

AD models: Ten-month-old app/ps1/tau transgenic mice (3xTg-AD) express three mutant allele homozygotes, including Psen1 mutant homozygotes, APPSwe and tauP301L transgenic homozygotes (Tg [APPSwe, tauP301L] 1Lfa).

Spinal cord injury model: 4-6-week-old healthy female Kunming mice were anesthetized by intraperitoneal injection of 3.6% chloral hydrate and fixed on the operating table in the prone position. After disinfection, an incision was made in the middle of the back about 2 cm long, centered on the T10 spinous process. The vertebrae were exposed, and T10 lamina was removed to fully expose the spinal cord. The right T10 spinal cord was completely removed with iris scissors, or damaged by needle puncture, which led to paralysis of the ipsilateral hind limb, and no response to stimulation. The muscle and skin were then sutured in turn. The local activation in animals was achieved through the illumination externally without an implant. The model mice were intravenously injected with 40 μg of *D*-, *L*-, or *rac*-NPs once every five days for 60 days, and NIR (980 nm laser with the optical collimator to be a broad beam [21 cm, diameter], 600 mw/cm^2^) illumination for 12 h every day.

All animal experiments were conducted in accordance with the institutional ethical guidelines and the Committee on Animal Welfare of Jiangnan University (JN. No20240430t0800809[198]; JN. No20240315t0720928[094]).

**3.13 In *vivo* PA brain imaging**

PA brain imaging was performed with a multispectral optoacoustic tomography scanner (MSOT, iThera Medical). The 3xTg-AD mice were anesthetized with 1.5% isoflurane delivered via a nose cone, and the PA images were captured at different time points.

**3.14** **Brain tissue fluorescence imaging**

3xTg-AD mice were intravenously injected with *D*-NPs-Cy5, and then brain tissues were collected at different times. The images were acquired with an *in vivo* imaging instrument (IVIS Lumina XRMS Series III) under the excitation wavelength of 650 nm.

**3.15 Measurement of Aβ_1-42_ and p-Tau in CSF**

Detection of Aβ_1-42_ and p-Tau concentration in cerebrospinal fluid (CSF) was by ELISA (Jingkang bio, Shanghai). In brief, 96-well microplates were first coated with 100 μL of Aβ_1-42_-ovalbumin (OVA) and p-Tau-OVA and incubated at 37°C for 2 h. Then, each well was washed three times and blocked with 200 μL of blocking buffer for 2 h at 37°C. Next, 50 μL of serially diluted standard Aβ_1-42_, p-Tau solution, or CSF was added to different wells, and then 50 μL of anti-Aβ_1-42_ monoclonal antibody or anti-p-Tau monoclonal antibody was added to each well and incubated at 37 °C for 30 min. After washing three times, 100 μL of horseradish-peroxidase-labeled goat anti-mouse IgG was added to each well at 37 °C. Thirty minutes later, 100 μL of tetramethylbenzidine substrate was added and allowed to react for 15 min at 37 °C in the dark. Lastly, 50 μL of termination solution was added to each well and mixed thoroughly. The optical density (OD) at a wavelength of 450 nm (OD_450_) was measured with a microplate analyzer.

**3.16 Morris water maze (MWM) test**

To assess the cognitive abilities of mice, the Morris water maze (MWM) test was conducted. The diameter of the maze was approximately 1.2 m, and the water temperature in the maze was maintained at 21 ± 2 °C and drained daily. The mice were trained before the formal experiment, during which the starting positions were balanced and the platform with a diameter of 10 cm was fixed 1 cm below the water surface. Within a 120 s latency period, the mice were placed at different positions around the border of the maze in a semi-random order and allowed to reach the platform. This test was performed once a day for 5 days. If the mice found the platform successfully within 120 s, they were allowed to stay on the platform for 15 s. If they failed, they were placed on the platform for 15 s. After 24 h of the final learning trial, their memory retention was evaluated. The trained mice were placed at two starting points far from the platform and allowed to swim freely for 120 s. The motion tracks were recorded and analyzed with a computerized video tracking system. In addition, the time they spent looking for the platform was recorded.

**3.17 Nissl staining**

The whole brains of each mouse were harvested, fixed with 10% formalin, embedded in paraffin, and sections of 3 µm were prepared. The paraffin sections were then deparaffinized with xylene, rehydrated with ethanol, and stained with Nissl staining solution. After rinsing with water, the sections were dehydrated with 100% ethanol and xylene and finally blocked with neutral gum. Observations were conducted under an inverted microscope to detect Nissl bodies in the cytoplasm of hippocampal neurons.

**3.18** **Immunofluorescent and** **immunohistochemical staining**

For immunofluorescent (IF) staining, the mice brain and spinal cord tissue sections were deparaffinized with xylene and rehydrated with ethanol. Following rinsing three times with PBST, the sections were blocked with 1.5% BSA for 1 h and then washed three times with PBST. The primary antibodies (Map2, Brdu, Ibal, NLRP3, GFAP, laminin, IGF-1, Aβ, p-Tau) were incubated with the sections at room temperature for 2 h or at 4 °C overnight. The sections were washed three times with PBST (5 min each time), and then incubated with the secondary antibodies from Thermo Fisher (goat anti-mouse IgG [H+L] Highly Cross-Adsorbed Secondary Antibody, Alexa Fluor™ Plus 555, and goat anti-rabbit IgG [H+L] Cross-Adsorbed Secondary Antibody, Alexa Fluor™ 488) for 2 h. After washing three times, the sections were incubated with DAPI in the dark for 15 min to stain the nuclei. An inverted fluorescence microscope (Leica DMI8; Germany) was used for IF observations and image processing. Firstly, the prepared samples were under a microscope for observation and placed in the area to be photographed in the center of the field of view. Then in “live” mode, with the laser intensity of 10, “Pinhole” value of 1 AU, and “Gain” value of 800. After adjusting the focal length, the imaging was captured at the speed of 10 s and averaging scan number of two. Finally, the imaging was saved for further analysis.

For immunohistochemical (IHC) staining, paraffin sections after treatment with bio-dewaxing solution, ethanol, and 3% hydrogen peroxide were blocked with 1.5% BSA for 1 h. Then, anti-Tau Recombinant Polyclonal Antibody (clone 6HCLC, 710080; Thermo Fisher Scientific) or beta Amyloid Polyclonal Antibody (clone CT695, 51-2700; Thermo Fisher Scientific) was incubated with the sections at room temperature for 2 h or at 4 °C overnight. After washing with PBST, horseradish peroxidase (HRP)-labeled secondary antibody was added to the sections. After 2 h, diaminobenzidine (DAB) staining solution and hematoxylin staining solution were added. Photographs were then obtained under the microscope.

**3.19** **Evaluation** **of hind limb locomotor ability**

To analyze their footprints, the hind limbs of SCI mice were immersed in blue dye (*27*). A narrow track (100 cm long and 5 cm wide) was lined with white paper and the mice were allowed to pass. The distance from the start to the end of a step using the back paw was defined as stride length. The distance from the left outermost toe to the right outermost toe was defined as stride width. The footprints images were acquired, and the average stride length and width were measured.

**3.20 In *vivo*** **metabolism and toxicity assessment of chiral NPs**

We chose C57BL/6 mice to assess the metabolism and toxicity of chiral NPs, and the mice were divided into four groups for the metabolism experiment (at least three animals per group). One group was the control, and the other groups were intravenously injected with *D*-, *L*-, or *rac*-NPs (2 mg/kg, corresponding to a Ni dose of 0.1 mg/kg). The amount of Ni in the brain, blood, liver, and kidney was quantified by ICP-MS. For toxicity assessment, different groups of mice were intravenously injected with *D*-, *L*-, *rac*-NPs, or NiCl_2_ at the dose of 100 μg Ni/kg, once every 5 days up to 60 days. Serum samples were collected to detect hepatorenal toxicity related indicators, including alanine aminotransferase (ALT), aspartate aminotransferase (AST), blood urine nitrogen (BUN), and creatinine (CRE). The main organs were harvested and fixed with 4% paraformaldehyde, and then embedded in paraffin, sliced, and stained with hematoxylin and eosin (H&E).

For embryotoxic and teratogenic effects, the animals were divided into five groups. One group was the control, and the other groups were intravenously injected with *D*-, *L*-, *rac*-NPs (2000 μg Ni/kg), or NiCl_2_ (400 μg Ni/kg) from day 6 to 13 of gestation, that is, the organogenetic period. The mice were sacrificed by cervical dislocation on day 18 of gestation, and the uteri of all sacrificed mice were examined for morphological alterations. The fetuses of the control, *D*-NPs, and NiCl_2_ groups were fixed in 95% alcohol for double staining (alizarin red S and Alcian blue) to observe skeletal anomalies.

**3.21 Quantitative real-time PCR**

Total RNA, extracted from cell or tissues samples, was prepared using the UNIQ-10 column total RNA extraction kit (Sangon Biotech, China) according to the manufacturer’s instructions. The concentration of extracted total RNA was quantified with a NanoDrop spectrophotometer. The RNA was then reverse-transcribed into cDNA with the MightyScript First Strand cDNA Synthesis Master Mix (Sangon Biotech, China). The 2×SG Fast qPCR Master Mix (High Rox) kit (Sangon Biotech, China) was used to quantify mRNA expression. Finally, 20 μL of qPCR samples were detected according to the following thermal cycling program: 95 °C for 3 min; 40 cycles of 95 °C for 3 s; and 60 °C for 30 s. After the reaction, the amplicons were quantified and a melting analysis was performed with the Applied Biosystems 7900HT Fast Real-Time PCR System (Waltham, MA, USA), according to the instructions. All the qPCR primers are shown in **Table S8**.

**3.22 Calculating the binding affinity between proteins and NPs**

An ATA Instruments Nano Isothermal Titration Calorimetry (ITC) Low Volume isothermal titration calorimeter was used to obtain binding affinities. The experimental process was as follows: *D*-, *L*-, or *rac*-NPs and NGL-1 were placed under vacuum for 20 min to remove all the bubbles; 350 μL of *D*-, *L*-, or *rac*-NPs solution (10 mM) was injected into the sample cell; and then 50 μL of NGL-1 protein (0.1 mM) was injected into the NPs solution using a computer-controlled microsyringe, with 2 μL per injection (25 injections in total) and an injection interval of 300 s. The temperature was set at 25°C. The stirring rate was 300 rpm. NanoAnalyze software was used to model with a constant blank model and an independent model.

**3.23 Interaction force measurement between cells and NPs**

The force of the interactions between *D*-/*L*-/*rac*-NPs and neurons was measured by AFM (*28, 29*). Firstly, the surface of the silicon nitride atomic force microscope probes was modified with *D*-, *L*-, or *rac*-NPs, and then the interaction force between the probe and the living neurons was directly measured. Specifically, neurons were incubated on the glass substrate, and the atomic force microscope probe moved downward to approach the surface of the glass substrate at a rate of 2 μm s^-1^. After contacting the substrate surface, downward pressure was continued until the pressure reached the set value (500 pN) so that the chiral NPs modified on the probe surface could fully react and combine with the cell surface. The atomic force microscope probe was then moved up 1.5 μm at the same rate. In this way, a series of force-stretch curves were obtained by pressing and stretching different positions on the cell surface to obtain the force curve of chiral NPs and cells.

**3.24 Nuclear Magnetic Resonance (NMR)**

The excess ligand in the original chiral NPs aqueous solution was removed by centrifugation at 16200 g for 10 min and the *D*-NPs were then resuspend in H_2_O at a concentration of 20 mM. The NGL-1 protein was dissolved in H_2_O at a concentration of 0.2 mM. Then 0.5 mL of the *D*-NPs and 0.5 mL of the NGL-1 were respectively mixed with 0.5 mL of H_2_O to obtain final concentrations of 10 mM and 0.1 mM. 0.5 mL of the 20 mM *D*-NPs and 0.5 mL of 0.2 mM NGL-1 were mixed at a concentration ratio of 100:1 to acquire *D*-NPs- NGL-1 complex. Taking 0.5 mL of *D*-NPs, NGL-1, and *D*-NPs-NGL-1 complex solution for NIR illumination (980 nm, 400 mW/cm^2^, 10 min). Finally, 100 μL of *D*-NPs, *D*-NPs + NIR, NGL-1, NGL-1+NIR, *D*-NPs+NGL-1, and *D*-NPs+NGL-1+NIR samples were respectively mixed with 490 μL of D_2_O and transferred to NMR tubes. We used a WET1D pulse sequence with shaped selective pulses to suppress the remaining water signal to produce the ^1^H NMR spectra of D_2_O. Each spectrum was composed of 32 frequency domain scans with a spectral width of 16025.6 Hz, and an acquisition period of 2 s. NMR spectra were preprocessed with Bruker TopSpin 3.6.5 software.

**3.25 Human samples**

Serum and cerebrospinal fluid (CSF) samples were donated from identified AD patients (serum n = 70, CSF n = 53) and healthy humans (Serum n=56, CSF n=40) aged 50-90 years. The study was approved by the Institutional Review Board of Beijing Tiantan Hospital of Capital Medical University (KY-2021-028-01), and informed consent was obtained from all participants. All participants underwent a screening process that included a medical history, and physical and neurological examinations, and had no history of a significant psychiatric condition or traumatic brain injury that could compromise brain function. Based on this assessment, we excluded samples from patients with the following conditions: those taking antibiotics, probiotics, prebiotics, symbiotics, and other drugs that affect the gut microbiota within 3 months of sampling; those with severe heart, brain, liver, kidney, or hematopoietic system diseases or other serious primary diseases; and those with severe malnutrition, drug addiction, or alcohol abuse.

**3.26 Data analysis**

All statistical analyses were performed with GraphPad Prism version 8.0, using either two-tailed unpaired t-test to compare two groups or a one-way or two-way analysis of variance (ANOVA) to compare more than two groups, followed by Tukey’s multiple comparisons test. Each ‘n’ represents an independent biological sample. Values represent means ± s.d. A value of P < 0.05 was considered to indicate statistical significance (*P < 0.05, **P < 0.01 and ***P < 0.001).

For cell sample imaging, each included 500-1000 cells per well for independent observations and randomly selected 10-50 cells from any region for imaging and fluorescence statistics. Each experiment was repeated at least three times.

For axonal length of neuron statistics, 10-50 cells from any region of each well were chosen for independent observations and to calculate the average length. Each experiment was repeated five times.

For the imaging of tissue samples, the sections were collected for staining and imaging, at least five animals per condition.

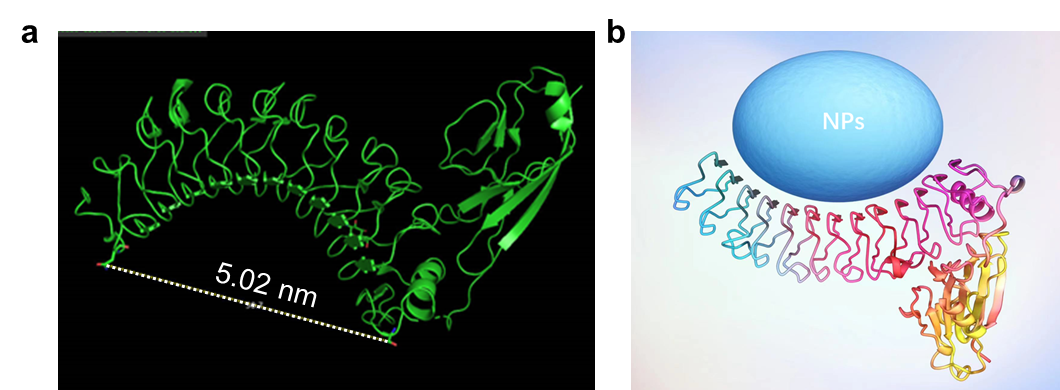

**Figure S1. (a)** 3D structure of NGL-1 fragment. **(b)** The schematic of the interaction between NGL-1 and NPs.

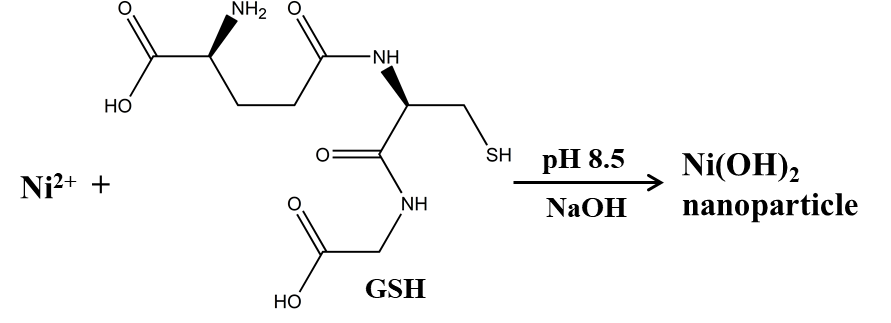

**Figure S2.** Schematic illustration of the formation of chiral NPs.

| 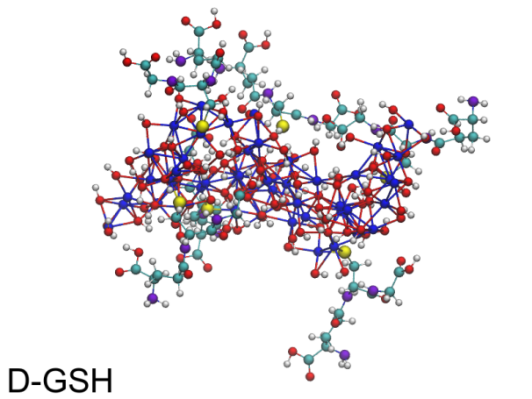 | 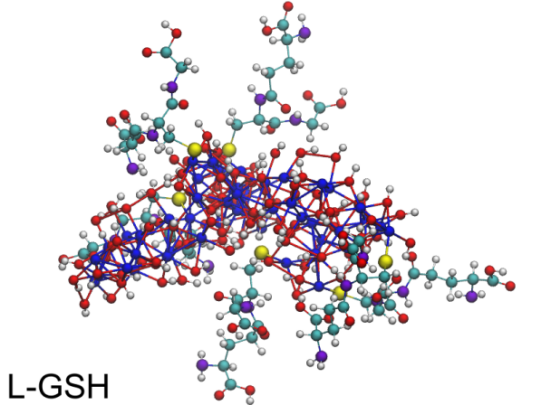 |
| --- | --- |
| 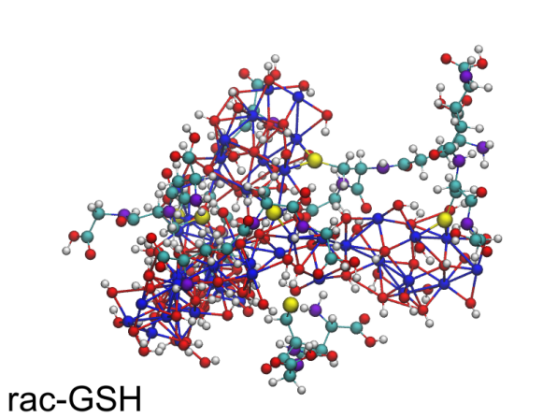 | 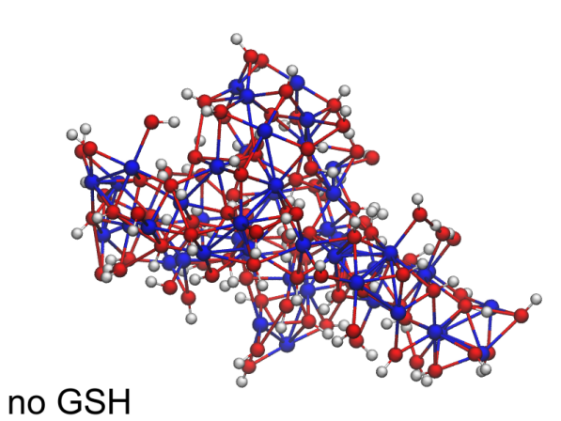 |

**Figure S3.** Final structures of the Ni(OH)_2_ clusters after 200 ns of MD simulation. Nickel atoms are presented in blue, oxygen atoms are presented in red, hydrogen atoms are presented in white, sulfur atoms are presented in yellow, carbon atoms are presented in cyan, and nitrogen atoms are presented in violet.

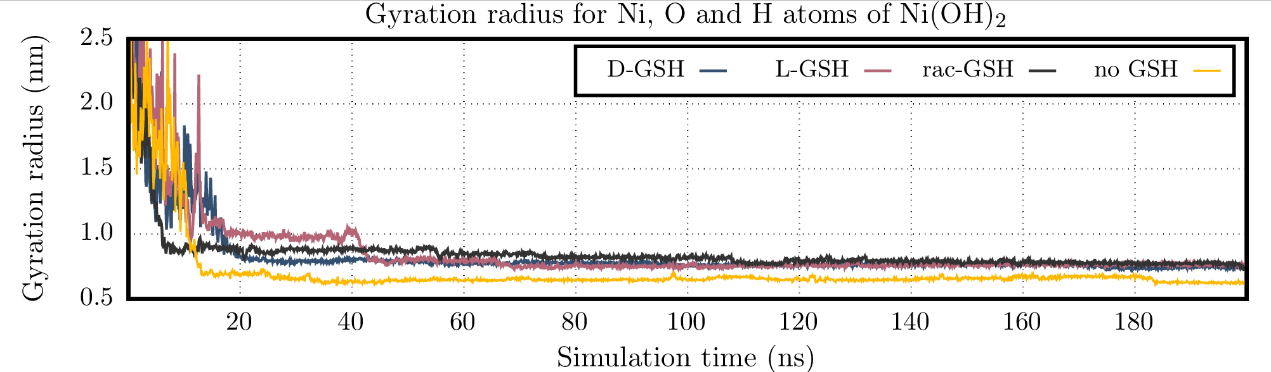

**Figure S4.** Gyration radii for Ni, O, and H atoms of Ni(OH)_2_ units obtained along each MD trajectory for the Ni(OH)_2_ NP growth.

**Comments:** The growth of nanoparticles takes place in macroscopic timescales and involves very large numbers of atoms, rendering the computer simulations of these processes unfeasible. Nonetheless, smaller model systems can be used to track the formation of small clusters, providing insight into the earlier stages of the growth of larger structures. The model system comprised 48 Ni(OH)_2_ units, 6 GSH molecules (thiolate form), 3600 water molecules, and 3 Ni²⁺ cations to neutralize the GSH charges. All molecules were randomly placed in a cubic box with an initial edge length of 5 nm. Energy minimization was performed using the steepest descent algorithm until forces below 500 kJ/mol/nm were reached. The growth process was monitored along 200 ns of MD simulations using the constant-NpT ensemble (T = 300 K, p = 1 bar).

All the interactions were described using the Charmm36 force field, with atomic partial charges for Ni(OH)_2_ units described by the CM5 model, obtained using the GFN1-xTB Hamiltonian. These are the same parameters used for the MD simulations of the functionalized NPs, except for the σ_ij_ Lennard-Jones parameters for the Ni-O, Ni-S, and S-O atomic pairs, which were set to the distances values between these atoms in the GSH-NP structure. With these modifications we were able to emulate the formation of “new bonds” (or very strong cohesion) between Ni(OH)_2_ units and also between Ni(OH)_2_ and GSH units. Simulations were performed for pure enantiomeric *D*-GSH or *L*-GSH systems and for the racemic one (3 *D*-GSH + 3 *L*-GSH), in addition to one system without ligands. The initial structure for the *L*-GSH system was obtained by mirroring all the Cartesian coordinates of the *D*-GSH model after the energy minimization step.

The aggregation of Ni(OH)_2_ units was monitored by the radius of gyration of these atoms along the MD simulations. Since these units (and the other system components) were randomly placed in the simulation box, the radius of gyration values was very large at the beginning of the simulations, dropping steeply during the first 10 ns and relaxing more slowly afterwards (**Figure S4**). Although some relaxation still occurred up to 200 ns (and possibly after that if trajectories were extended to longer timescales), visual inspection showed that all systems had a single cluster comprising all Ni(OH)_2_ units and ligands after 60 ns, so small changes and fluctuations after this point should be ascribed to structural reorganization of the cluster and not to further aggregation. So, all structural and thermodynamic analyses were performed for the time window between 60 and 200 ns. The final structures corresponded to distorted Ni(OH)_2_ lamellae with the GSH molecules (if present) attached to the border of the lamellae by the sulfur atom (**Figure S3**).

This information indicated that the growth mechanism for the early stages of NP formation was driven by the highly favorable interactions between Ni and O atoms of different Ni(OH)_2_ units, which was larger than their interaction with GSH molecules, whose S atoms interacted with exposed Ni atoms on the surface of the growing Ni(OH)_2_ cluster.

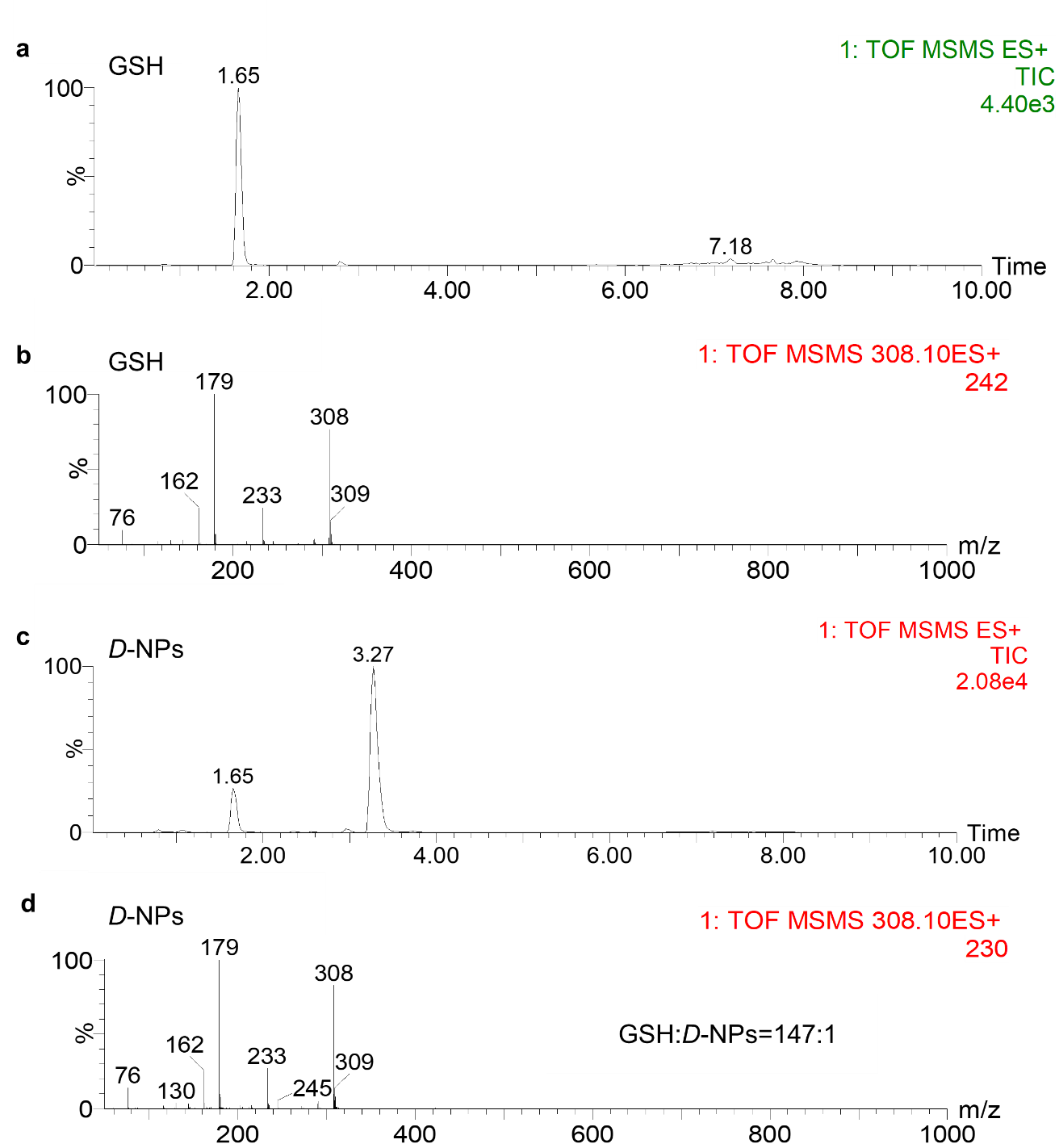

**Figure S5.** Calculated number of GSH ligands on chiral *D*-NPs using liquid chromatograph mass spectrometry (LC-MS). **(a, b)** LC-MS spectrum of GSH. **(c, d)** LC-MS spectrum of chiral *D*-NPs.

**Comments:** The density of Ni(OH)_2_ is assumed to be the same as in the bulk. Therefore, the number of GSH ligands on NP surfaces can be calculated to be 147. Specifically, m_GSH_ is 12 mg; m_D-NPs_ is 9.8 mg; ρ_Ni(OH)2_ is 4.15 g/cm^3^; Avogadro’s number *N_A_* = 6.02×10^23^ mol^-1^; M_GSH_ = 307.33g/mol; number of ligands = (m_GSH_×NA×ρ_Ni(OH)2_×V)/(M_GSH_×m_D-NPs_) ≈ 147.

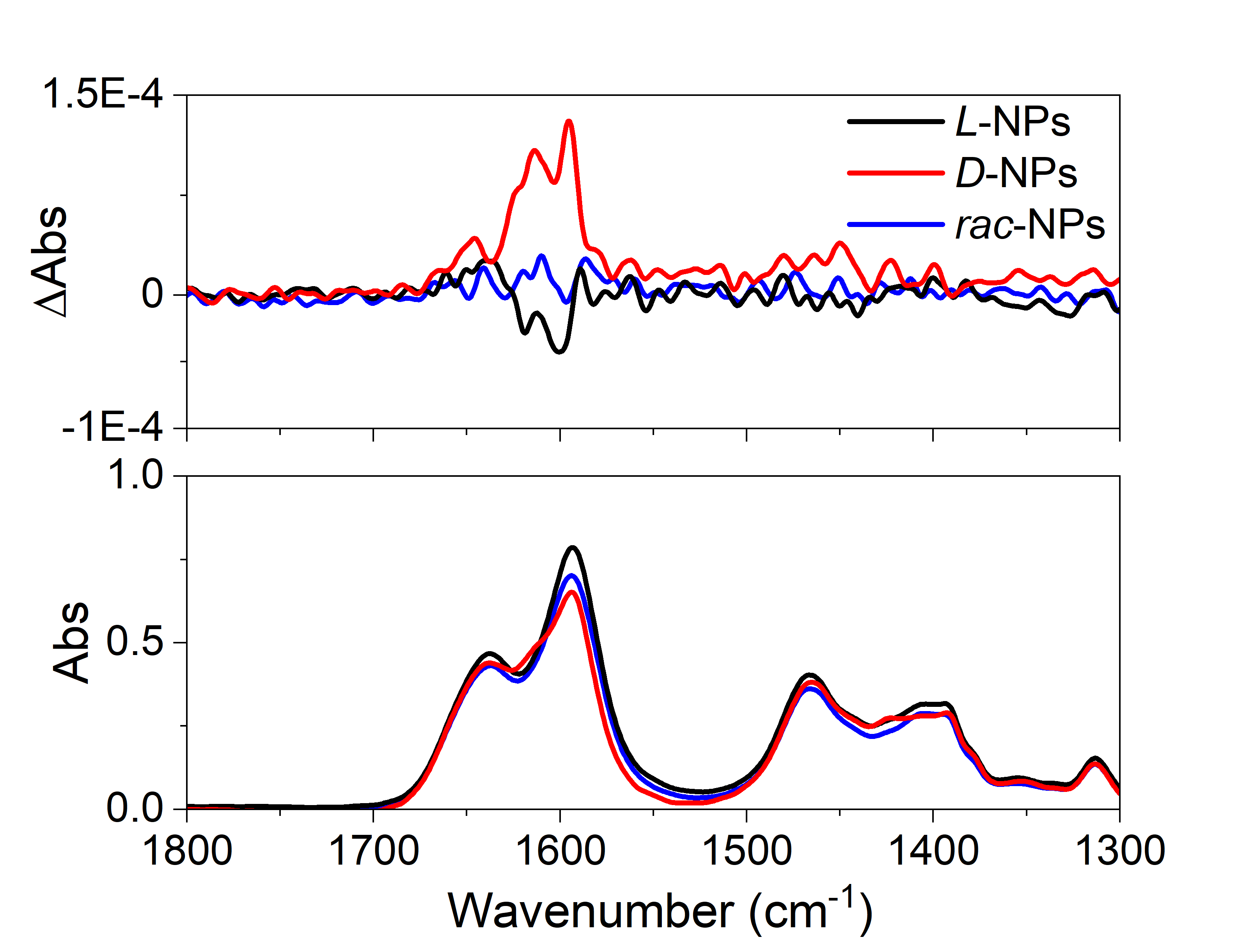

**Figure S6**. The Vibrational Circular Dichroism (VCD) Spectroscopy of chiral *L*-NPs, *D*-NPs, and *rac*-NPs.

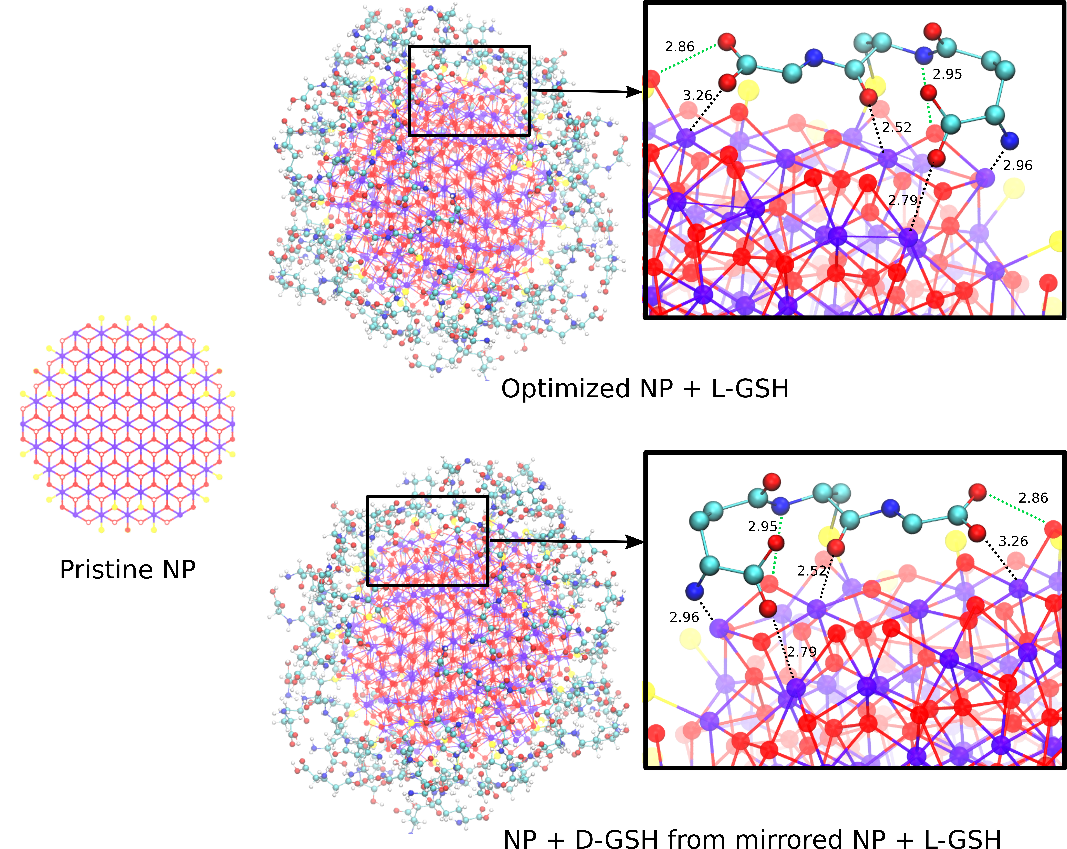

**Figure S7.** Pristine Ni(OH)_2_ NP with the experimental crystallographic structure (left) and the optimized *L*-GSH NP (top right) and its mirrored structure, the *D*-GSH NP (bottom right). Insets depict one GSH molecule and the interactions between its O and N atoms and the Ni and O atoms on the surface of the NP (distances in Å are indicated close to the dotted lines for each interaction, black lines correspond to direct Ni-O and Ni-N coordination, whereas green lines correspond to hydrogen bonds between GSH and OH groups from Ni(OH)_2_). Ni atoms are depicted in violet, O in red, N in blue, and S in yellow (H atoms have been omitted for clarity).

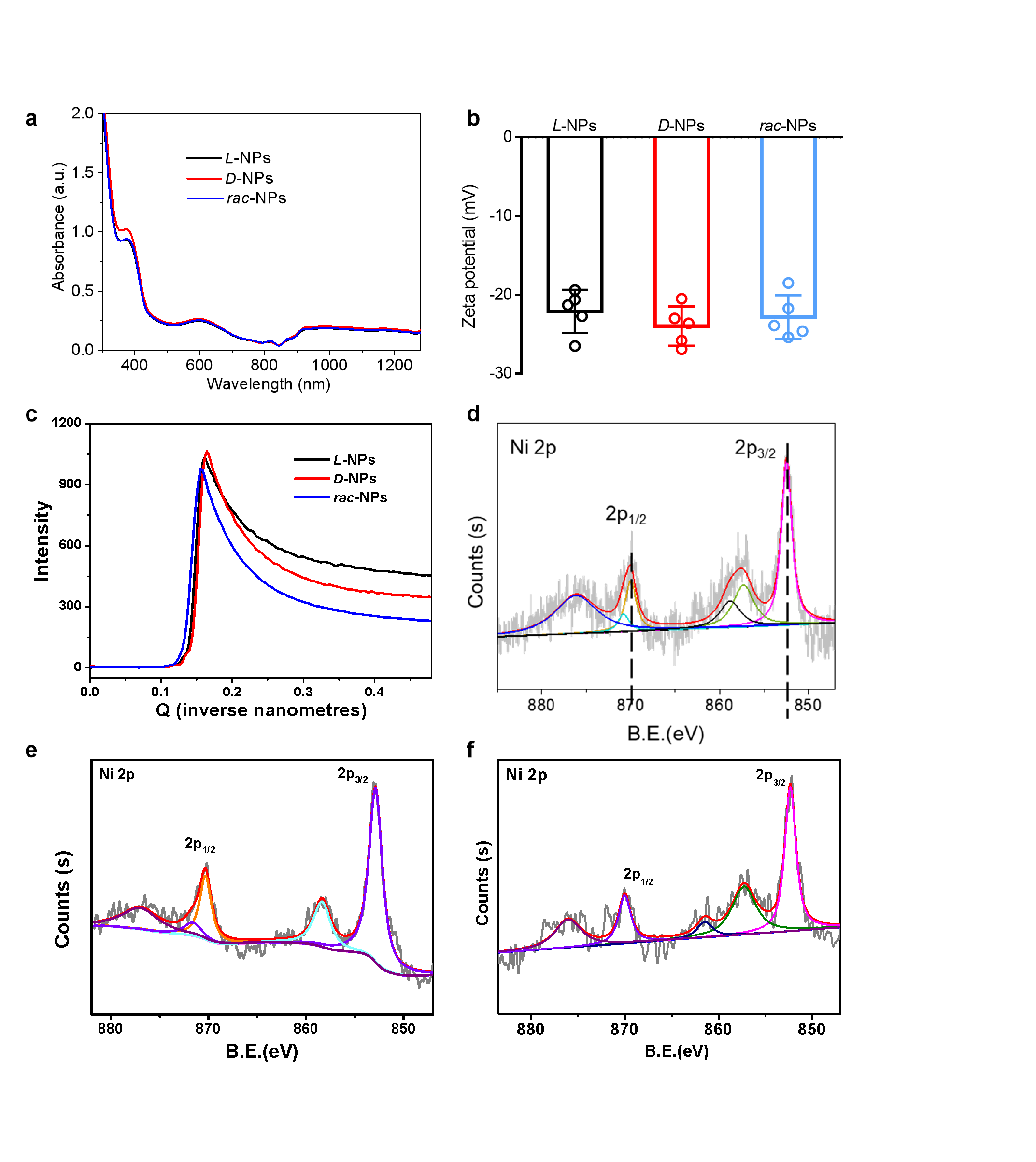

**Figure S8. (a)** UV-Vis spectra of *L*-NPs, *D*-NPs, and *rac-*NPs. **(b)** The zeta potential of *L*-NPs, *D*-NPs, and *rac*-NPs. **(c)** Small angle X-ray scattering (SAXS) of *D*-, *L*-, and *rac*-NPs. **(d-f)** X-ray photoelectron spectra (XPS) of chiral **(d)** *D*-NPs, **(e)** *L*-NPs, and **(f)** *rac*-NPs.

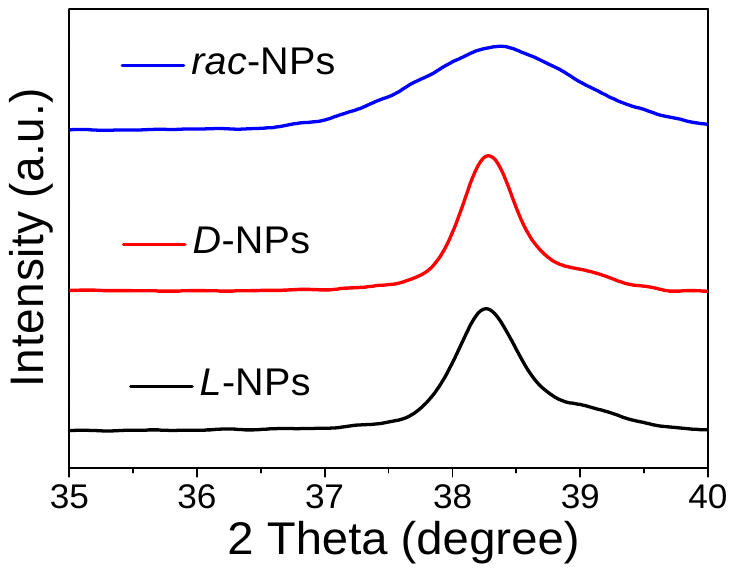

**Figure S9.** XRD spectra of *D*-, *L*-, and *rac*-NPs for 2θ between 35 and 40 degrees.

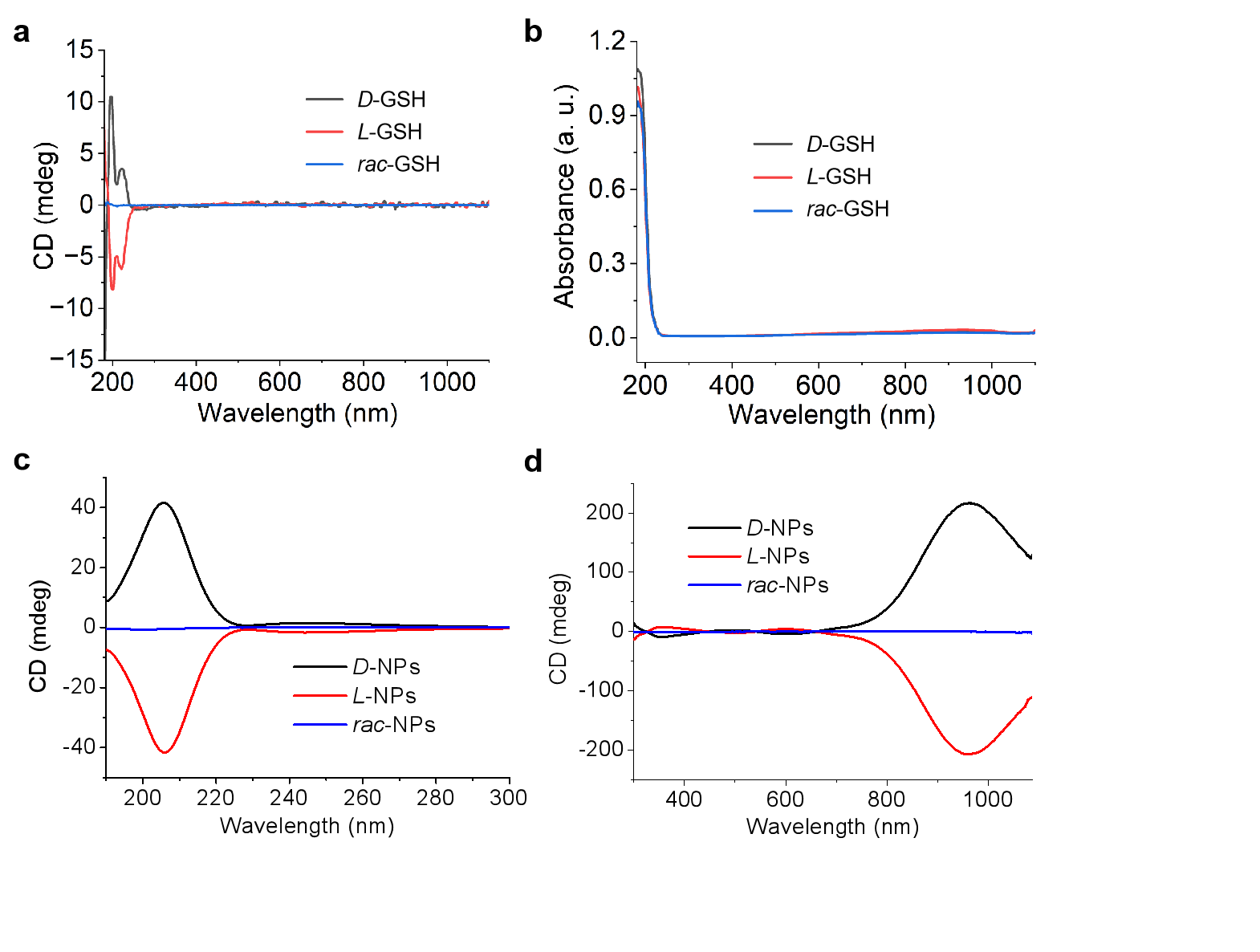

**Figure S10. (a, b)** The CD **(a)** and UV-Vis **(b)** spectra of *L-*GSH, *D-* GSH, and *rac*- GSH; **(c, d)** The CD **(c)** and UV-Vis **(d)** spectra of *L-*NPs, *D-*NPs, and *rac*-NPs.

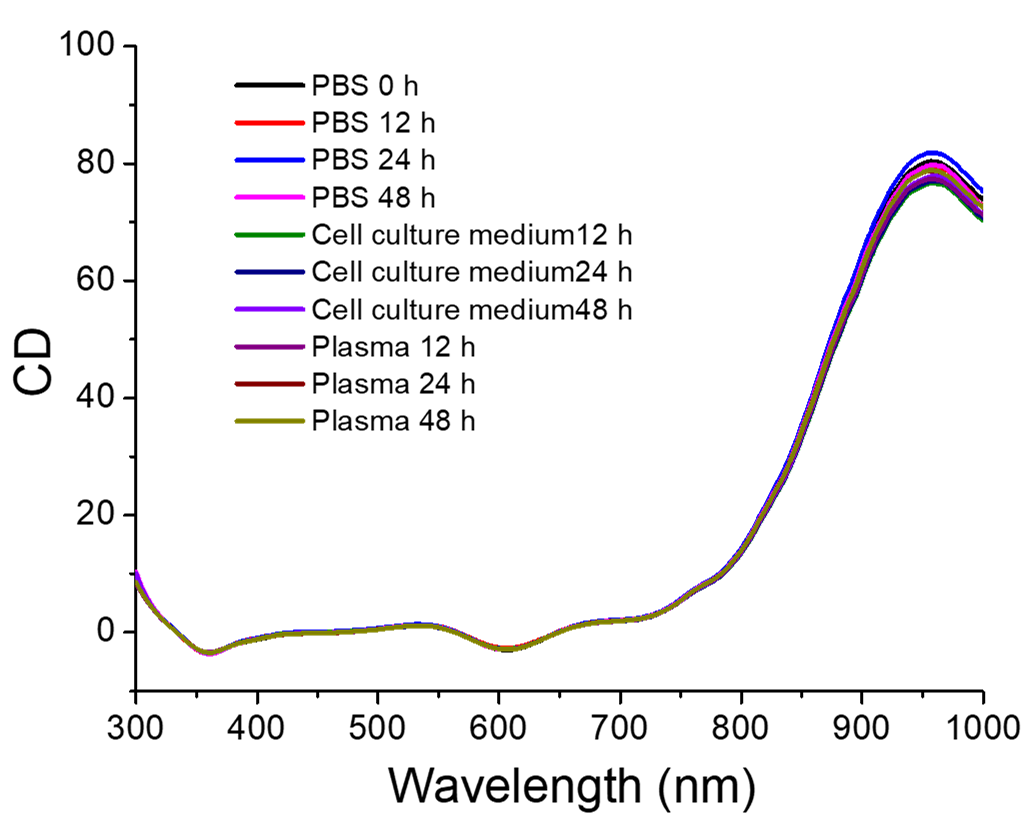

**Figure S11.** The CD spectra of *D*-NPs incubated with PBS, cell culture medium, and plasma for different times.

**Comments:** To prove the stability of chiral NPs against enzymatic hydrolysis of the GSH ligands, we cultured the *D*-NPs with PBS, cell culture medium, and plasma and then recorded CD spectra at different time points. The CD peaks show no significant changes, indicating the chiral NPs are stable against enzymatic degradation.

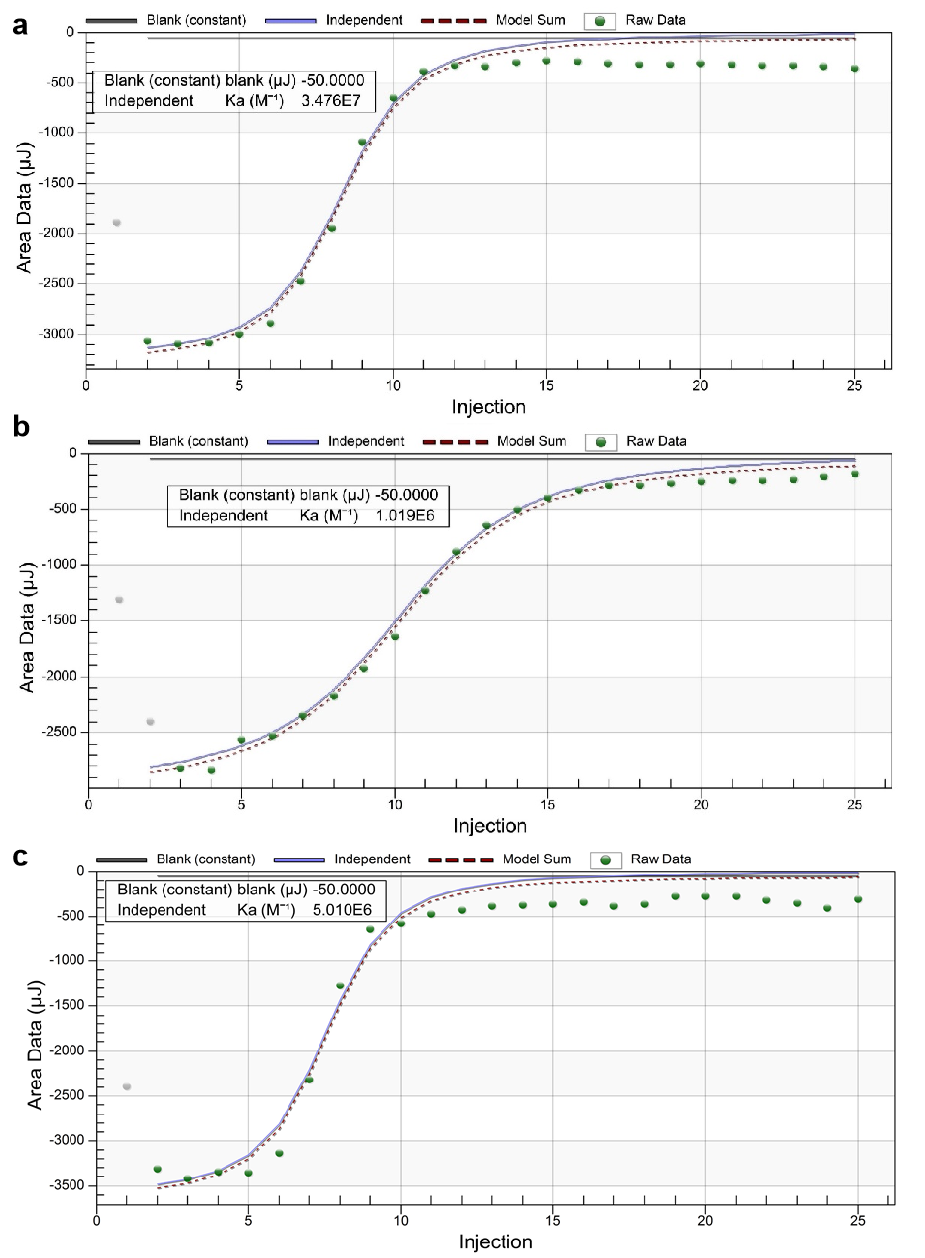

**Figure S12.** Binding affinity, *K_a_*, between NGL-1 and *D*-NPs **(a)**, *L*-NPs **(b)**, *rac*-NPs **(c)** in cell-free buffer. *K_a_* values for *D*-NPs, *L*-NPs, and *rac*-NPs are 3.476 × 10^7^ M^−1^, 1.019 × 10^6^ M^−1^, and 5.01 × 10^6^ M^−1^, respectively.

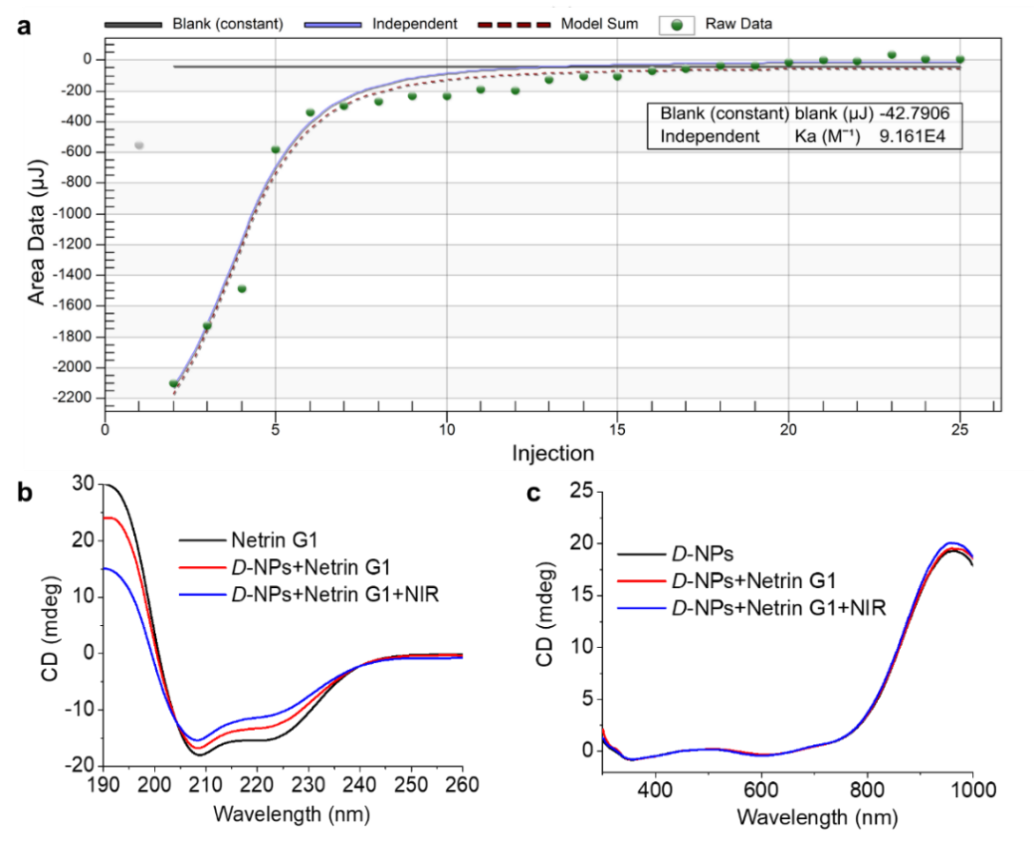

**Figure S13. (a)** Binding affinity, *K_a_*, between Netrin G1 and *D*-NPs in cell-free buffer. The *K_a_* for Netrin G1 and *D*-NPs was 9.161 × 10^4^ M^−1^. **(b, c)** CD spectra from 190 nm to 260 nm **(b)** and from 300 nm to 1100 nm **(c)** of *D*-NPs, Netrin G1, or *D*-NPs incubated with Netrin G1 at a concentration ratio of 500:1 with or without NIR illumination (980 nm, 400 mW/cm^2^, 10 min).

**Comments:** When Netrin G1 is incubated with *D*-NPs with a concentration ratio of 500:1 (*D*-NPs: Netrin G1), both the CD spectra of Netrin G1 from 190 nm to 260 nm and *D*-NPs from 300 nm to 1100 nm showed almost no change with or without NIR illumination **(Figure S13b, c)**. The value for binding affinity constant, *K_a_*, between *D*-NPs and NGL-1 (3.476×10^7^ M^-1^) was higher than the *K_a_* between *L*-NPs and NGL-1 by 34.1 times as well as between *rac*-NPs and NGL-1 by 6.9 times. It was more than 100-fold higher than that between *D*-NPs and Netrin G1 **(Figure S12** and **Figure S13a)**.

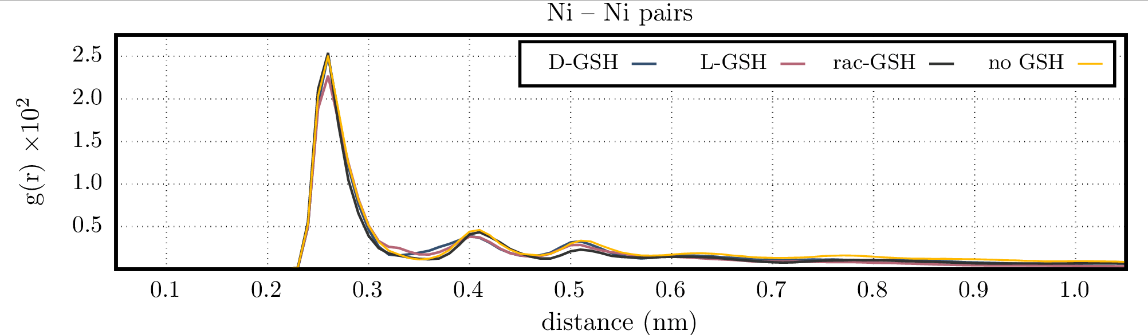

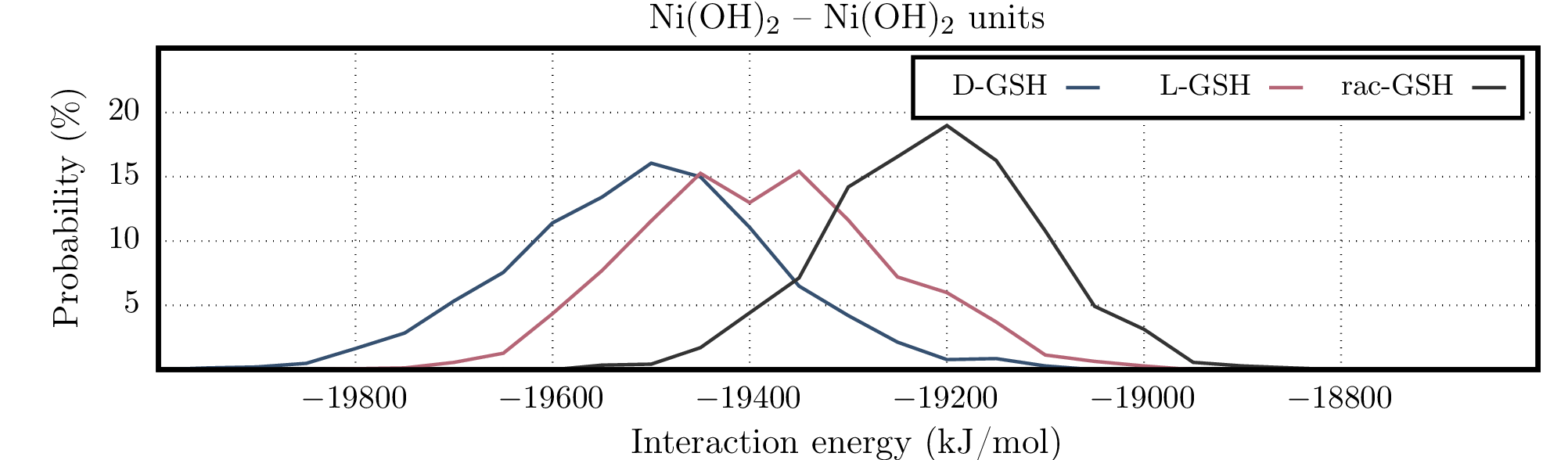

**Figure S14.** Top: Radial distribution function for Ni-Ni pairs from each molecular dynamics simulation (60–200 ns) for Ni(OH)_2_ aggregate growth. Bottom: Interaction energy between Ni(OH)_2_ units in the 60–200 ns interval.

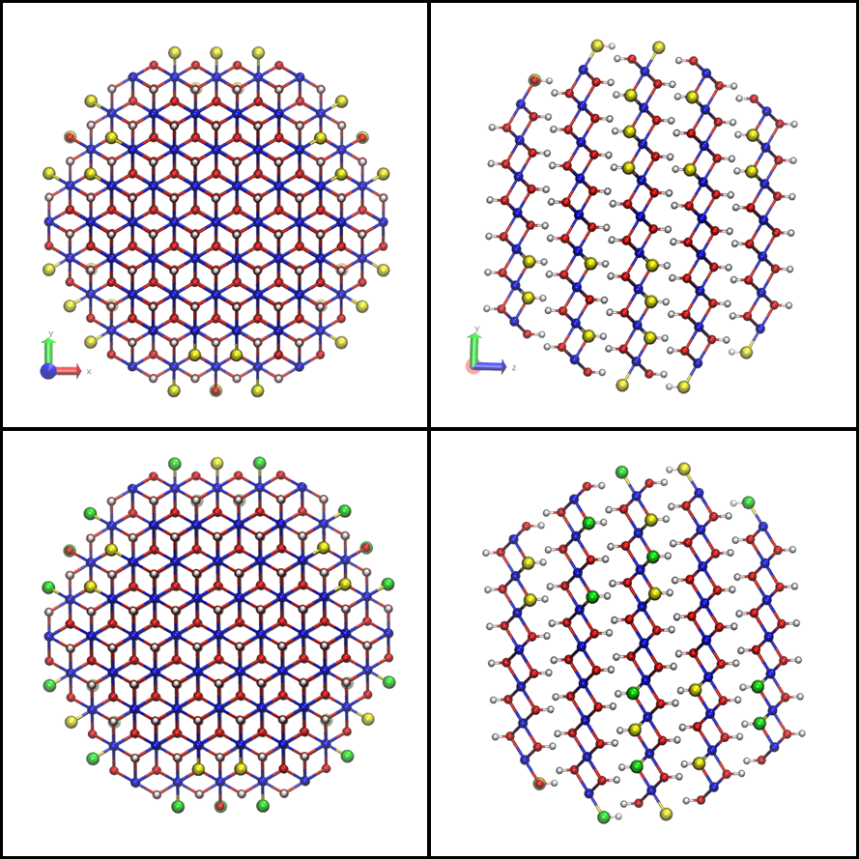

**Figure S15.** CPK representation of the model Ni(OH)_2_ NP highlighting the surface sites where GSH ligands were placed as initial models. The top row highlights all 42 low-coordination oxygen sites in yellow which were replaced by *L*-GSH ligands to yield the *L*-GSH-NP. The bottom row highlights in yellow all 21 sites which were replaced by *L*-GSH ligands and in green all 21 sites which were replaced by *D*-GSH ligands to yield the *rac*-GSH-NP. Nickel atoms are presented in blue, remaining oxygen atoms are presented in red and hydrogen atoms are presented in white.

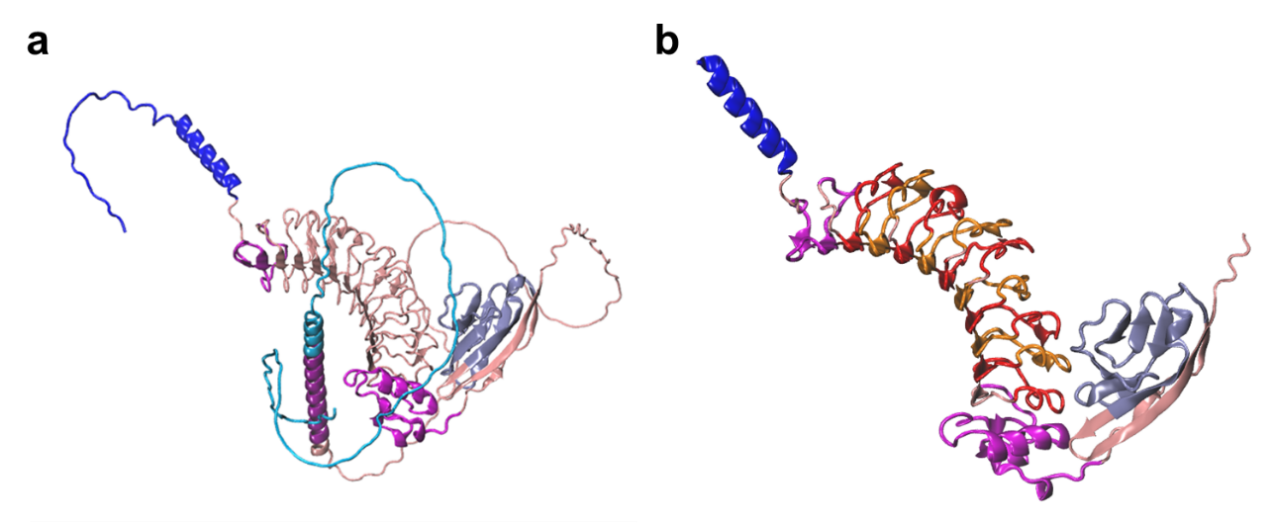

**Figure S16. (a)** Ribbon diagram structure of the full leucine-rich repeat-containing protein 4C (LRC4C) modeled by *AlphaFold* (https://www.uniprot.org/uniprotkb/Q8C031/ - label AF-Q8C031-F1). Regions are colored as follows: blue - signaling sequence (M1 to R43), magenta - leucine-rich repeat (LRR) N-terminal domain (P48 to I73), black - nine LRR (H79 to H291), the C-terminal domain LRR (N301 to Y352, highlighted in magenta) and the immunoglobulin domain (T366 to V428, highlighted in ice blue). **(b)** Ribbon diagram structure of the LRC4C fragment trimmed from P25 to P450 that was used in the simulations with the GSH-NP models.

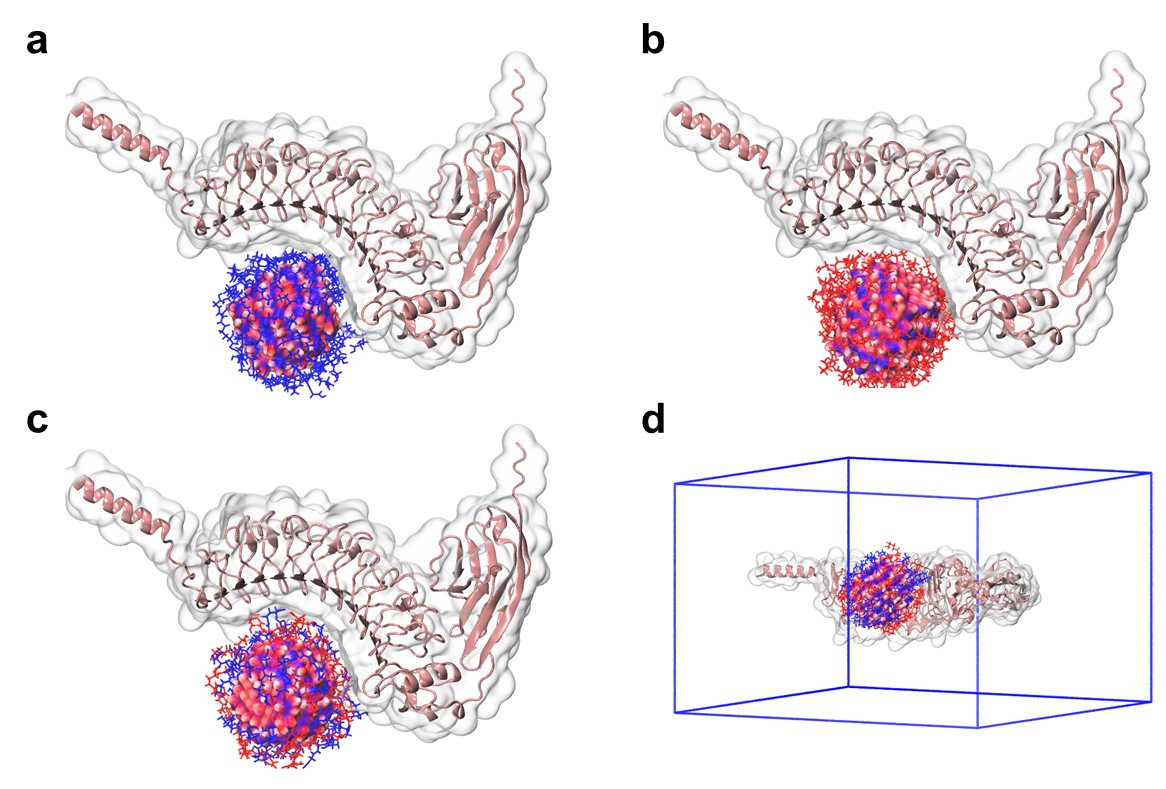

**Figure S17.** Initial structures for the GSH-NP–Protein complexes obtained after rigid-body grid-search using Themis software: **(a)** *D*-GSH-NP–Protein, **(b)** *L*-GSH-NP–Protein and **(c)** *rac*-GSH-NP–Protein. **(d)** Example of the complex centered in the simulation box. *D*-GSH molecules are depicted as blue bonds and *L*-GSH as red bonds. The atoms of Ni(OH)_2_ NP are represented as van der Waals spheres using the same color scheme described in **Figure S17**. The LRC4C molecule is depicted both by its ribbon diagram (in pink) and by the solvent accessible surface (gray transparent surface).

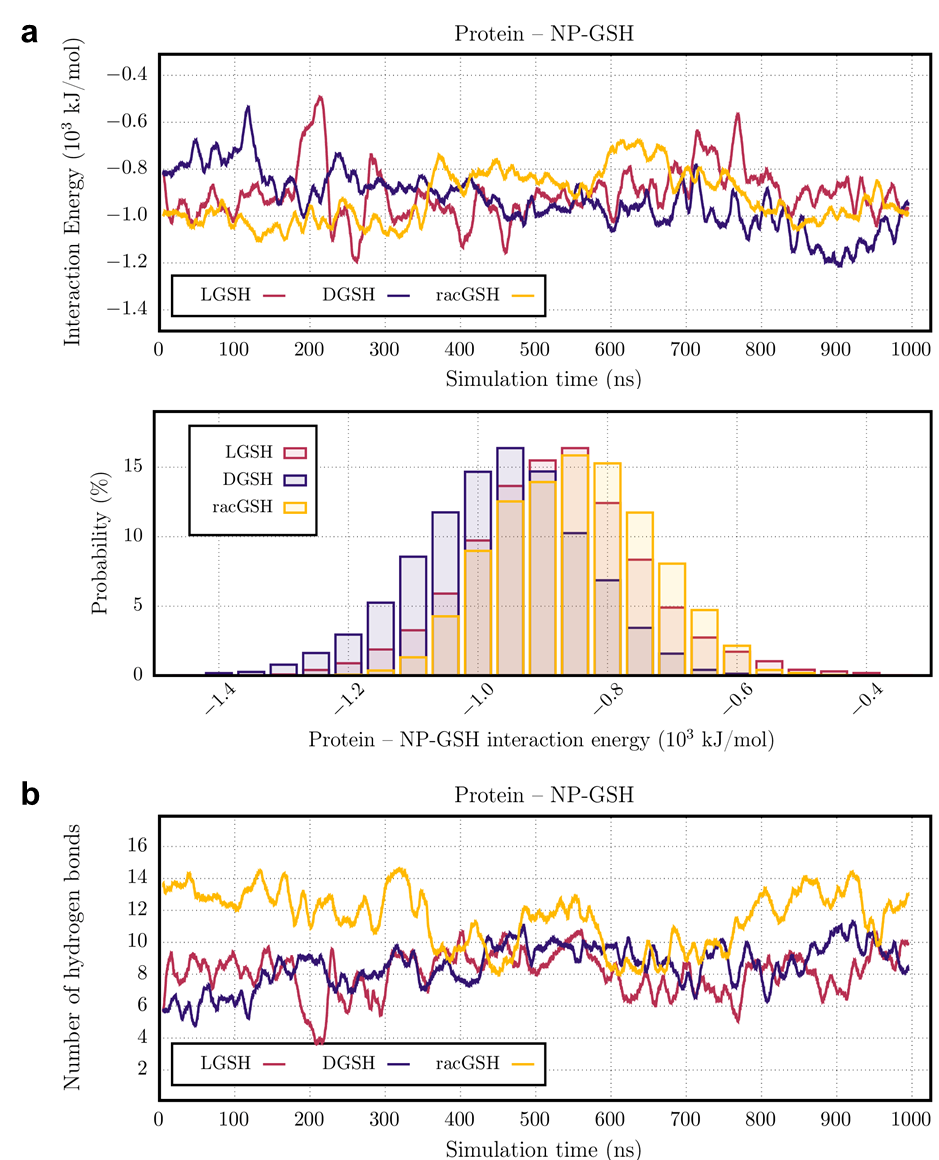

**Figure S18. (a)** Running average interaction energy (Coulomb+LJ) between functionalized NPs and the protein. **(b)** Running average number of hydrogen bonds between functionalized NPs and the protein.

**Comments：**The flexibility of NP ligands resulted in considerable interaction energy fluctuations during the production runs, so we present the 200-point running average for the interaction energies **(Figure S18a)**, which had a slow relaxation during the first 350 ns. The average interaction energies obtained from 350-1000 ns were –994 ± 127 kJ/mol for the *D*-NP, –903 ± 135 kJ/mol for the *L*-NP, and –857 ± 117 kJ/mol for the *rac*-NP. Although the averages seem close to each other as compared to the error estimates, the running average shows that the interaction energies for the *D*-NP is consistently lower (more negative) than the values for the *L*-NP and *rac*-NP, in this time interval, except for two interaction peaks of *L*-NP in the range between 400 and 500 ns.

All three NPs remained in contact with the protein fragment, even though considerable energetic fluctuations were observed along the simulations, but no dissociation event was observed (in which case potential energy should become null). The favorable interactions arise mainly from a strong hydrogen bonding network between the ligands of the NPs and the protein fragment **(Figure S18b)**, whose average number in the window between 350 and 1000 ns amounted to 10.9 ± 2.5 for *rac*-NP, 9.1 ± 1.9 for *D*-NP, and 8.3 ± 2.1 for *L*-NP.

| 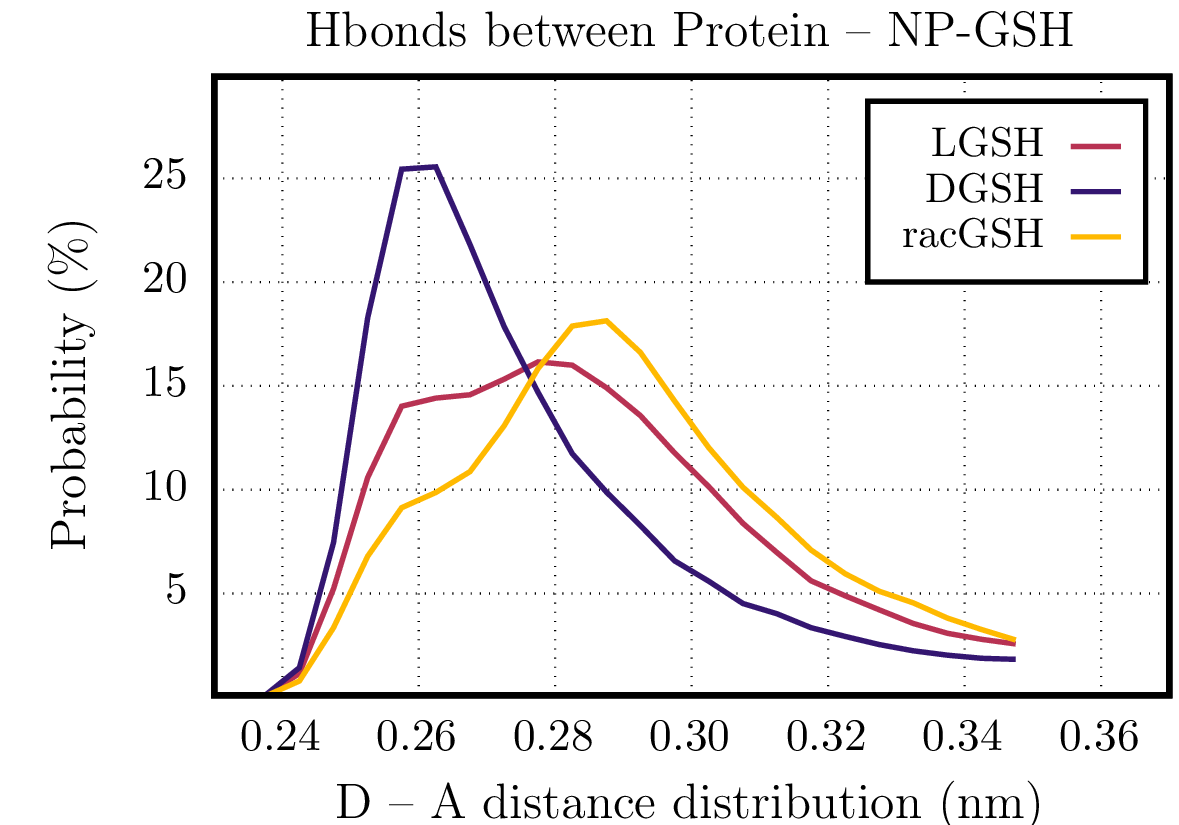 | 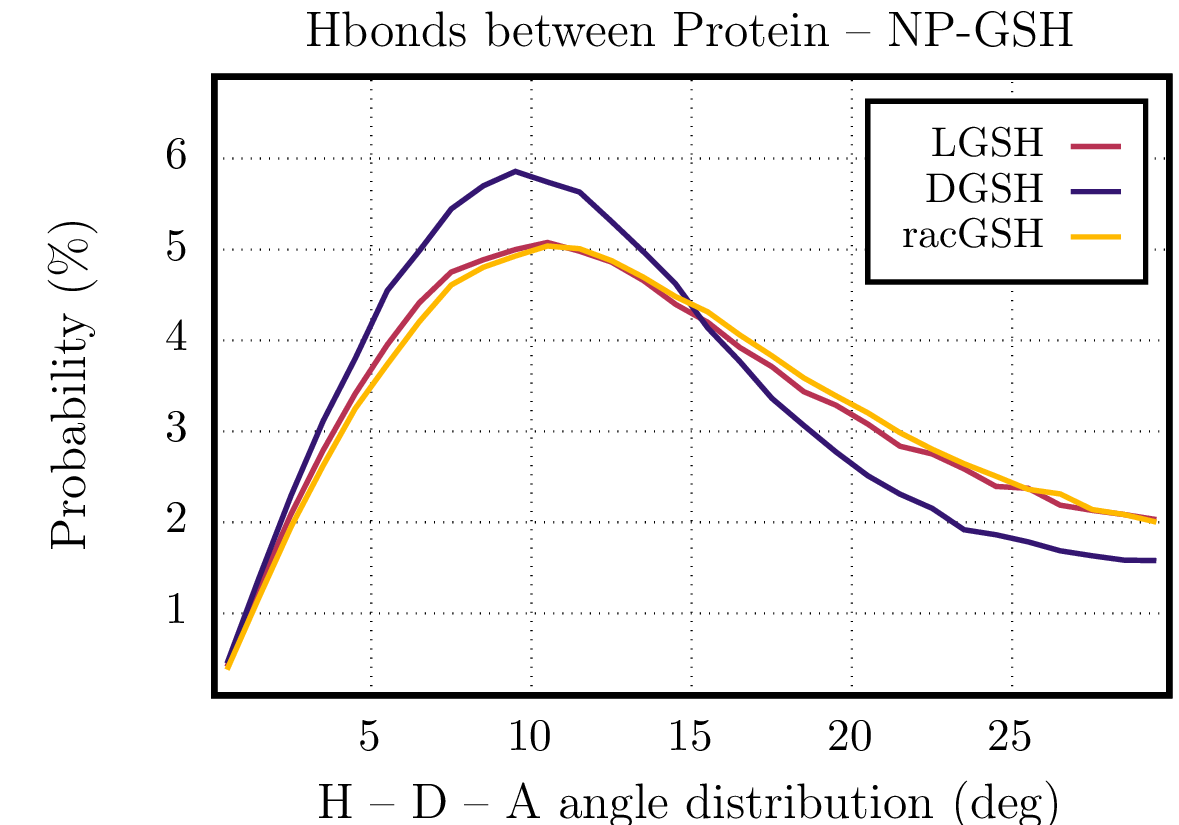 |
| --- | --- |

**Figure S19.** Donor-acceptor (D-A) distance distribution (left) and hydrogen-donor-acceptor (H-D-A) angle distribution (right) for the hydrogen bonds between the functionalized NPs and the protein.

**Comments:** The binding becomes stronger, but the total number of hydrogen bonds become smaller for *D-*NPs compared to the *L-* and *rac-*NPs. This finding indicates that the handedness of the ligands directly affects the efficiency of the hydrogen bonding, since a smaller number of interactions for the *D*-NPs yielded the stronger interaction. This reasoning is supported by the analysis of the distribution of donor-acceptor (D-A) distances and hydrogen-donor-acceptor (H-D-A) angles, which are sharper and shifted towards smaller values for the *D*-NPs as compared to the *L*-NPs and *rac*-NPs **(Figure S19)**. All data showed that the difference in the interaction energies arise from distortions of geometric parameters defining the H-bonds depending on the handedness of the NPs.

| 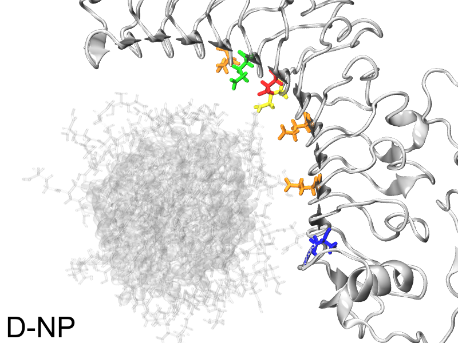 | 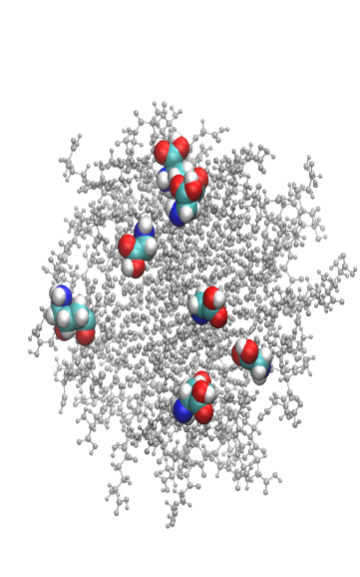 |
| --- | --- |
| 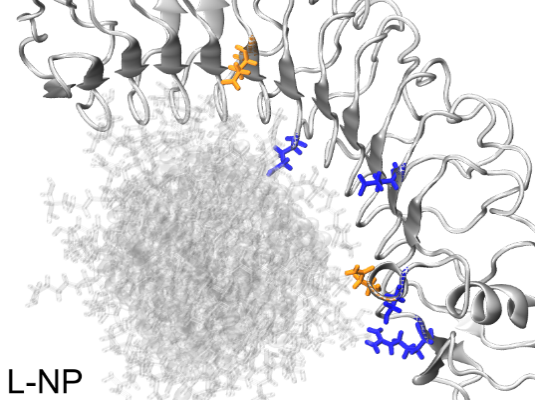 | 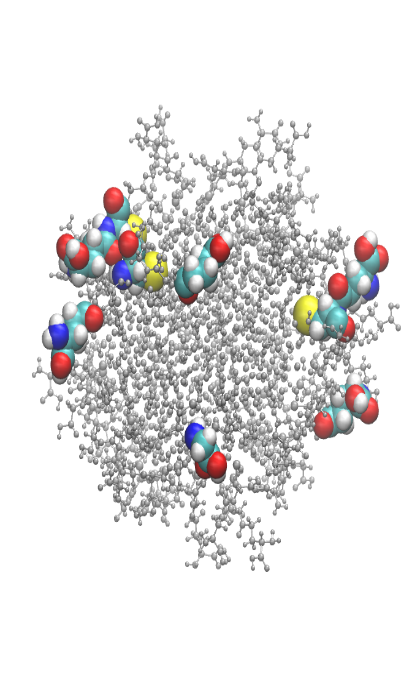 |
| 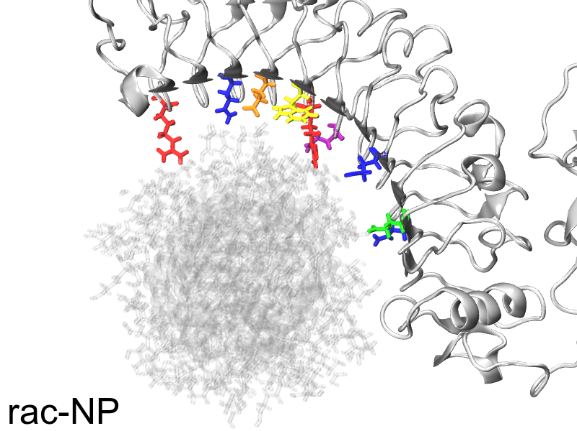 | 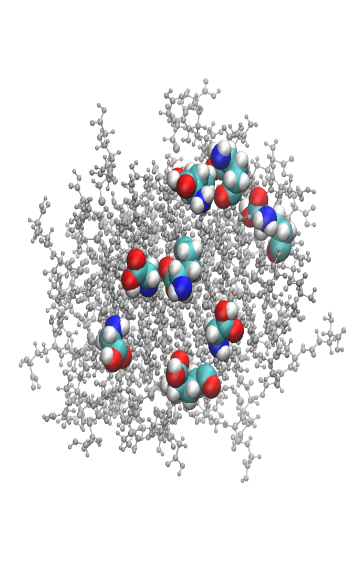 |

**Figure S20.** Final structures of GSH-NP–Protein complexes (left) and nanoparticles (right) after 1000 ns of MD simulations. Protein residues with hydrogen bond occupancy values larger than 30% are highlighted as follows: 30-50% (blue), 50-60% (green), 60-70% (yellow), 70-80% (orange), 80-90% (purple), and larger than 90% (red). The nanoparticles are depicted with the GSH fragments directly involved in hydrogen bonds (**Tables S2-S4**) as van der Waals structures.

| 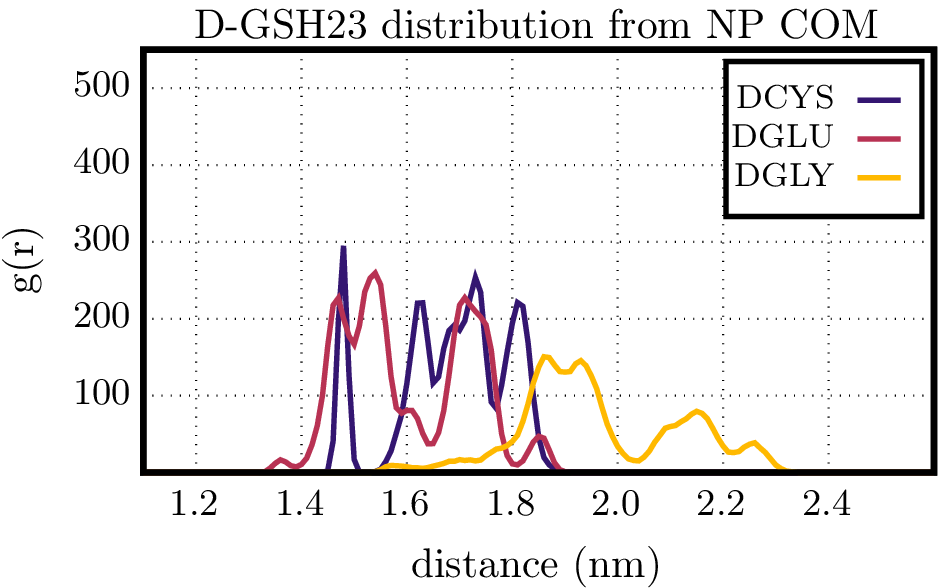 | 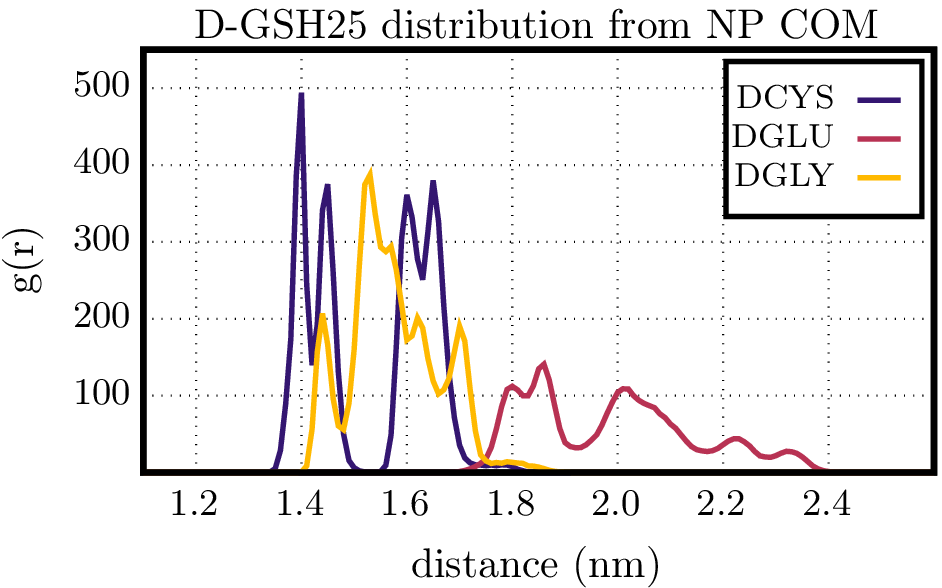 |
| --- | --- |
| 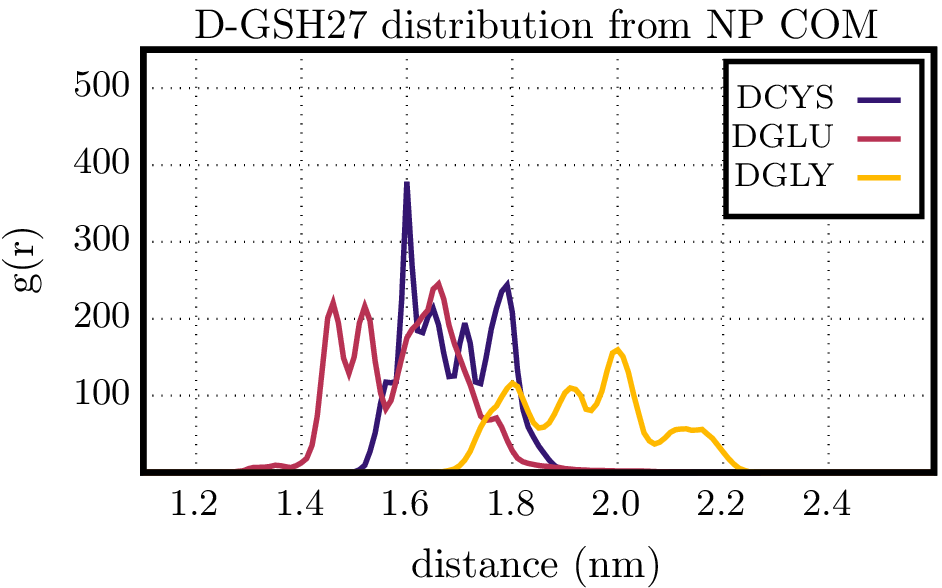 | 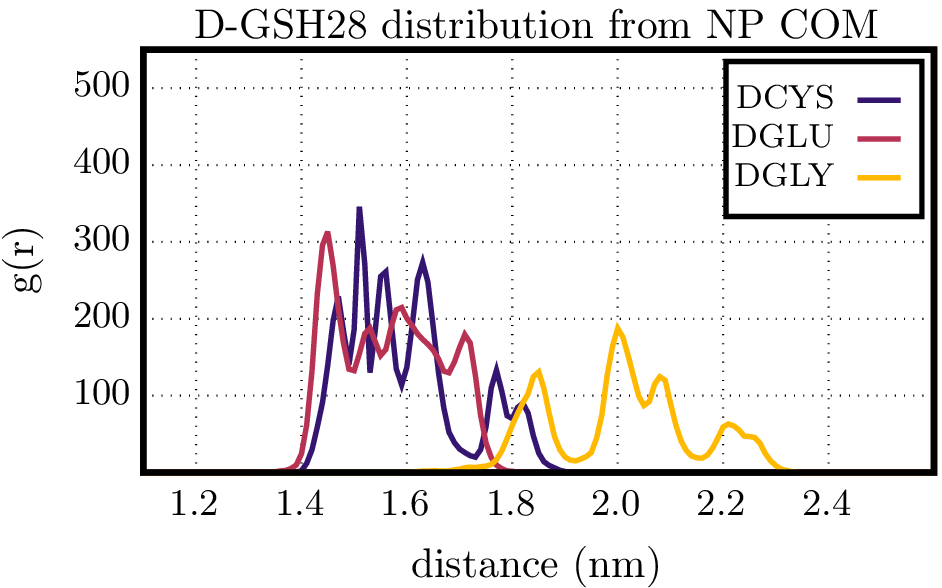 |
| 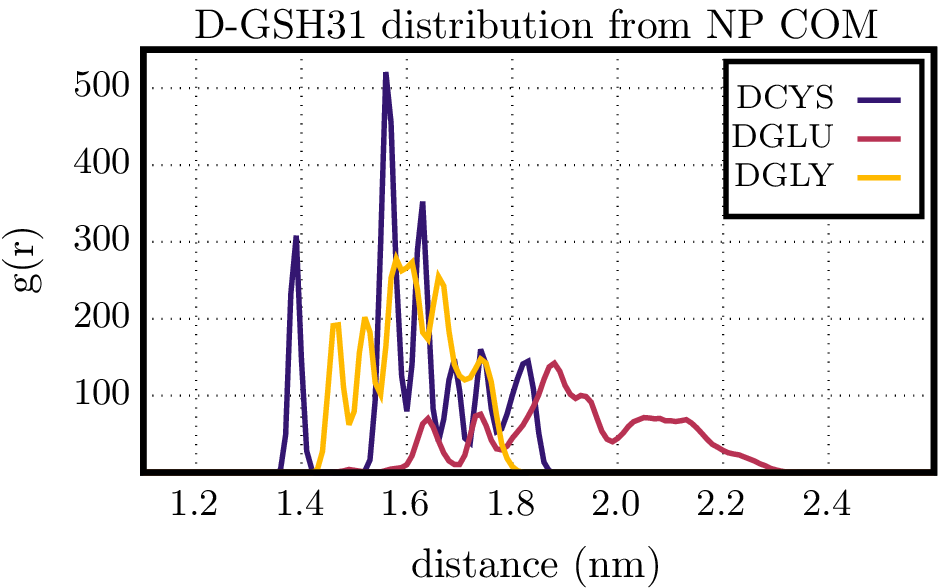 | 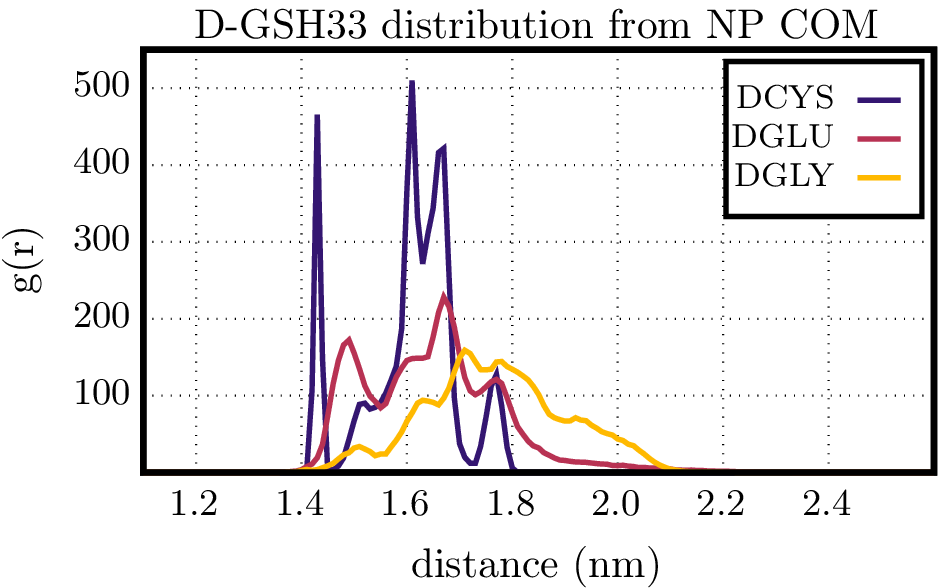 |
| 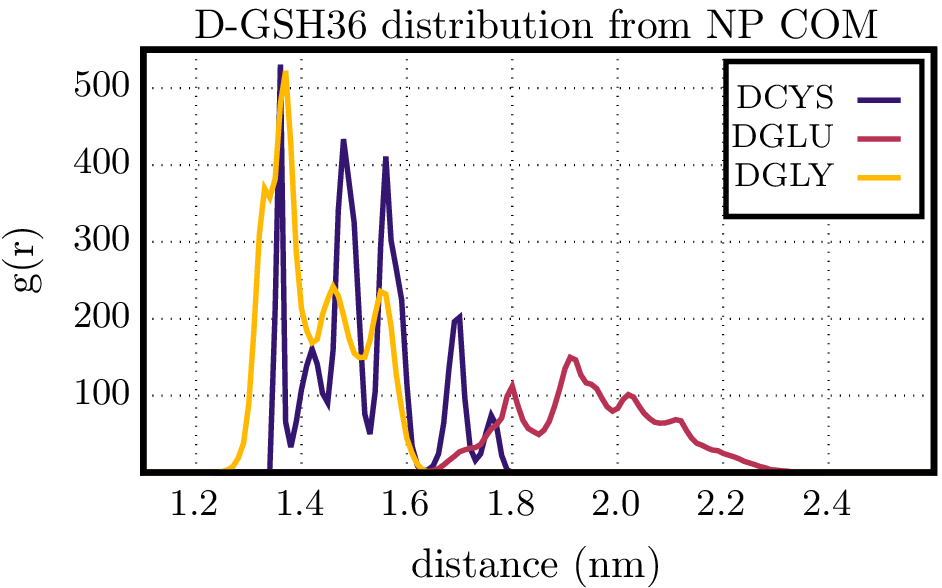 |  |

**Figure S21.** Radial distribution function of the amino acids comprising the *D*-GSH molecules hydrogen-bonded to the protein (**Table S2**) with respect to the NP center of mass.

| 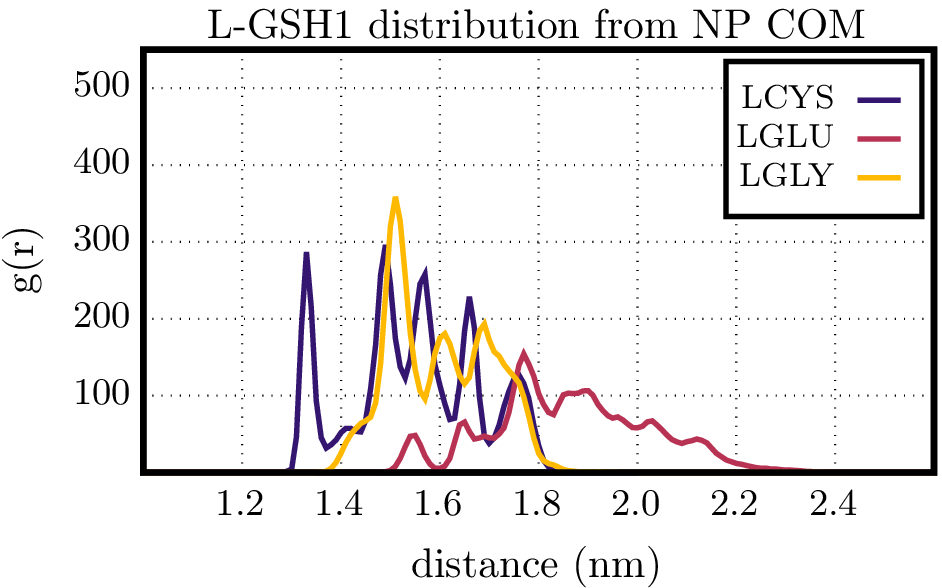 | 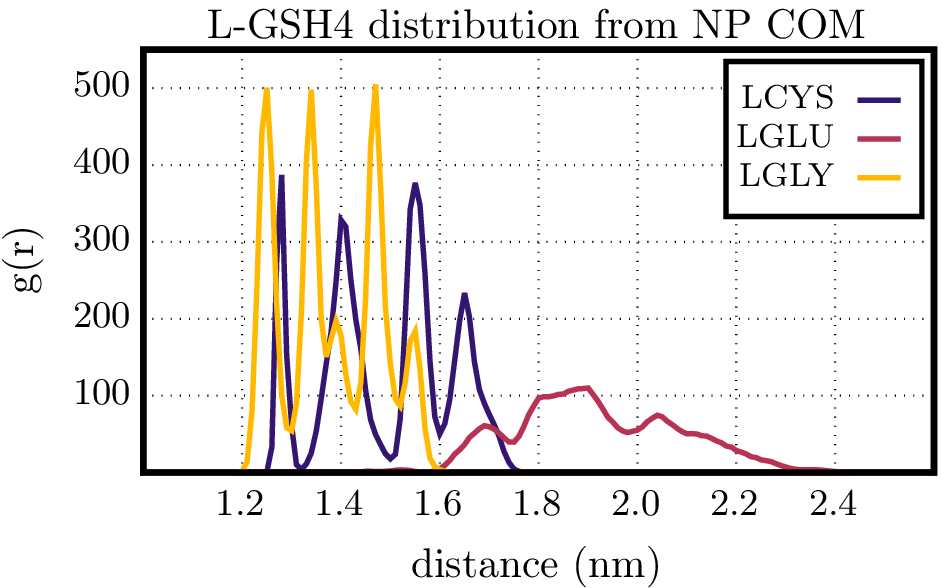 |
| --- | --- |
| 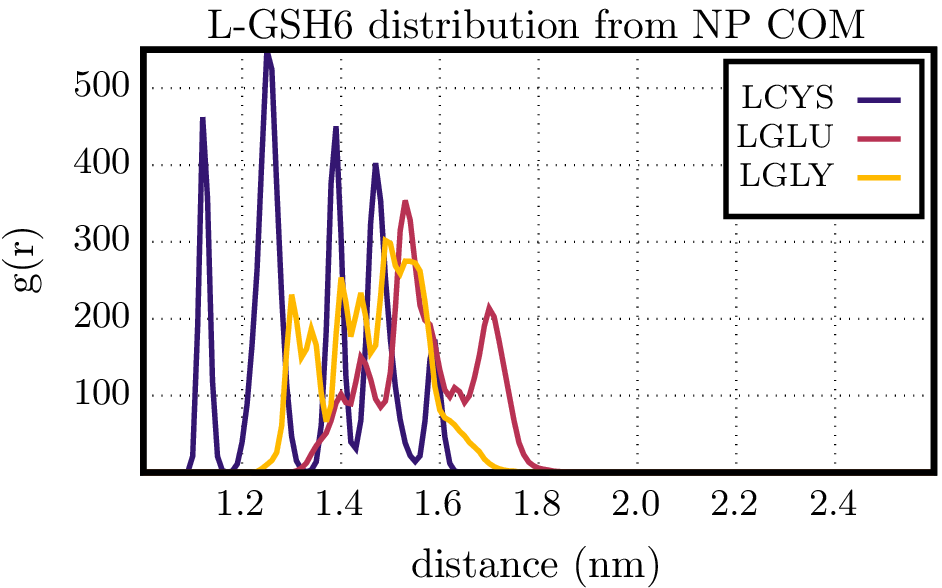 | 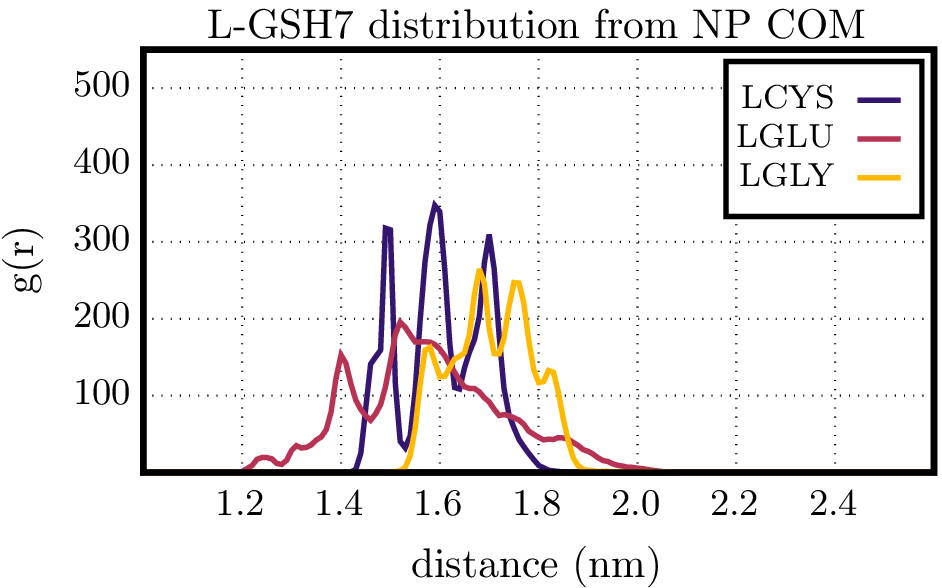 |
| 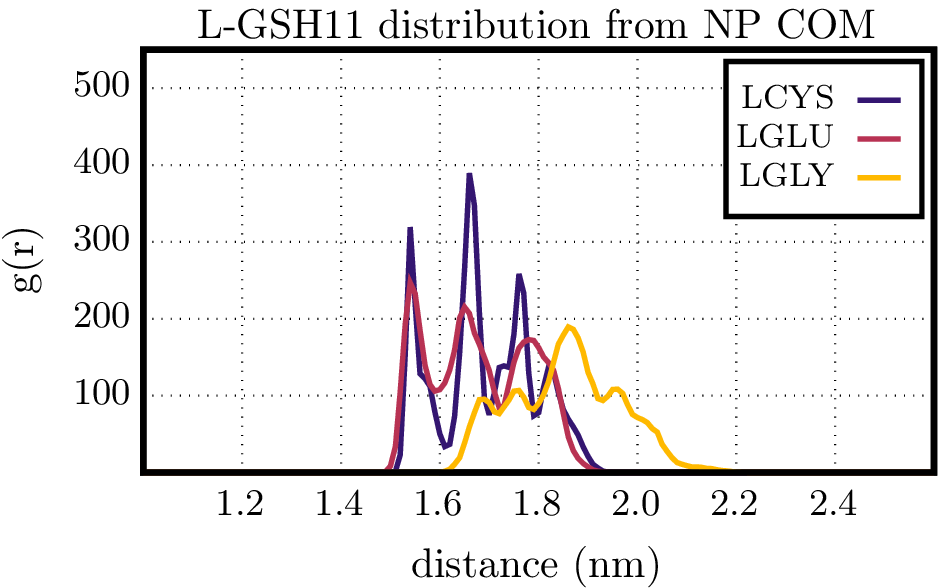 | 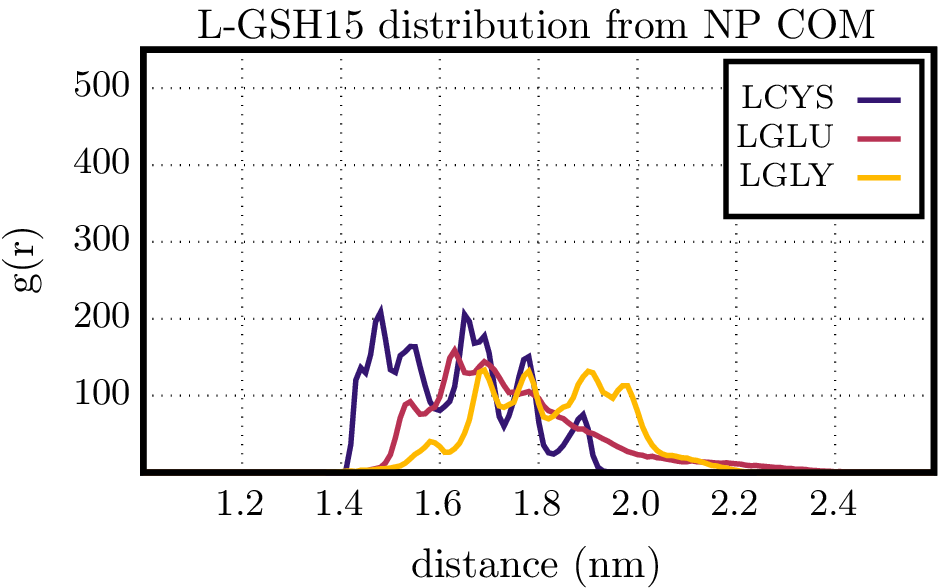 |
| 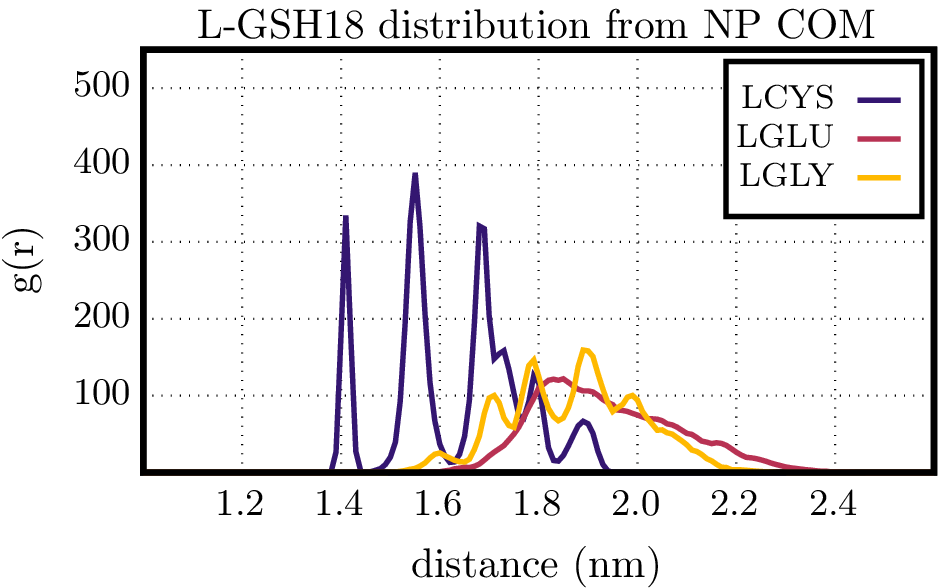 | 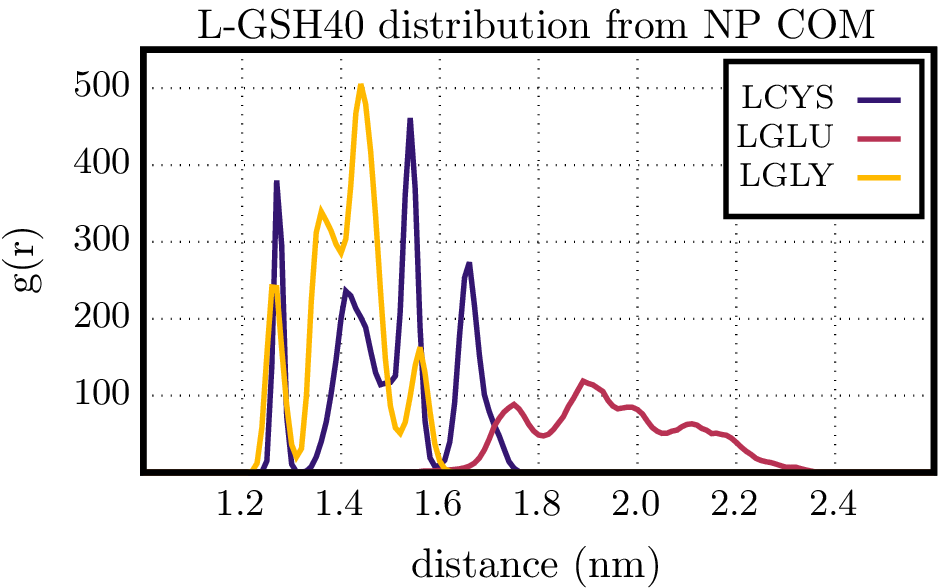 |

**Figure S22.** Radial distribution function of the amino acids comprising the *L*-GSH molecules hydrogen-bonded to the protein (**Table S3**) with respect to the NP center of mass.

| 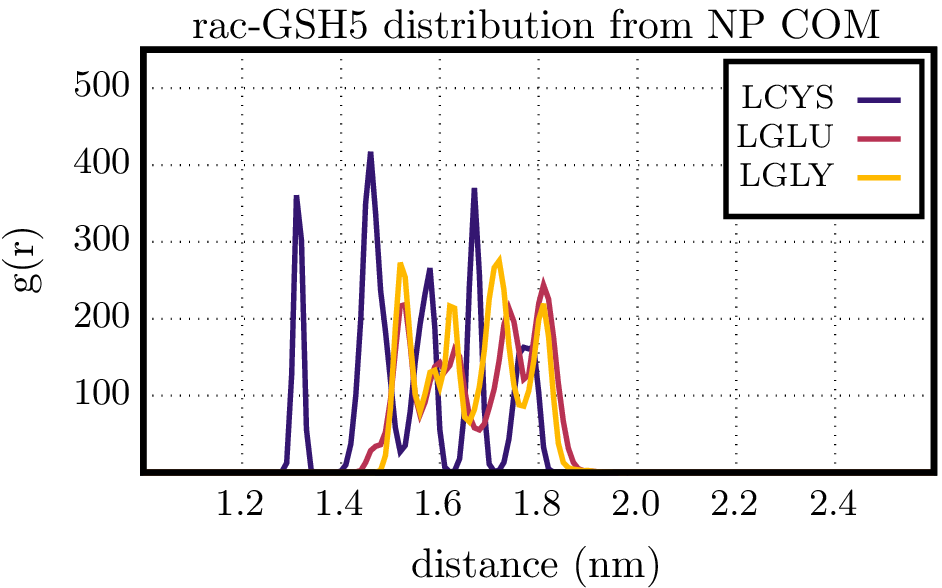 | 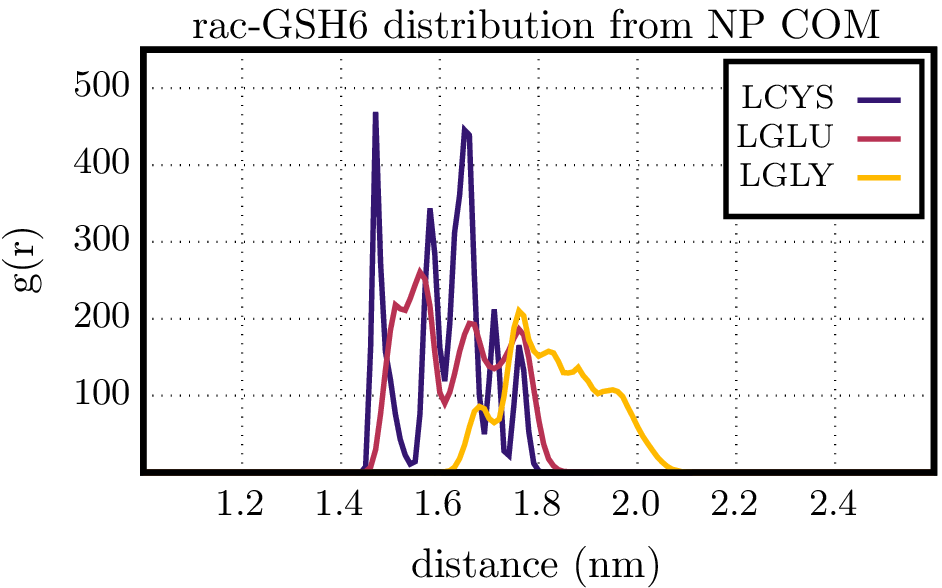 |
| --- | --- |
| 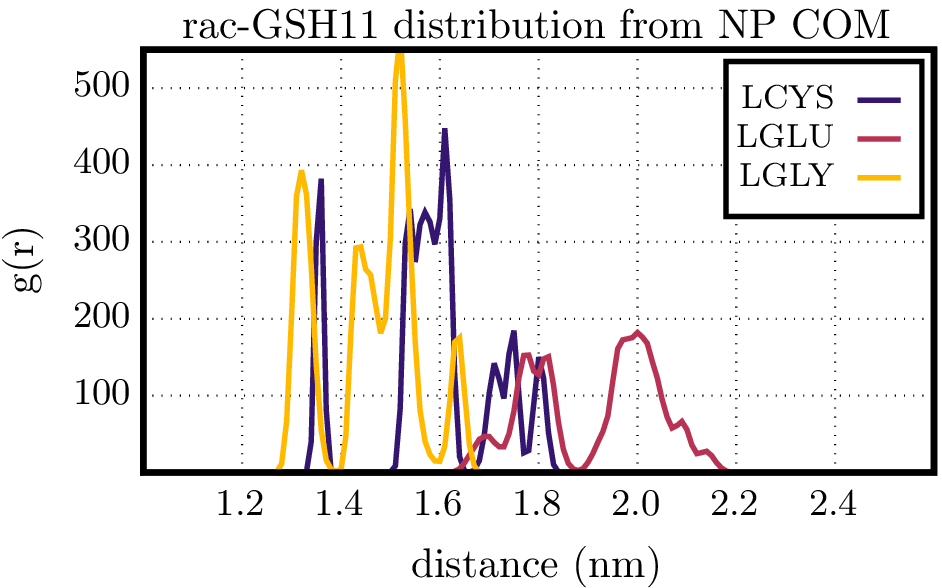 | 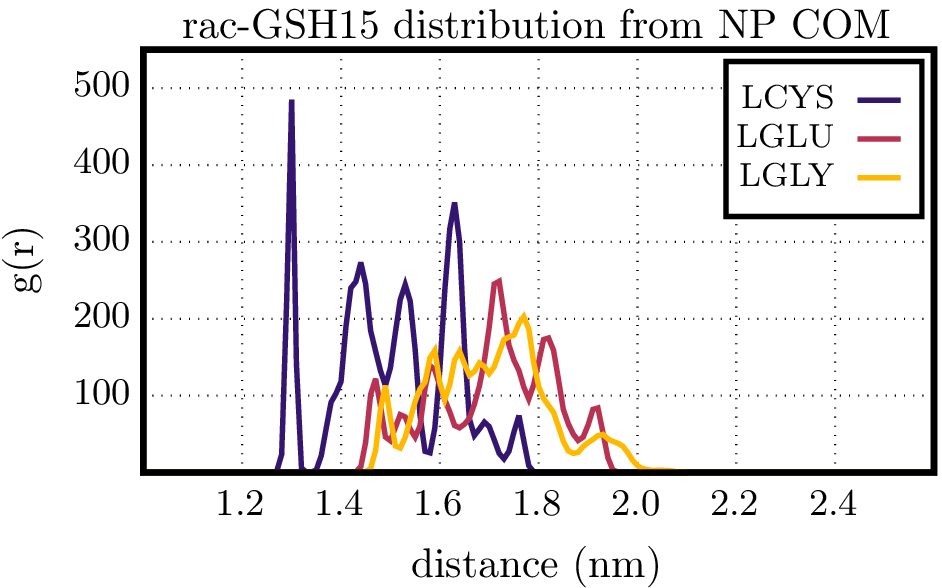 |
| 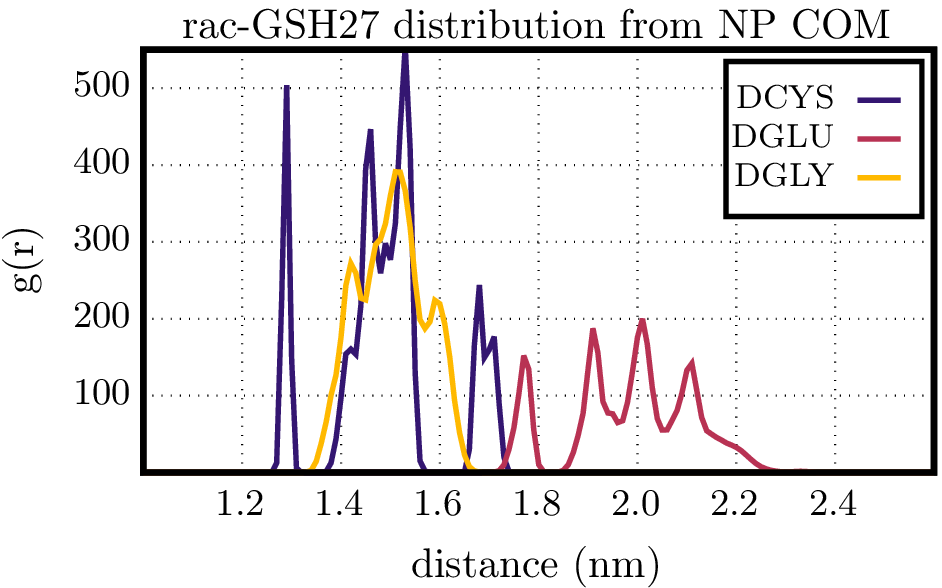 | 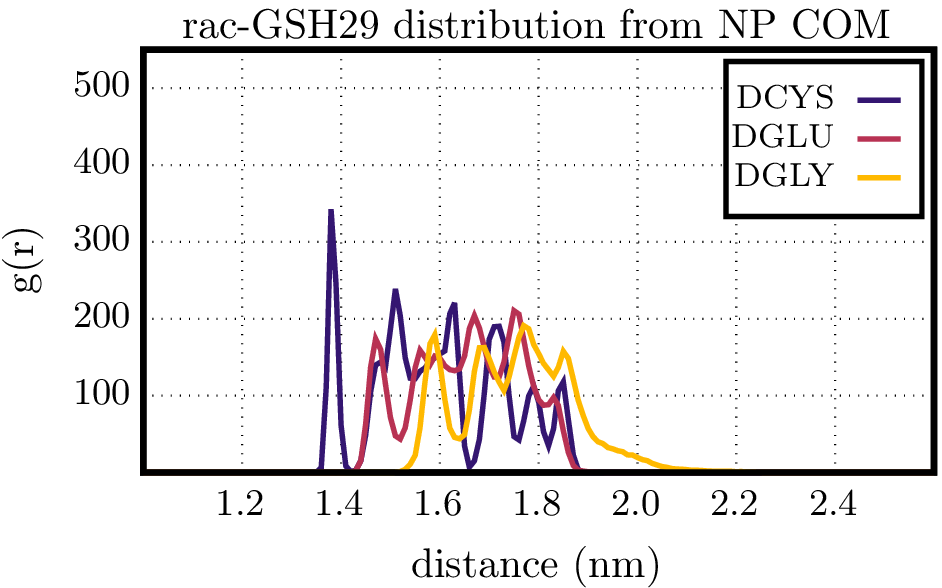 |
| 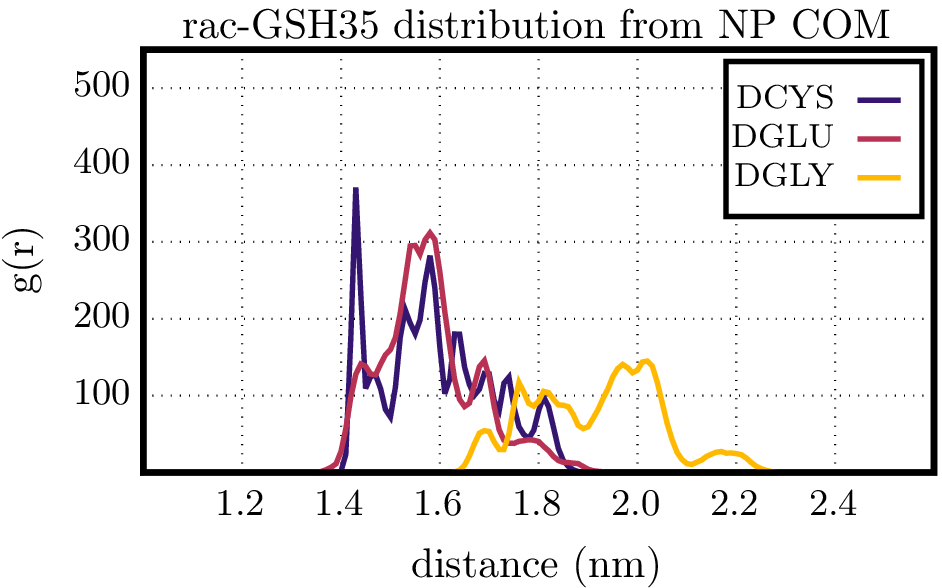 |  |

**Figure S23.** Radial distribution function of the amino acids comprising the *D*-GSH and *L*-GSH molecules hydrogen-bonded to the protein (**Table S4**) with respect to the NP center of mass.

**Comment:** As a general trend, most hydrogen bond interactions involved GLU and ASP anionic residues from the protein and, more specifically, residues GLU181 and GLU271 formed hydrogen bonds with all GSH-NPs, but with different occupancy in each case (**Table S2-S4**). Visual inspection of the last frame of each simulation shows that besides differing in occupancy, the spatial distribution of hydrogen bonds is also different for each enantiomer of the GSH-NP, with *D*-GSH-NP having nearly equally spaced interactions in the central region of the concave fragment of the protein where the NP fits in, while the *L*-GSH-NP interactions are more sparse and the *rac*-GSH-NP interacts preferably with the terminal portion of the protein (**Figure S20**). The GSH ligands bind to the NP surface through the thiolate of the central CYS amino acid, which remains closer to the surface (**Figure S21-S23**) and mostly unavailable to interact with the protein (only for the *L*-GSH-NP the CYS formed hydrogen bonds with the protein using its carbonylic oxygen atom attached to the backbone). Although GLY is often referred to as an apolar amino acid because its side chain is an aliphatic H atom, which is formally apolar, we must acknowledge that this amino acid has highly accessible carboxylic acid and amino groups, and although it is the shortest amino acid it can protrude away from the NP surface, sometimes farther away than GLU, which has a longer side chain (**Figure S21-S23**). It is also important to acknowledge that although GLY is not chiral on its own, it belongs to a chiral environment formed by CYS and GLU, so it is referred to as either *L*-GLY or *D*-GLY in these analyses because its interactions are tuned by the overall chirality of each GSH enantiomer.

Shorter collinear hydrogen bonds tend to be stronger due to the optimal arrangement of charged atoms and as such they have a larger contribution to the stabilization of the protein-NP complexes. So, it is not the total number of hydrogen bonds that stabilizes the complexes, their relative strength needs to be considered as well.

In the case of the *L*-GSH we should also mention the interactions with CYS from the GSH residue, which was the more distorted hydrogen bond observed for all systems, as expected for a residue closer to the NP surface and farther from the solution (**Figure S24**).

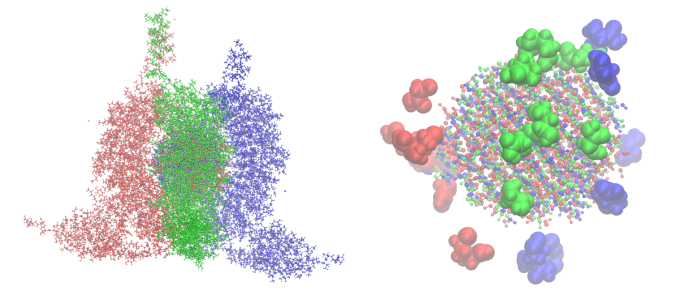

**Figure S24.** NP-protein complexes with the Ni(OH)_2_ cores aligned at the center (left) and NP core with the GSH amino acids directly involved in hydrogen bonds (right). In each representation the atoms from the *D*-GSH-NP system are depicted in red, from the *L*-GSH-NP system in blue and from the *rac*-GSH-NP in green.

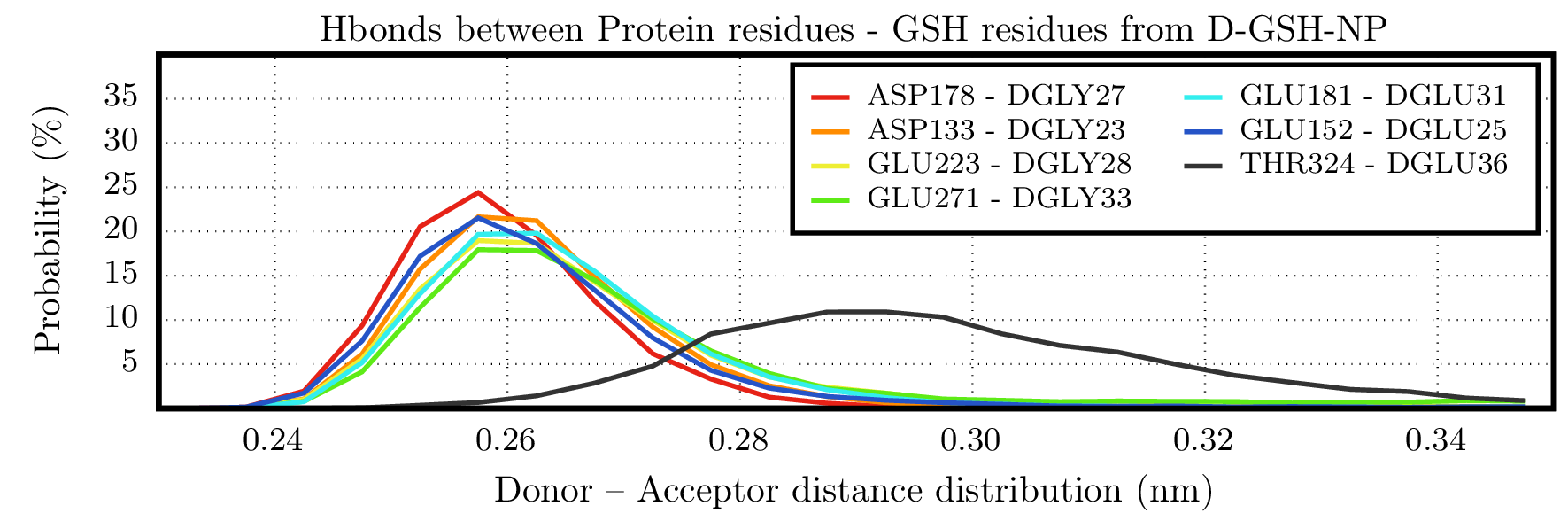

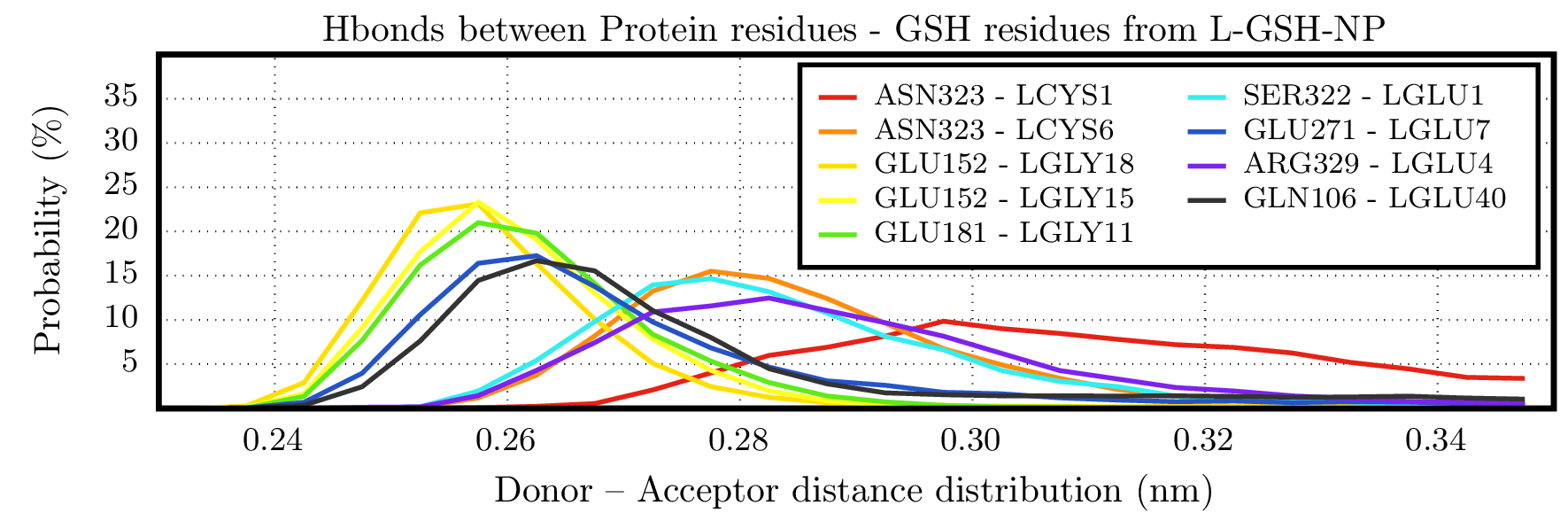

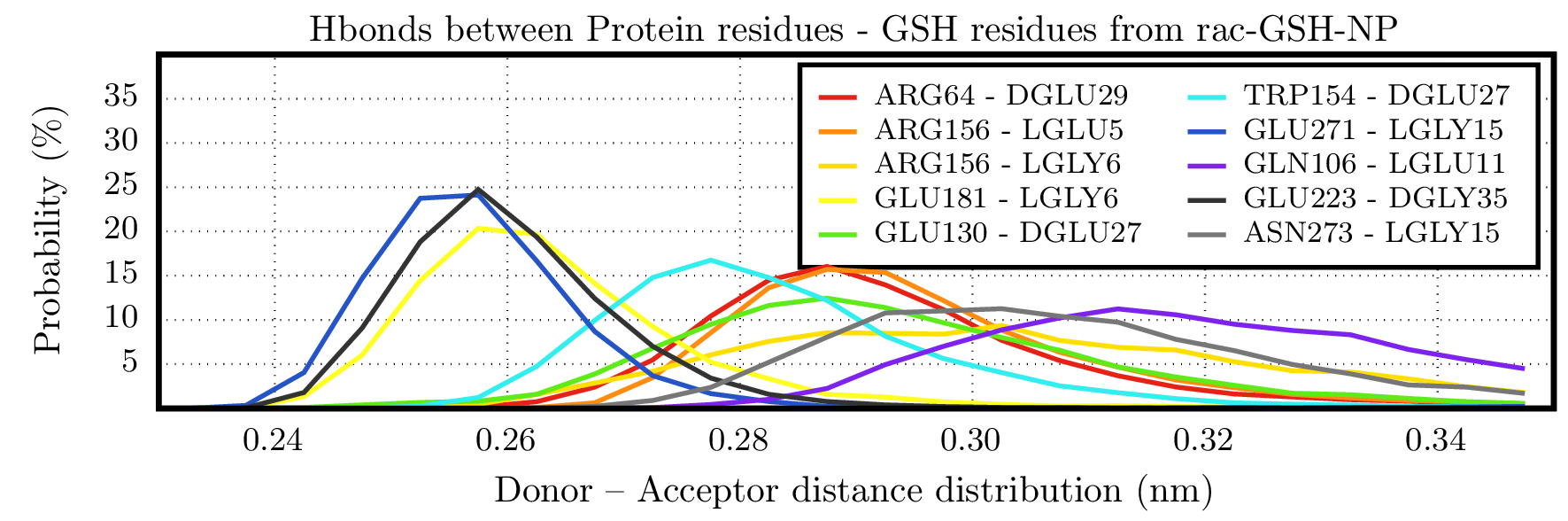

**Figure S25.** Hydrogen Donor-Acceptor (D-A) distance distribution for the protein residues, which presented the highest hydrogen bond occupancy values.

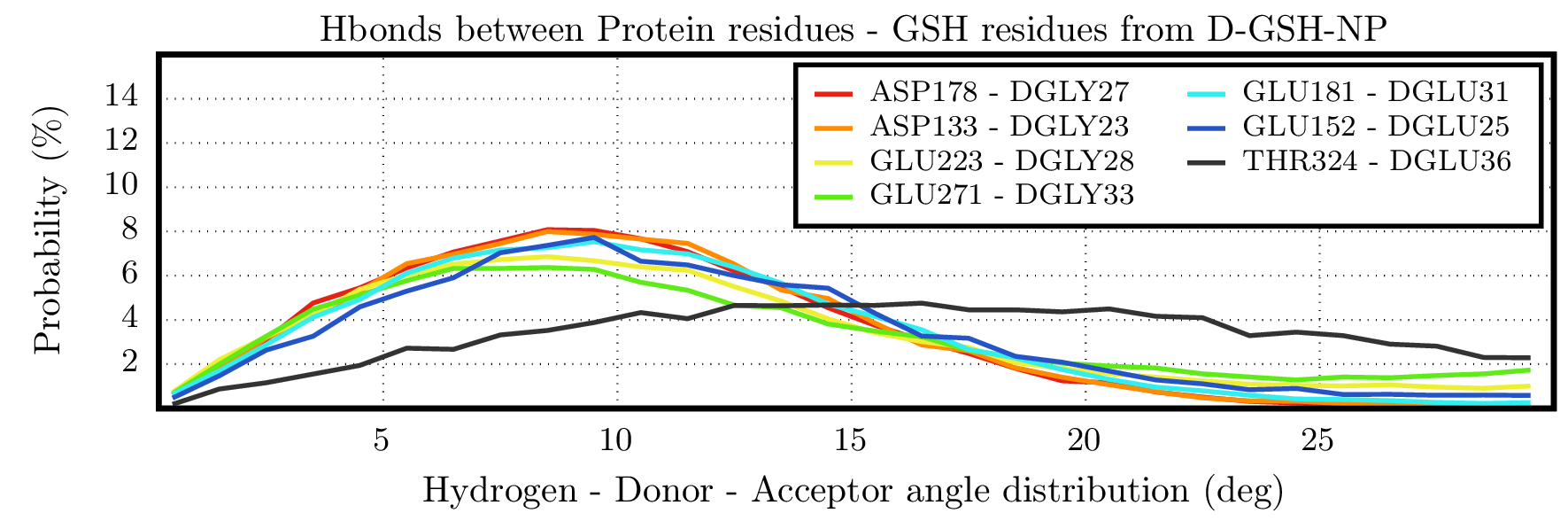

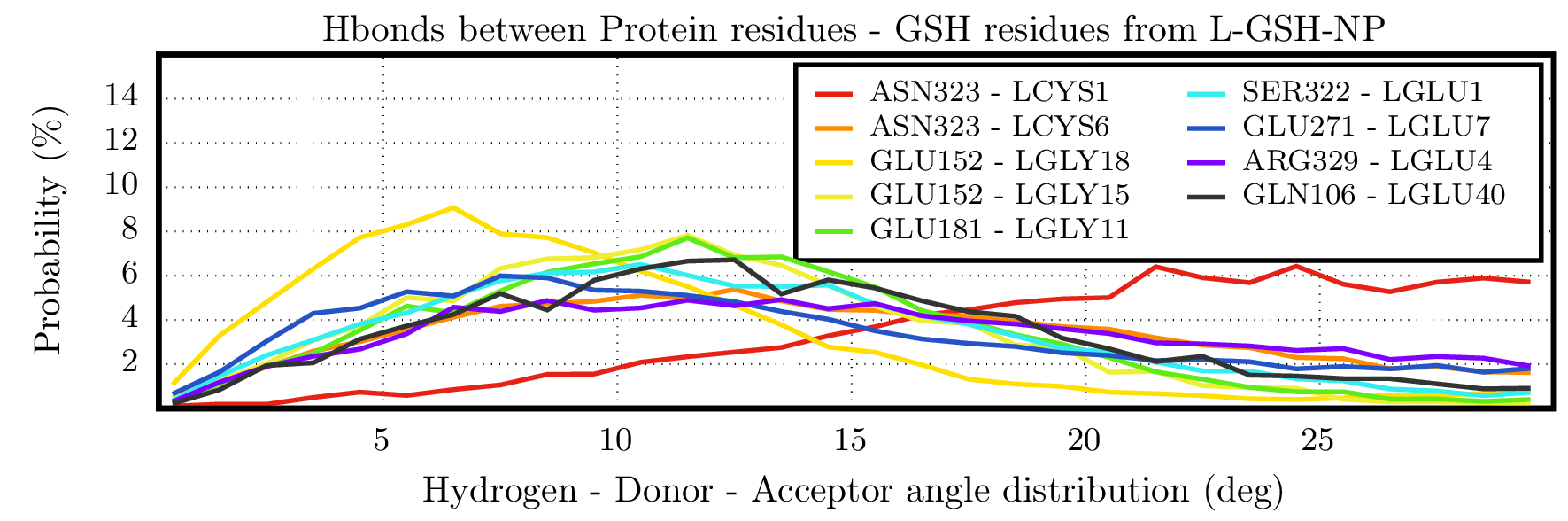

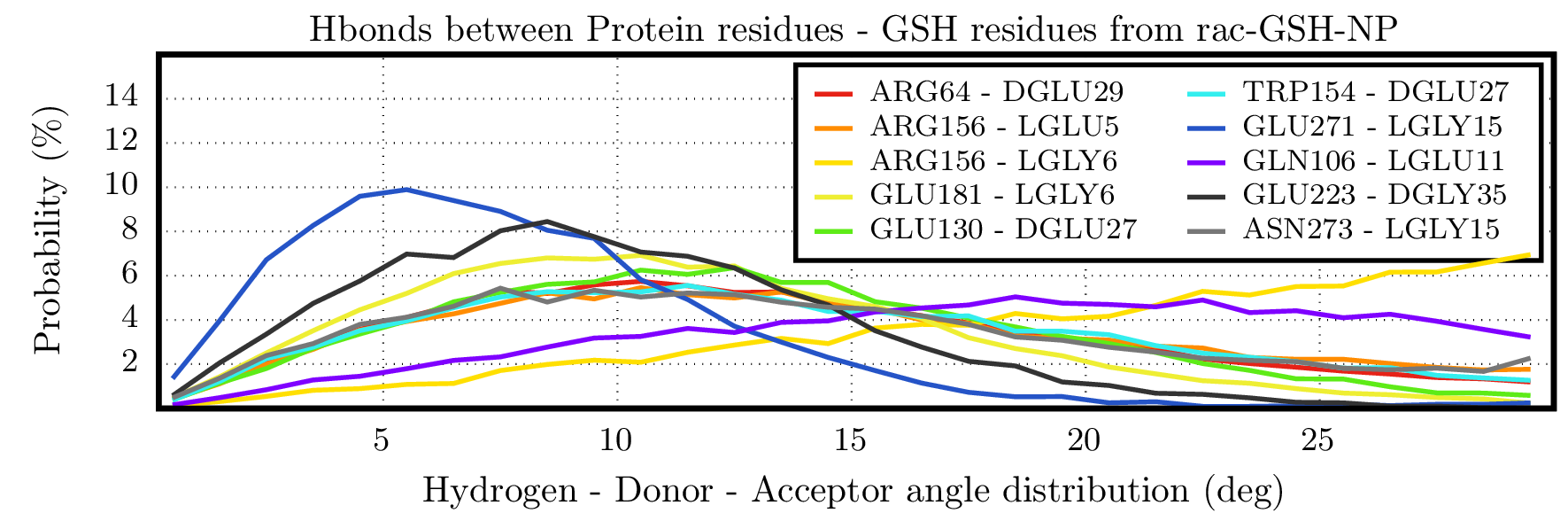

**Figure S26.** Hydrogen-Donor-Acceptor (H-D-A) angle distribution for the protein residues, which presented the highest hydrogen bond occupancy values.

**Comments:** In addition to assessing the relative thermodynamic stability of hydrogen bonds in terms of occupancy, an analysis of the geometric features of each hydrogen bond revealed that anionic residues from the protein had shorter donor-acceptor (D-A) distances (**Figure S25**) and hydrogen-donor-acceptor (H-D-A) angles closer to the ideal collinear arrangement (**Figure S26**), compared to cationic, polar, and nonpolar residues.

In this case, although the *rac*-GSH-NP complex had a greater number of hydrogen bonds, only 3 out of 10 were shorter and collinear. In contrast, the *D*-GSH-NP complex had 6 out of 7 stronger hydrogen bonds, while the *L*-GSH-NP complex had only 5 out of 9 meeting these criteria (**Figure S27-S29**).

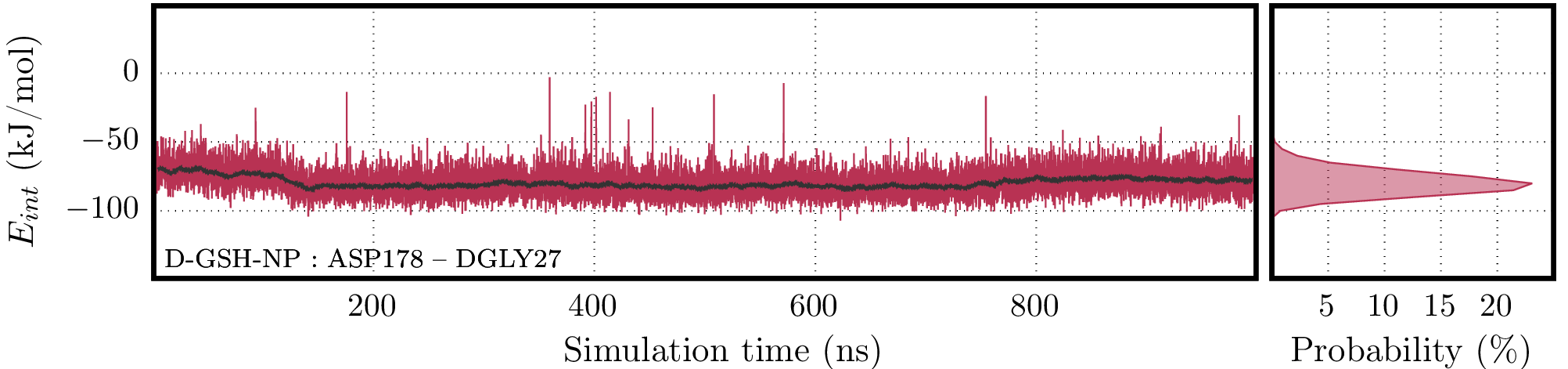

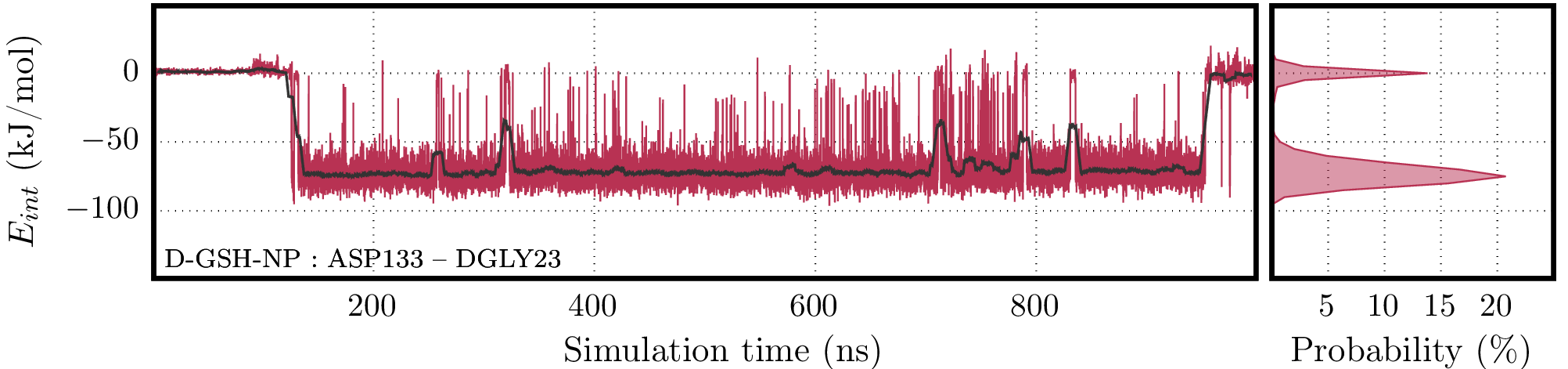

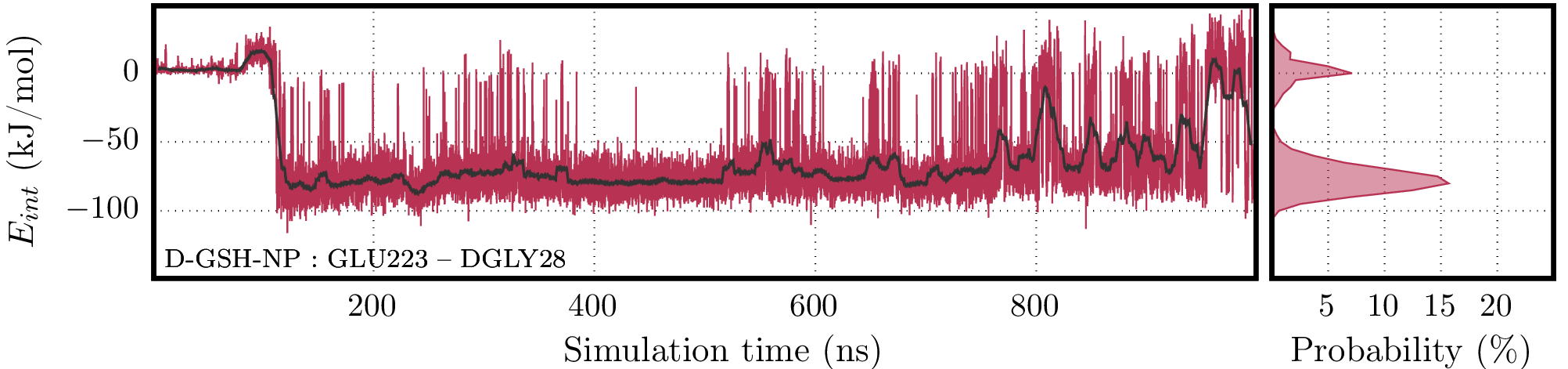

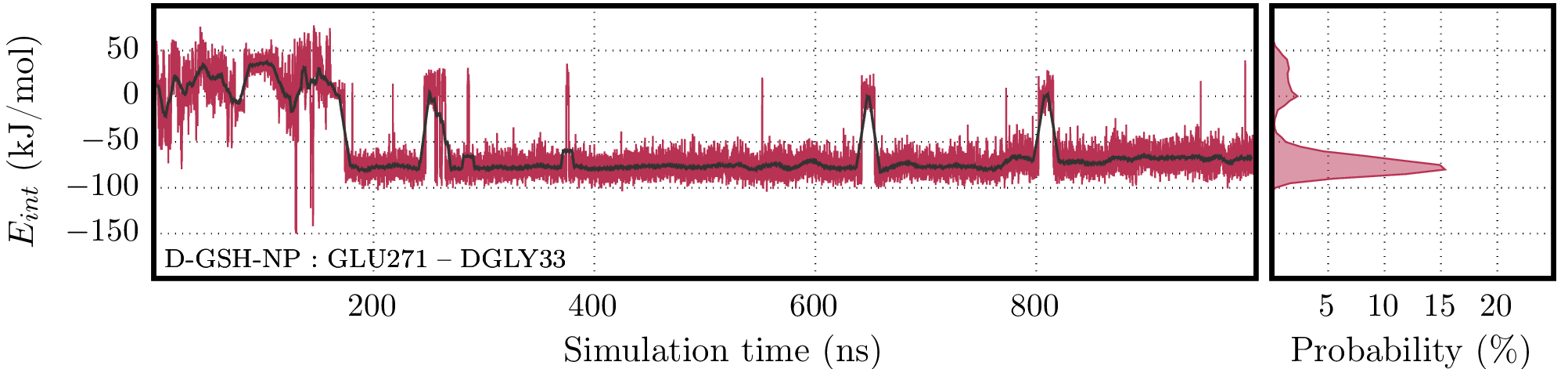

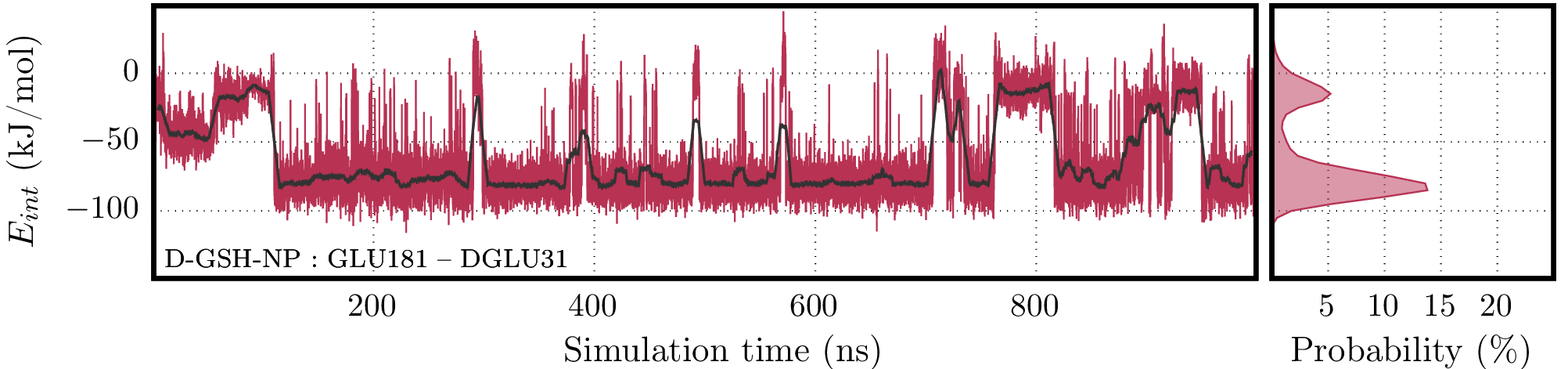

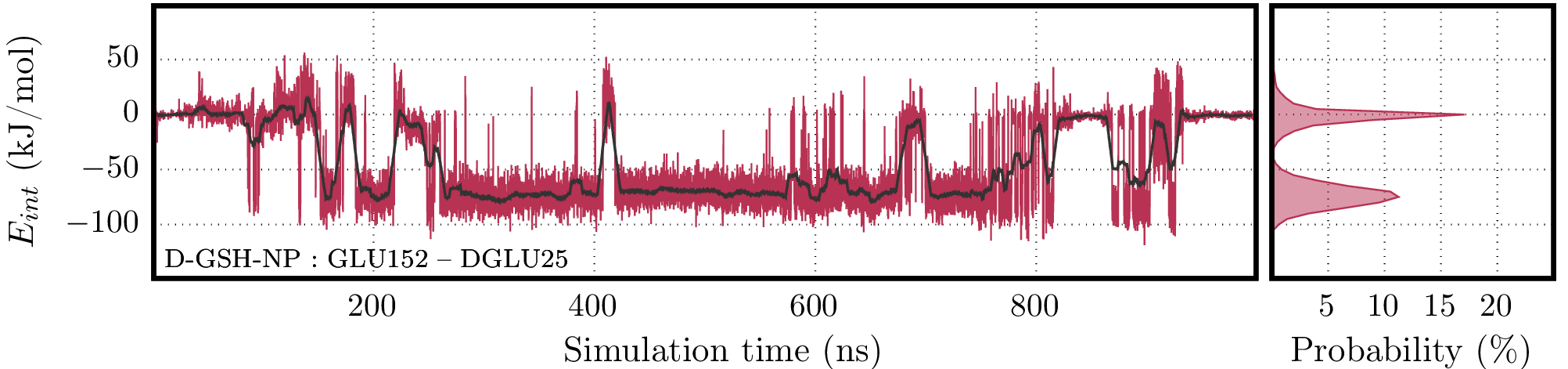

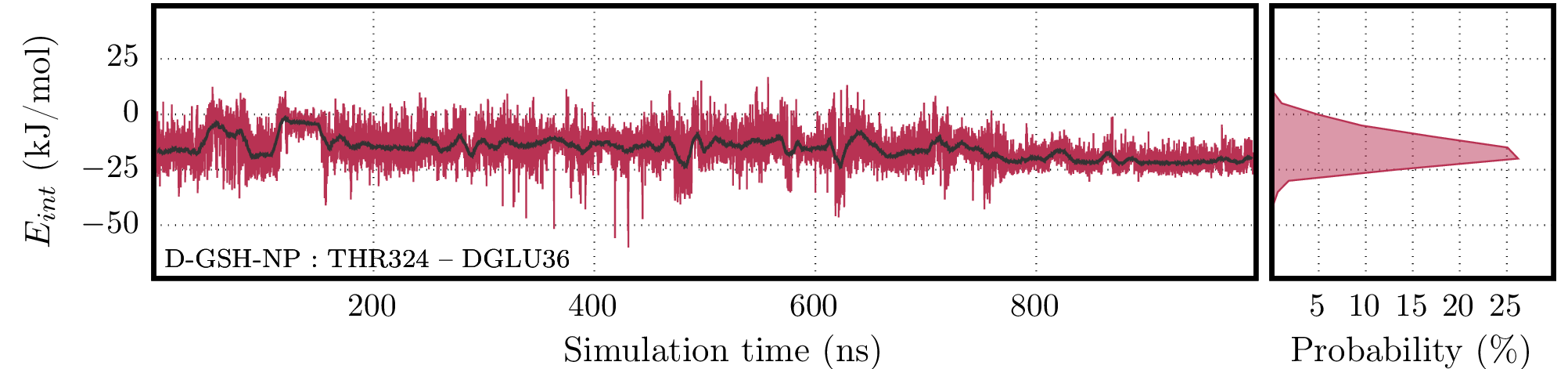

**Figure S27.** Interaction energy values for the interaction between *D*-GSH-NP and each one of the residues which presented the highest occupancy values in the hydrogen bond analysis. For each residue, the time series contains the raw data (red line) and its running average (black line) along with the histogram on the right panel.

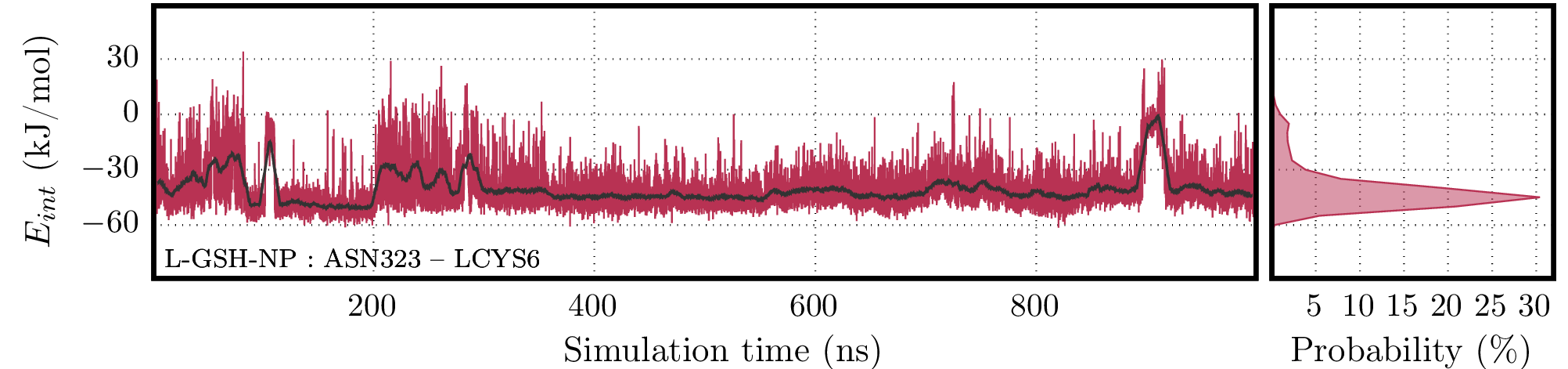

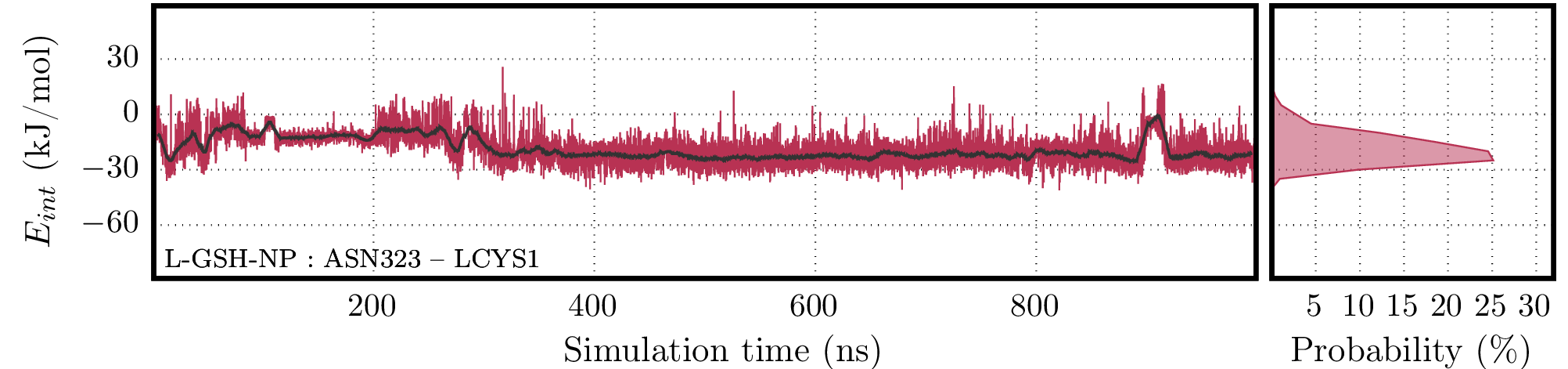

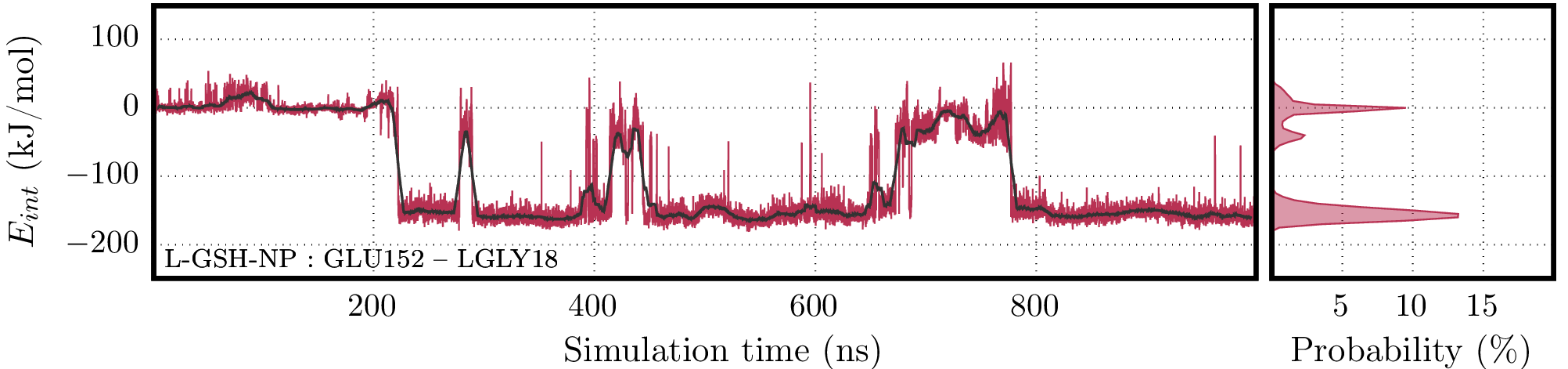

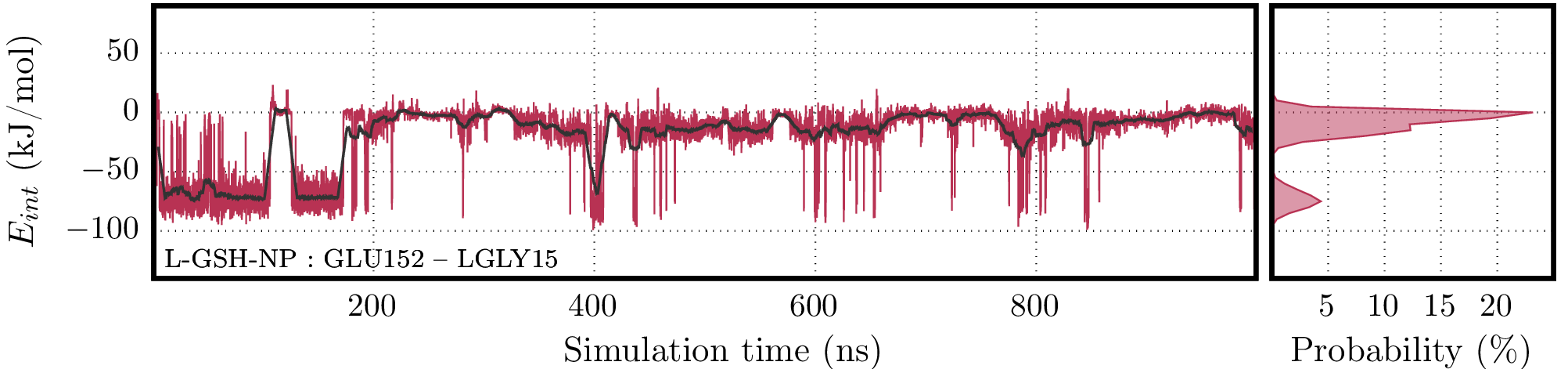

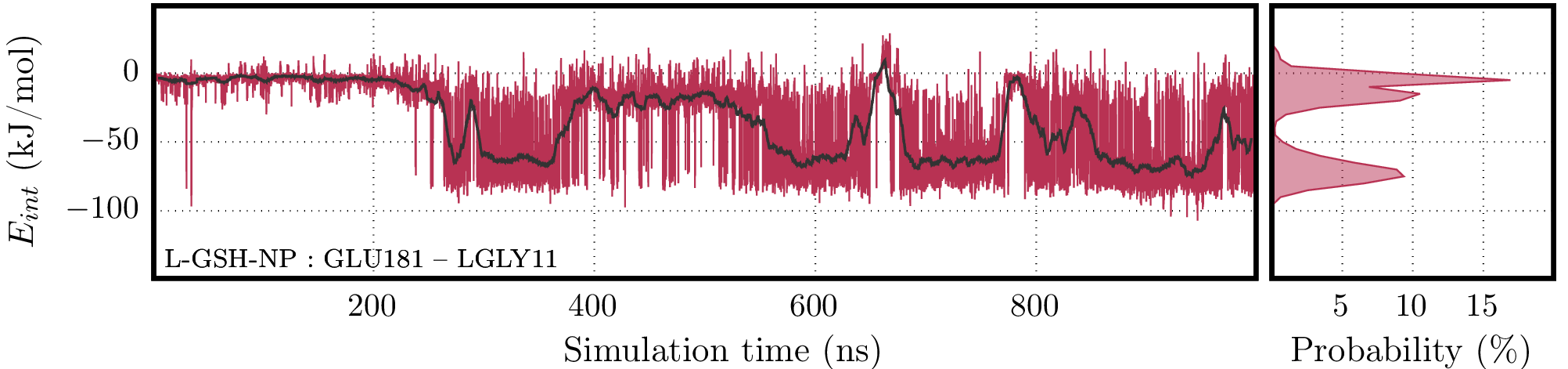

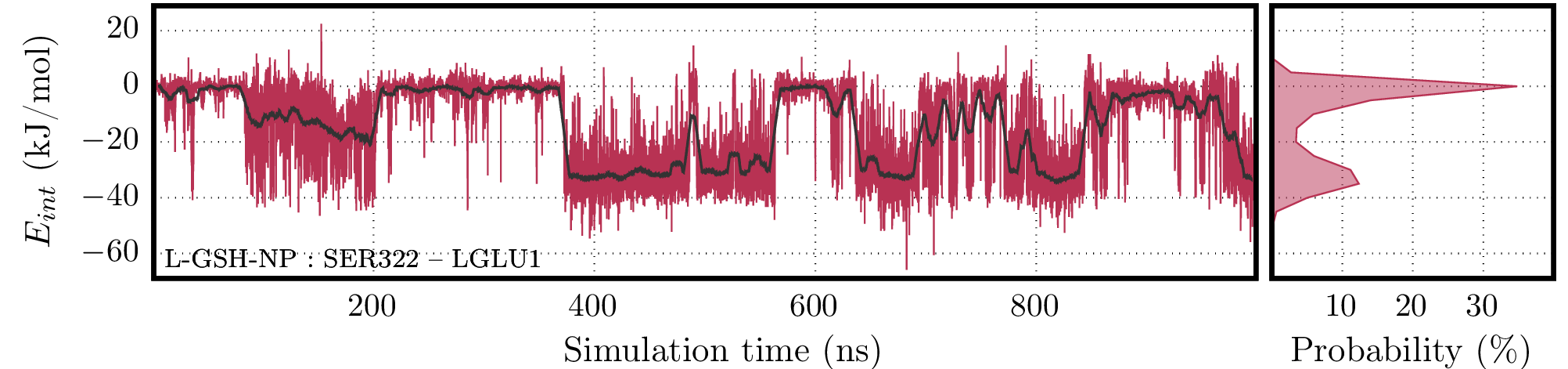

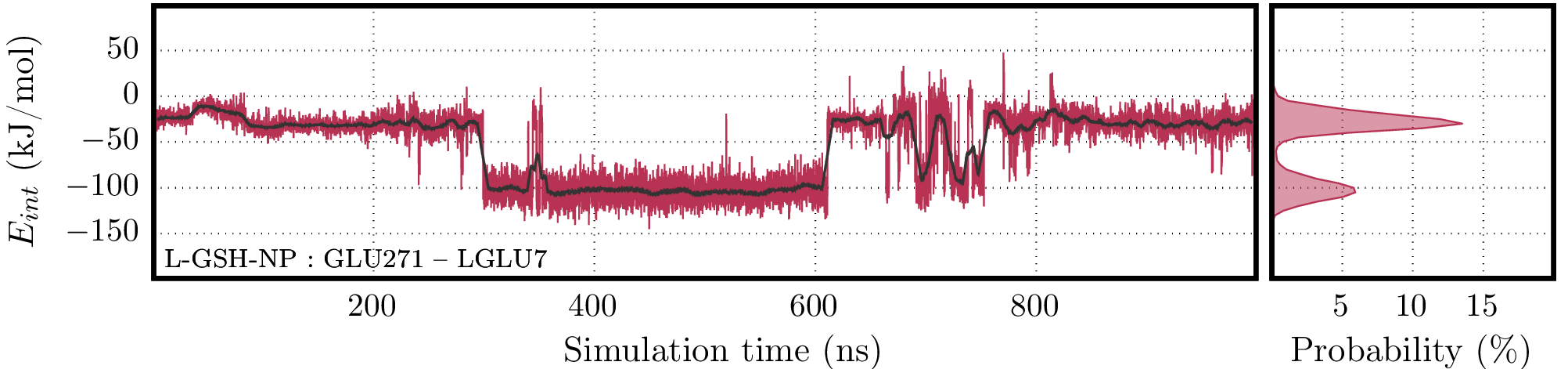

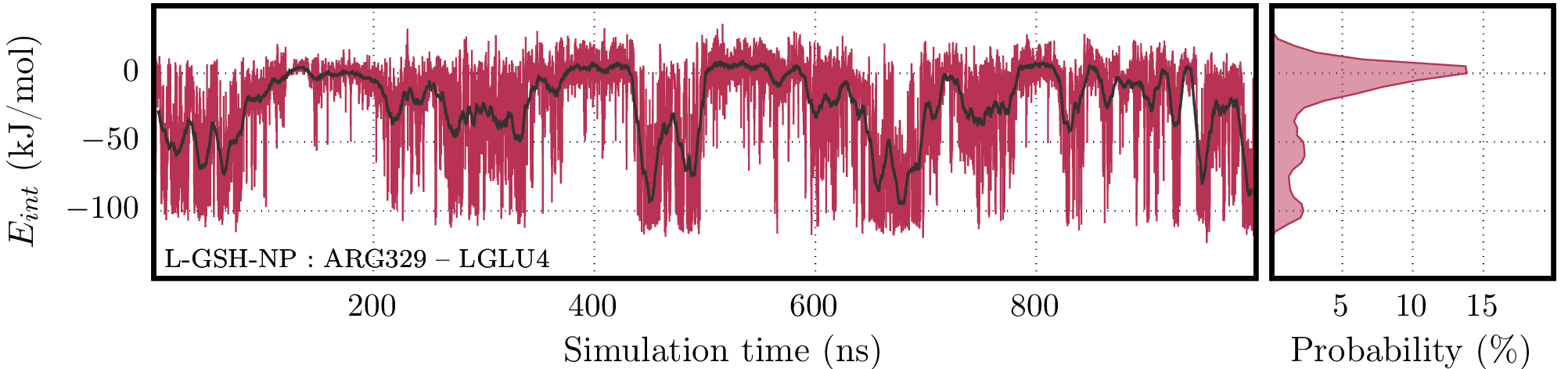

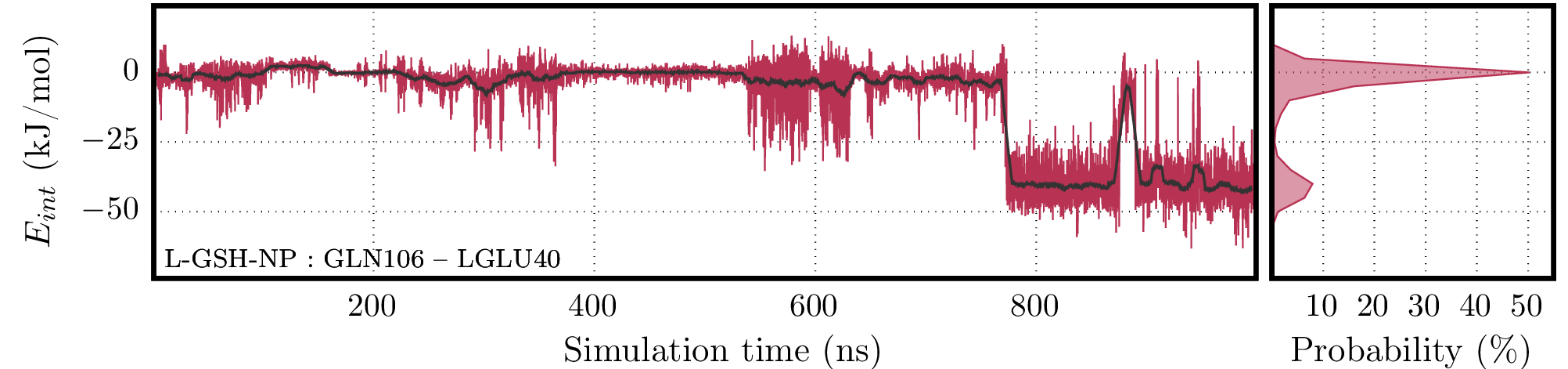

**Figure S28.** Interaction energy values for the interaction between *L*-GSH-NP and each one of the residues which presented the highest occupancy values in the hydrogen bond analysis. For each residue, the time series contains the raw data (red line) and its running average (black line) along with the histogram on the right panel.

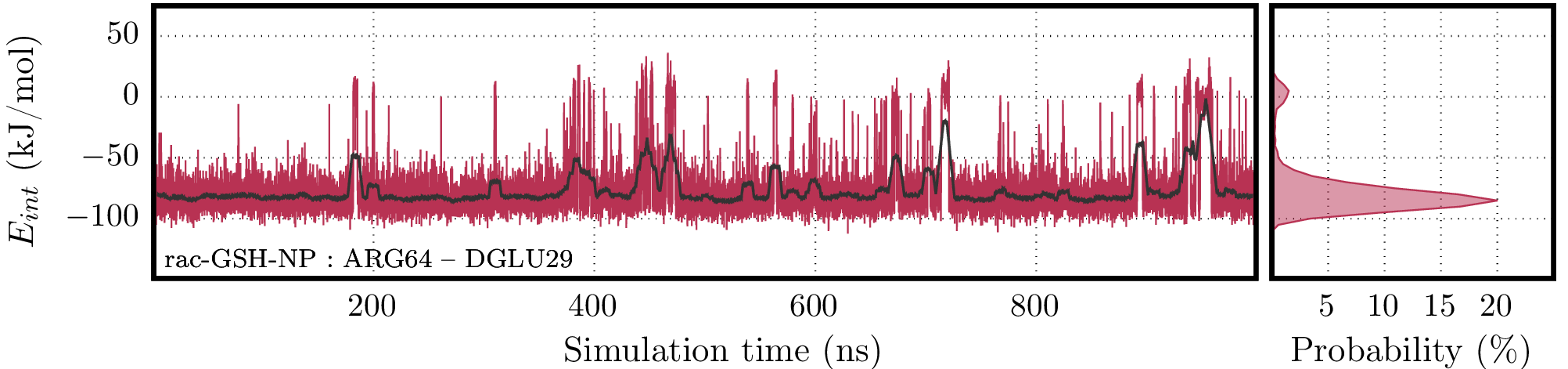

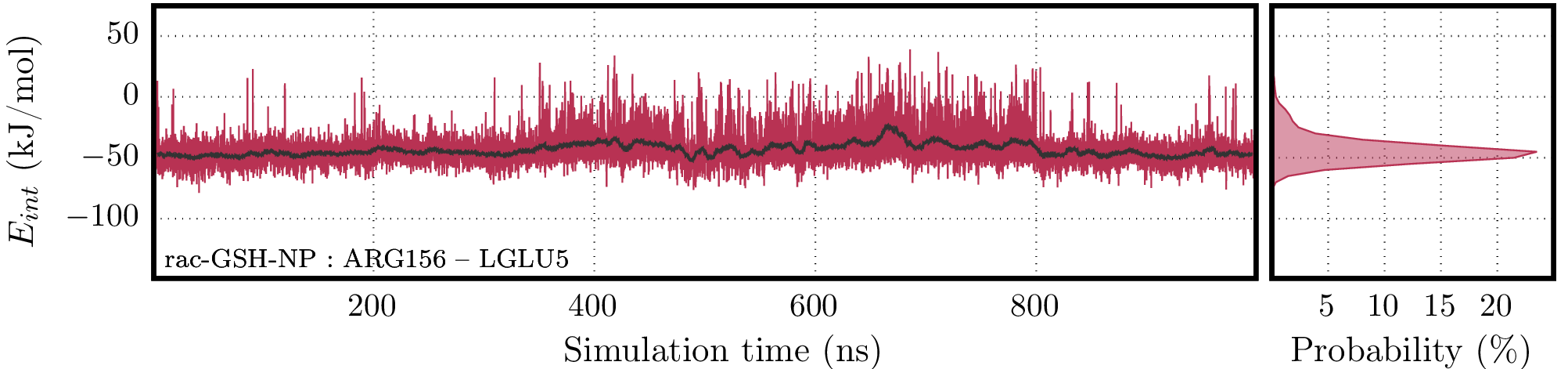

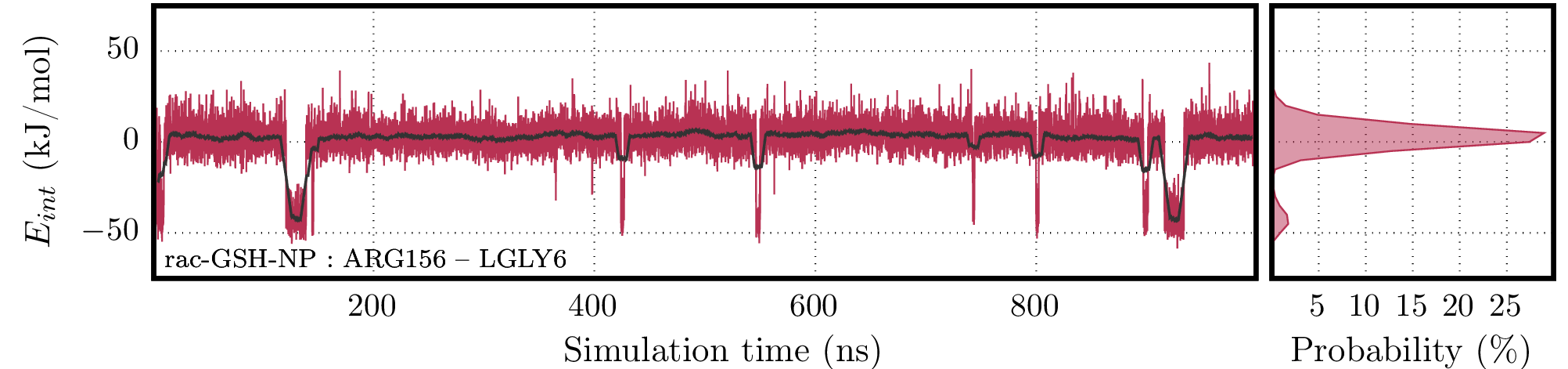

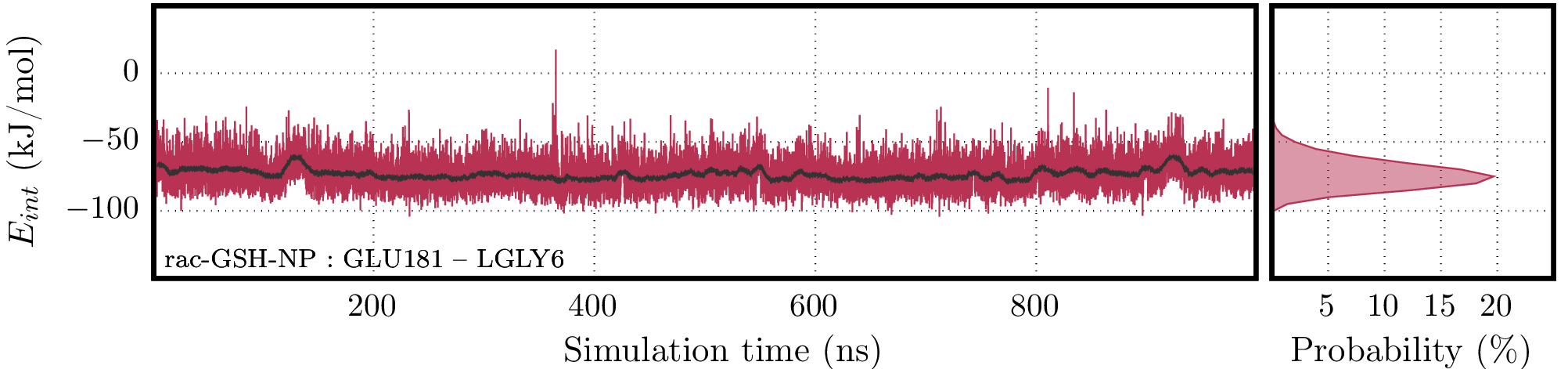

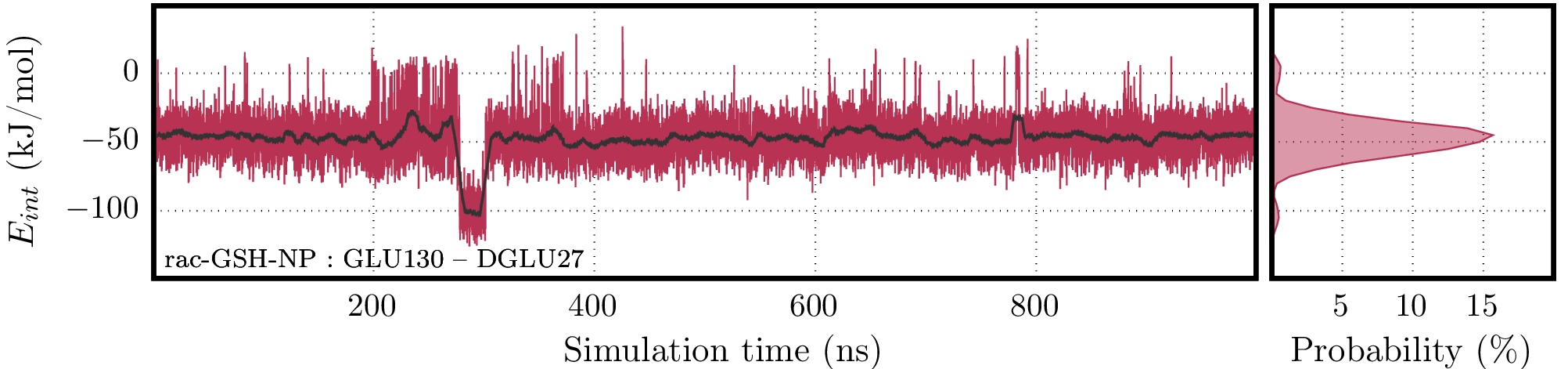

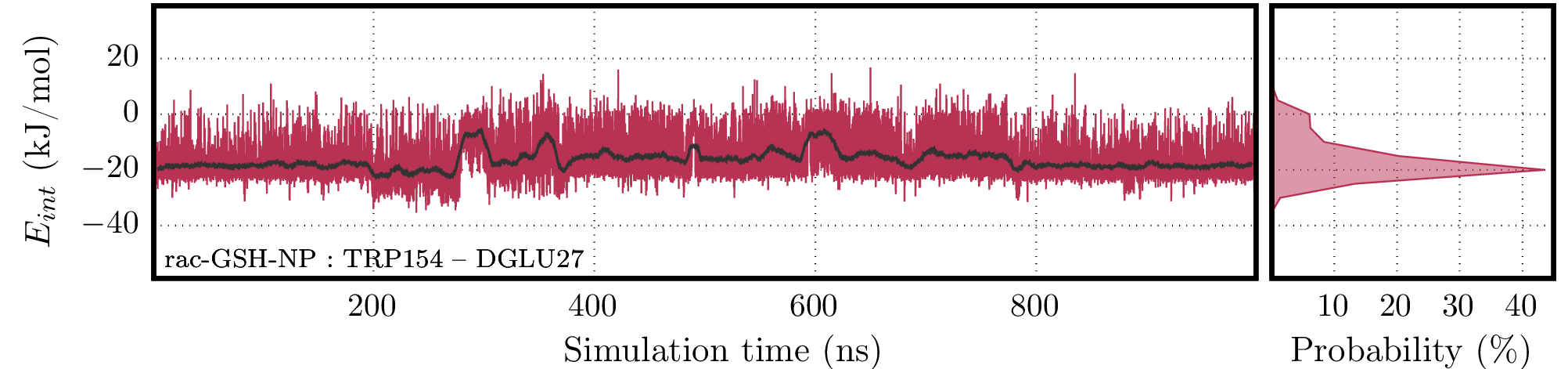

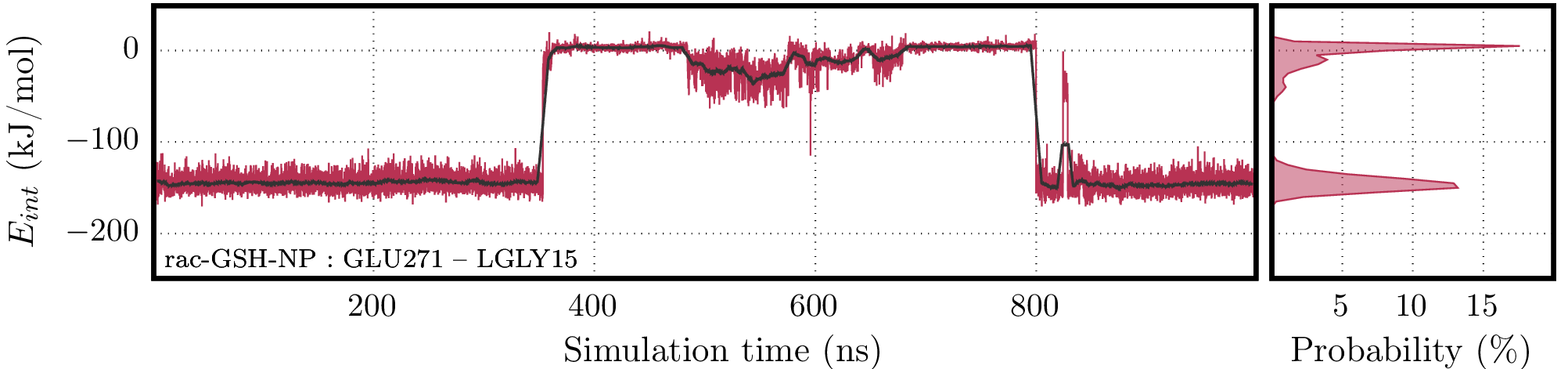

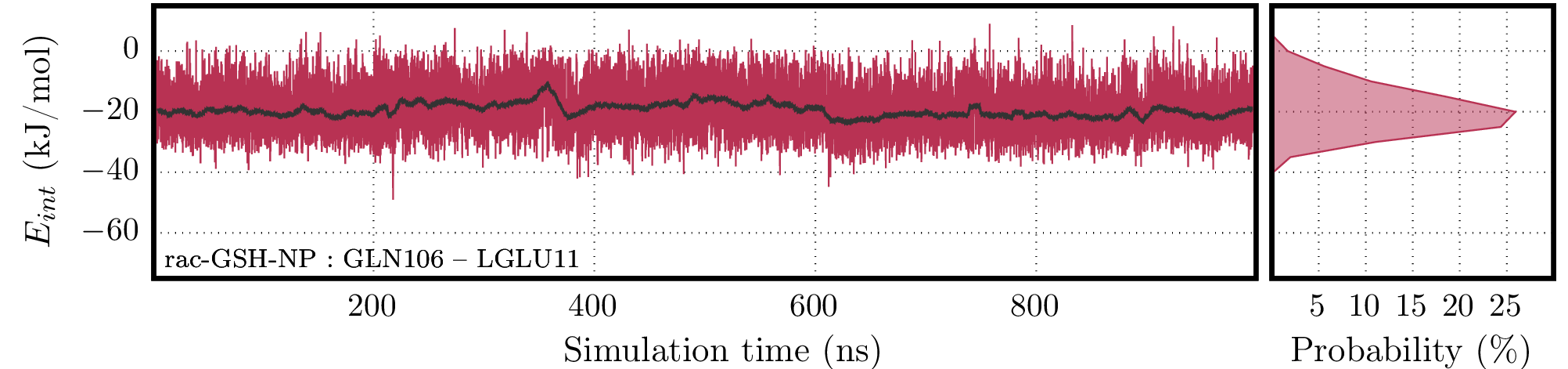

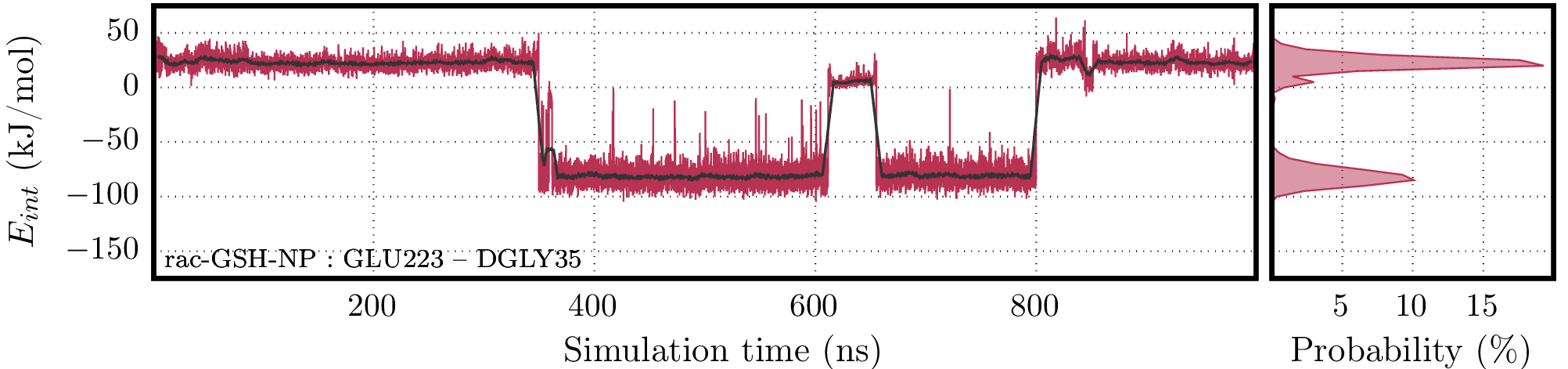

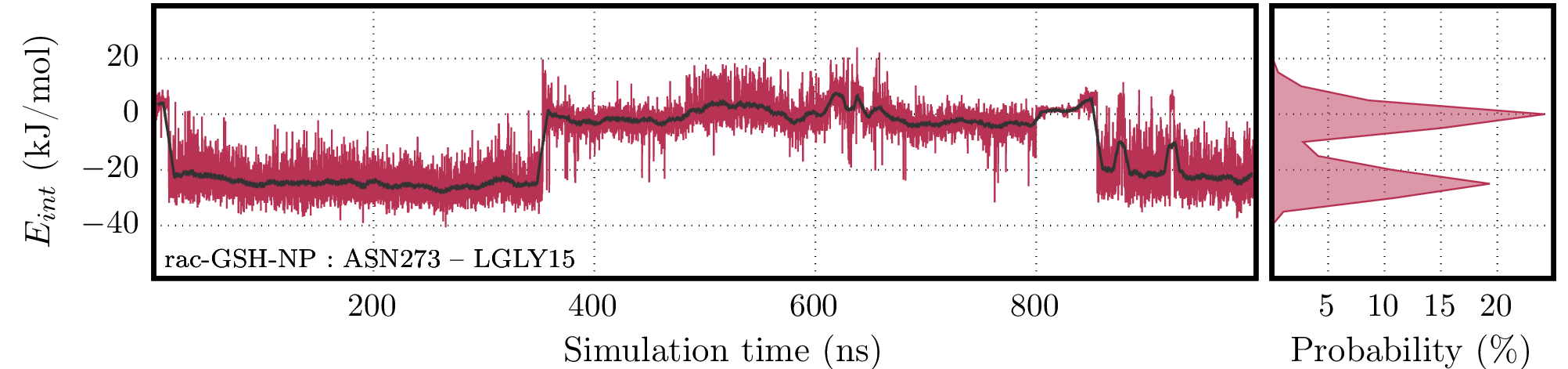

**Figure S29.** Interaction energy values for the interaction between *rac*-GSH-NP and each one of the residues which presented the highest occupancy values in the hydrogen bond analysis. For each residue, the time series contains the raw data (red line) and its running average (black line) along with the histogram on the right panel.

**Comments:** Although hydrogen bonds play a central role in the GSH-NP interactions with the protein, we must acknowledge that when GSH molecules on the NP surface form hydrogen bonds with the amino acids on the surface of the protein there are other interactions between these close lying molecules, such as dipole-dipole interactions and van der Waals interactions, all of which contribute to the stability of each hydrogen bond and to the overall stability of the complexes. The total interaction energies for the residues is computed as the sum of Coulomb and Lennard-Jones contributions (**Figure S27-S29**) and present some trends: (1) nonpolar (TRP) and polar residues (THR, ASN, GLN and SER) have weaker interactions as compared to charged residues (ASP, GLU and ARG); (2) *D*-GSH and *rac*-GSH had more stable interactions than *L*-GSH; and (3) *D*-GSH has fewer interactions than *rac*-GSH, but the interactions of *rac*-GSH were weaker on average and some of them presented repulsive plateaus, unlike the *D*-GSH interactions, which remained attractive throughout the trajectory.

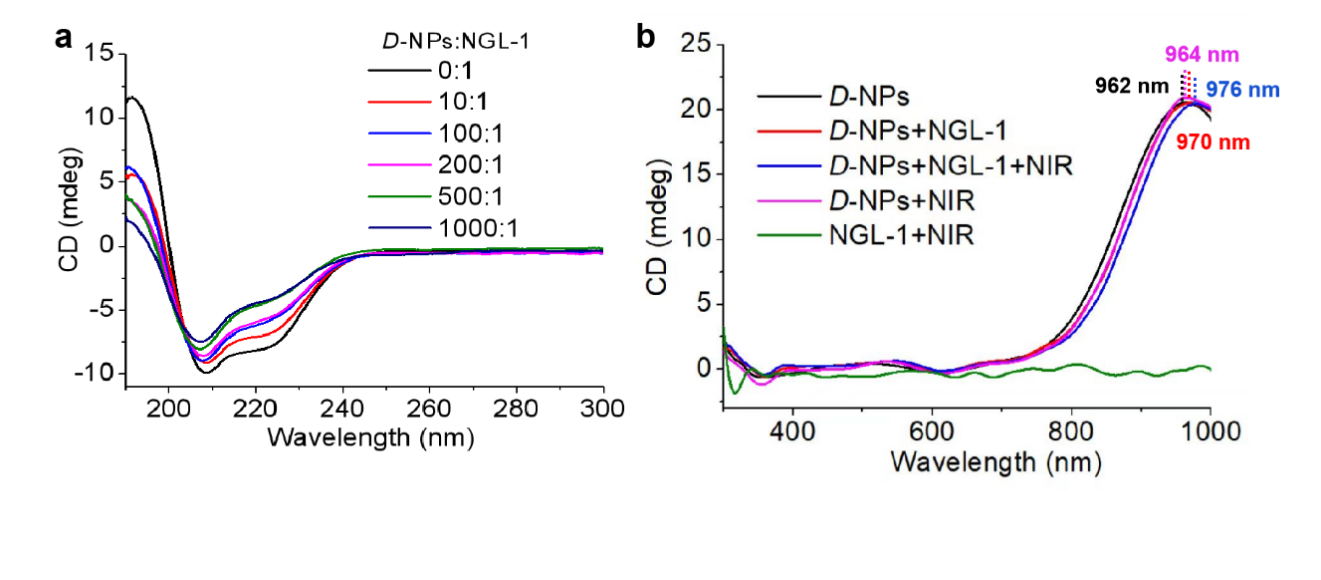

**Figure S30. (a)** CD spectra from 190 nm to 300 nm of *D*-NPs incubated with NGL-1 at different concentration ratios. **(b)** CD spectra from 300 nm to 1000 nm of *D*-NPs incubated with NGL-1 at a concentration ratio of 500:1 with or without NIR illumination (980 nm, 400 mW/cm^2^, 10 min).

**Comments:** The CD spectra of *D*-NPs and *L*-NPs displayed mirror-symmetry in the ultraviolet region from 190 nm to 300 nm **(Figure S30a)**; *rac-*NPs were CD silent (not shown). Moreover, NGL-1 displayed a stronger change of conformation when incubated with *D*-NPs at a concentration ratio of 500:1 (*D*-NPs: NGL-1) **(Figure S30b)**.

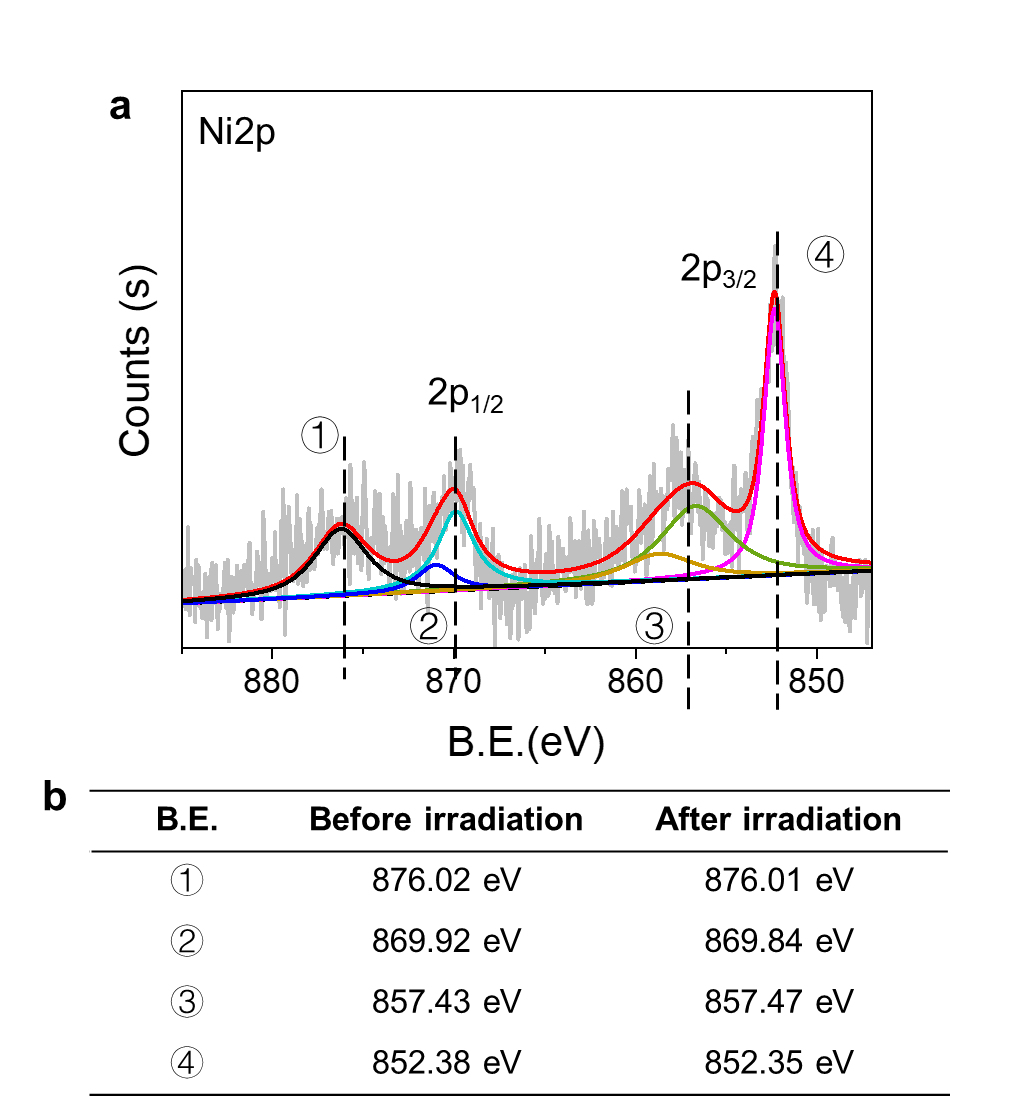

**Figure S31.** **(a)** XPS spectra of *D*-NPs after NIR illumination (980 nm laser, 400 mW/cm^2^, 10 min). **(b)** The binding energy of *D*-NPs before and after NIR illumination.

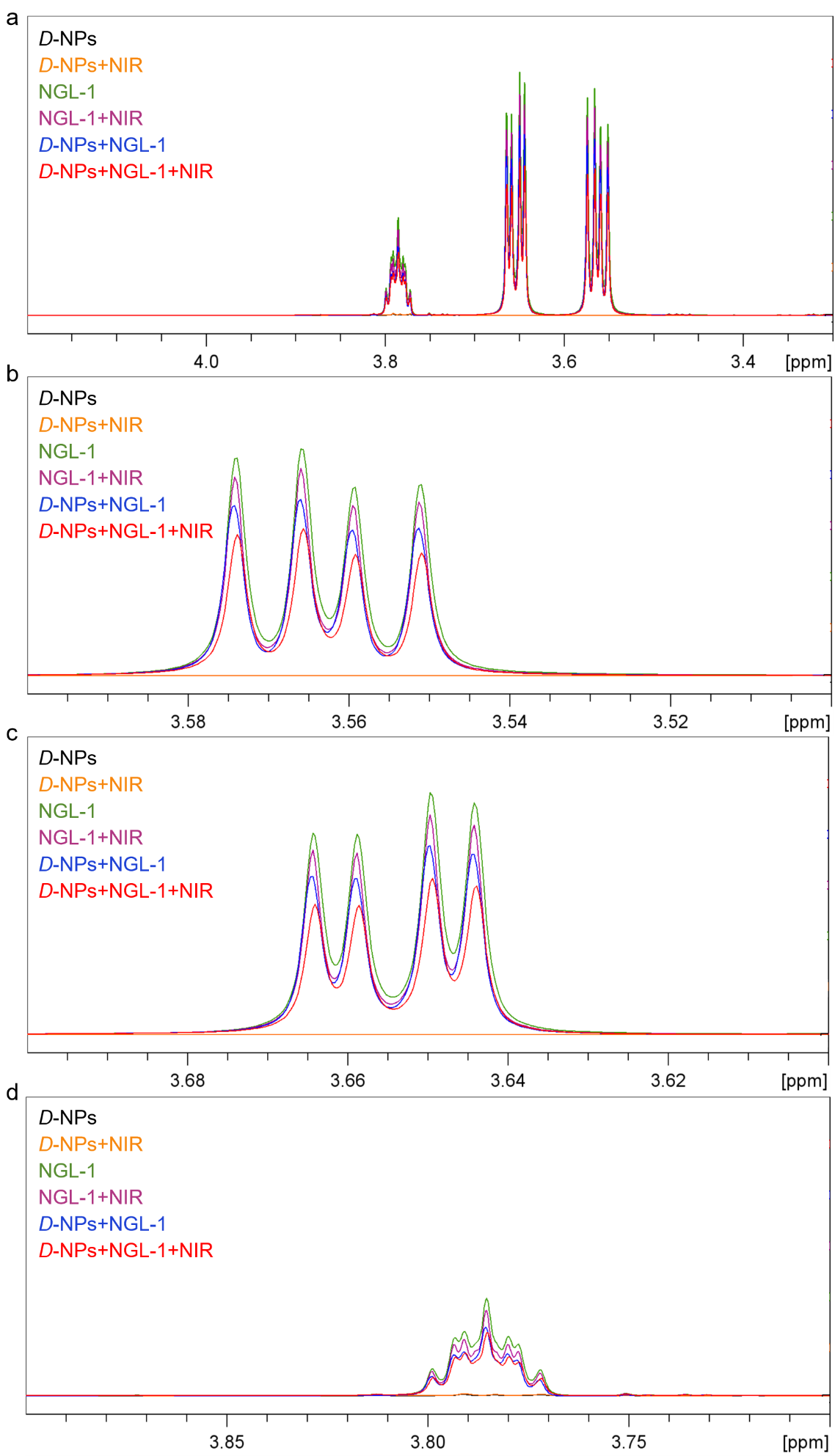

**Figure S32.** **(a)** ^1^H-NMR spectra in the region of 4.2-3.3 ppm of *D*-NPs, *D*-NPs+NIR, NGL-1, NGL-1+NIR, and *D*-NPs incubated with NGL-1 at a concentration ratio of 100:1 without or with NIR illumination (980 nm, 400 mW/cm^2^, 10 min) obtained using the Carr-Purcell-Meiboom-Gill (CPMG) sequence. **(b-d)** Zoom the ^1^H-NMR spectra of *D*-NPs, *D*-NPs+NIR, NGL-1, NGL-1+NIR, *D*-NPs incubated with NGL-1 at a concentration ratio of 500:1 without or with NIR illumination in the region of **(b)** 3.5-3.6 ppm, **(c)** 3.6-3.7 ppm, and **(d)** 3.7-3.9 ppm, respectively.

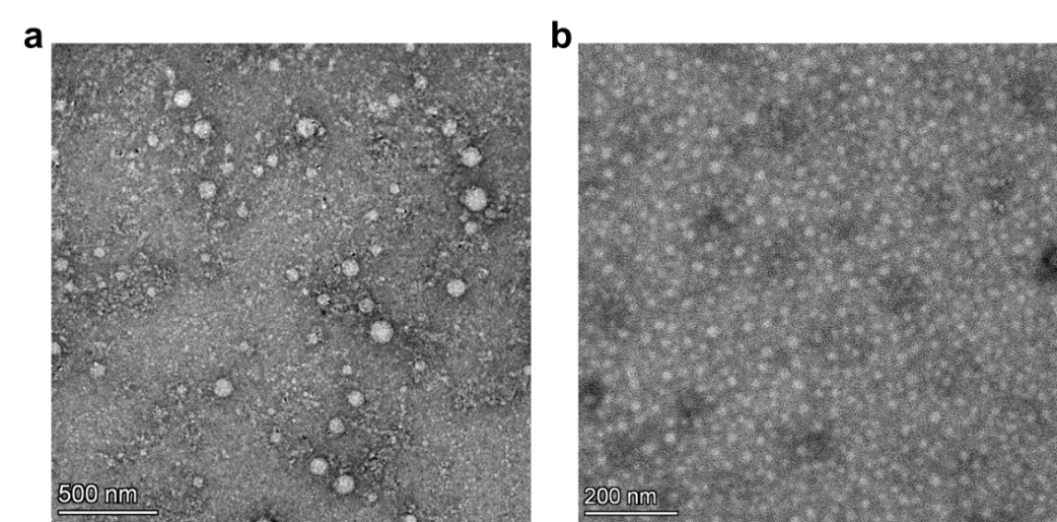

**Figure S33.** **(a)** Negatively stained TEM images of *D*-NPs incubated with NGL-1 after NIR illumination (980 nm, 400 mW/cm^2^, 10 min). **(b)** Negatively stained TEM images of *D*-NPs incubated with NGL-1 before NIR illumination (980 nm, 400 mW/cm^2^, 10 min). Strong agglomeration of NPs with proteins after NIR illumination is observed.

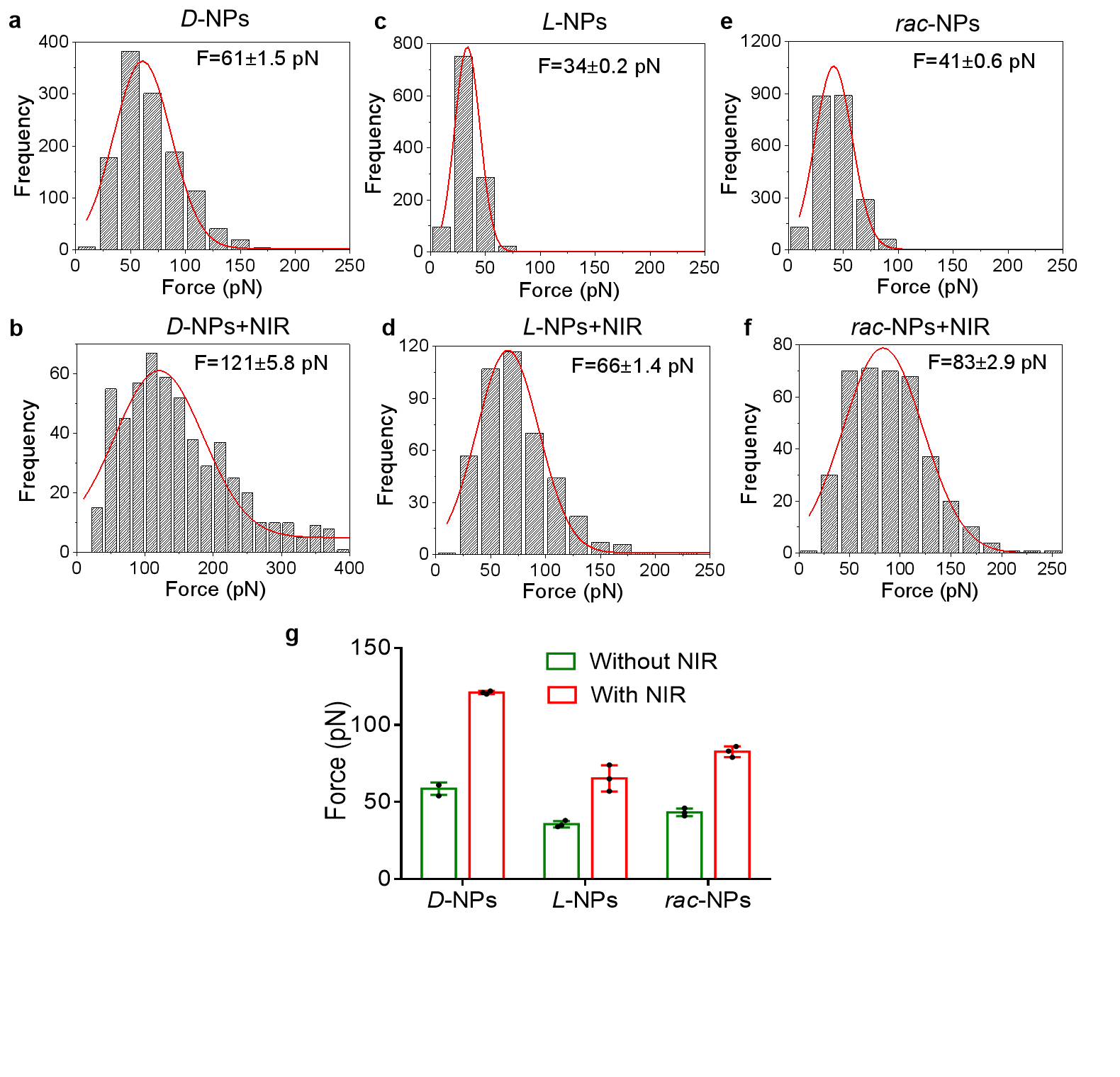

**Figure S34.** **(a-f)** The interaction force between *D*-, *L*-, or *rac*-NPs and neurons cells with **(a-c)** or without **(d-f)** NIR illumination (980 nm laser; 400 mw/cm^2^) measured by atomic force microscope (AFM). **(g)** The interaction force between *D*-, *L*-, or *rac*-NPs and neurons cells with or without NIR illumination. Data are presented as mean ± s.d. (n = 3).

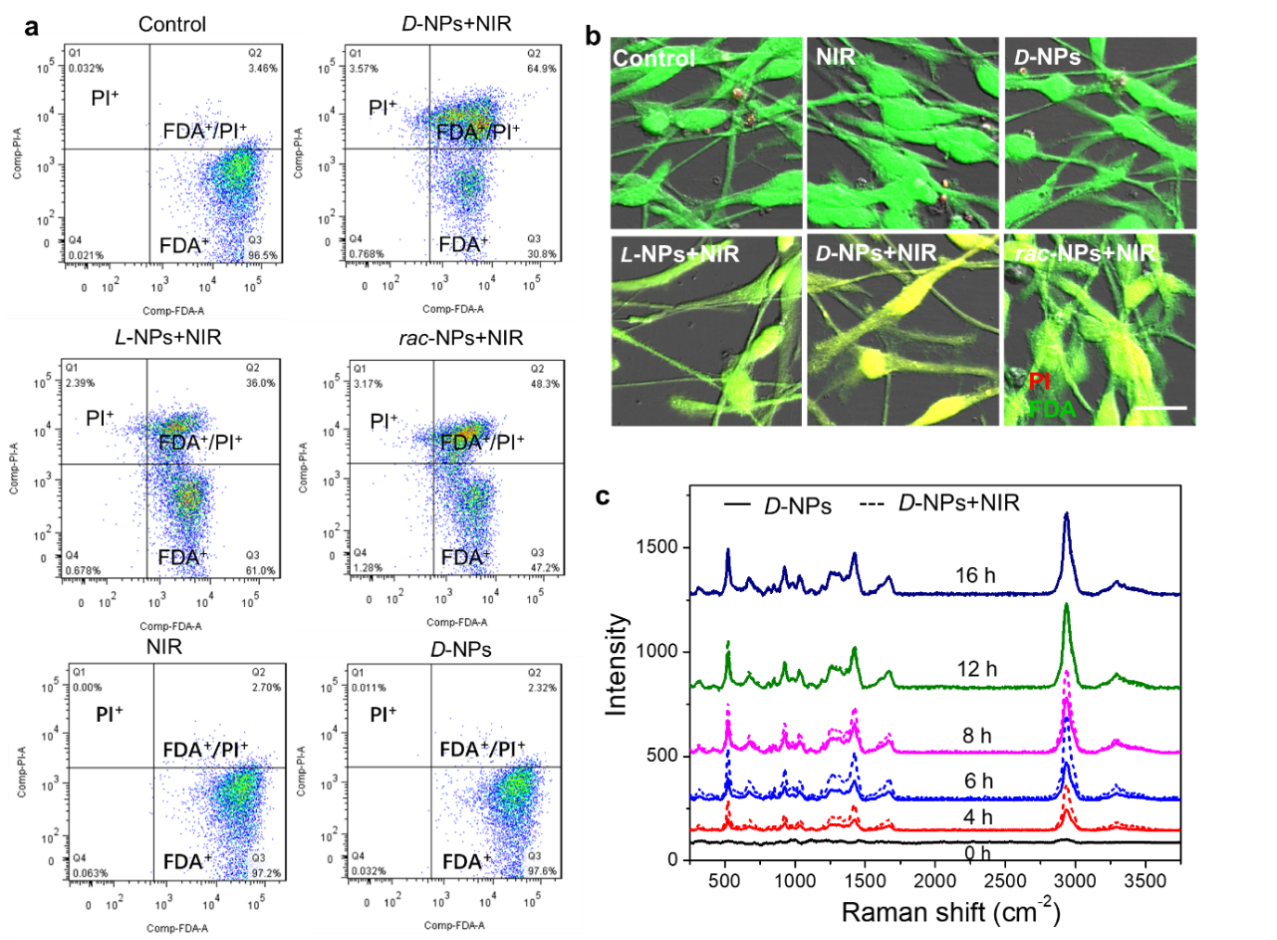

**Figure S35. (a)** Flow cytometric measurement of hippocampal neurons stained with propidium iodide (PI) and fluorescein diacetate (FDA). The cells incubation with *D*-/*L*-/*rac*-NPs under NIR light illumination (980 nm laser, 400 mW/cm^2^), or only incubation with *D*-NPs without NIR illumination, or only under NIR illumination without *D*-NPs incubation, and subsequently stained with PI and FDA, control represents hippocampal neurons cells without NPs and NIR treatment. FDA^+^, FDA^+^/PI^+^, and PI^+^ denotes the subpopulations of unpermeabilized cells, permeabilized-resealed cells, and dead cells, respectively. **(b)** Confocal imaging of hippocampal neurons stained with PI and FDA. The cells incubation with *D*-/*L*-/*rac*-NPs and under NIR light illumination (980 nm laser, 400 mW/cm^2^), or only incubation with *D*-NPs, or only with NIR light treatment and subsequently stained with PI and FDA. The images show unpermeabilized cells (green), permeabilized-resealed cells (green+red), and dead cells (red). **(c)** Raman spectra of hippocampal neurons incubated with *D*-NPs with or without NIR illumination (980 nm laser, 400 mW/cm^2^).

**Comments:** Both flow cytometry and confocal microscopy imaging indicate that chiral NPs enhance cell membranes permeability under NIR stimulation, leading to an increased rate of NPs entry into the neurons, which was confirmed by the Raman scattering spectra.

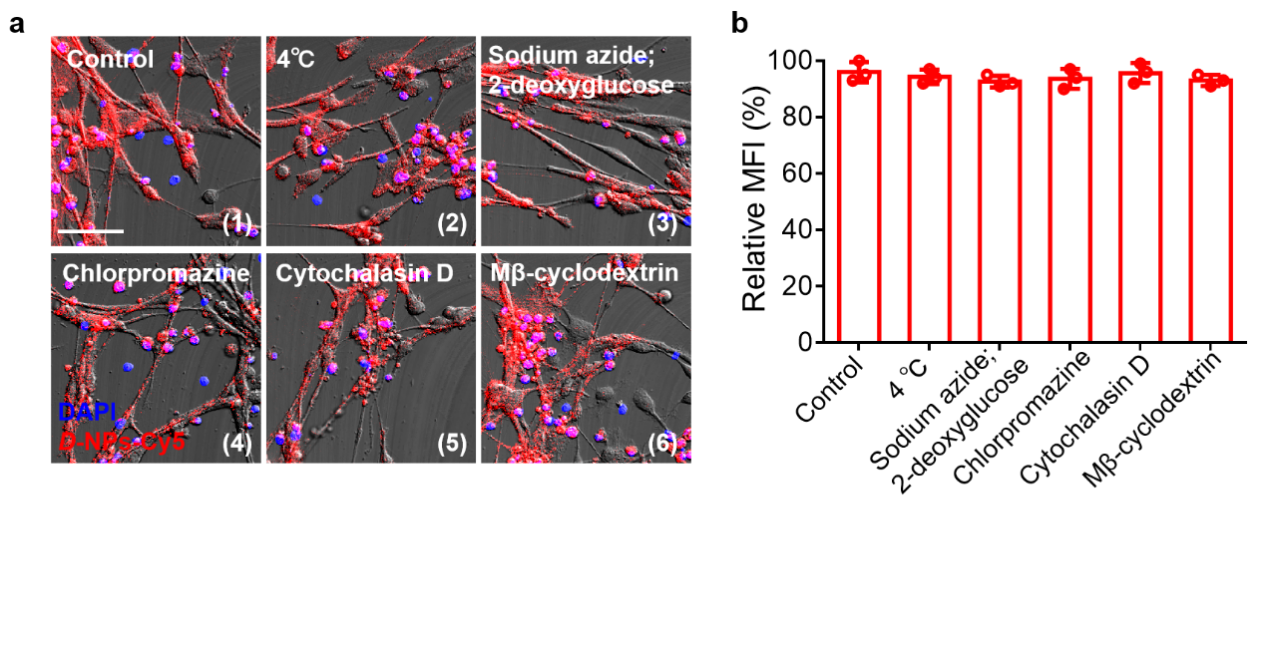

**Figure S36. (a)** Confocal images of hippocampal neurons after (1) incubation in PBS; (2) incubation at 4 ℃; (3) incubation with sodium azide and 2-deoxyglucose (block active internalization); (4) incubation with chlorpromazine (clathrin inhibitor); (5) incubation with cytochalasin D (phagocytosis inhibitor) and (6) incubation with Mβ-cyclodextrin (caveolae inhibitor); red: *D*-NPs-Cy5; blue: DAPI for nucleus; scale bar, 50 μm. **(b)** Relative fluorescence intensity of *D*-NPs-Cy5 incubated with hippocampal neurons for 12 h after specified treatments. Data are presented as mean ± s.d. (n = 3).

**Comments:** We found that the entry efficiency of D-NPs with NIR light was not affected when hippocampal neurons were treated with low temperature, clathrin inhibitor, caveolae inhibitor, active internalization inhibitor, or phagocytosis inhibitor, which may be due to non-disruptive fusion of NPs with cell membranes and subsequent pass through the bilayer by passive diffusion penetration.

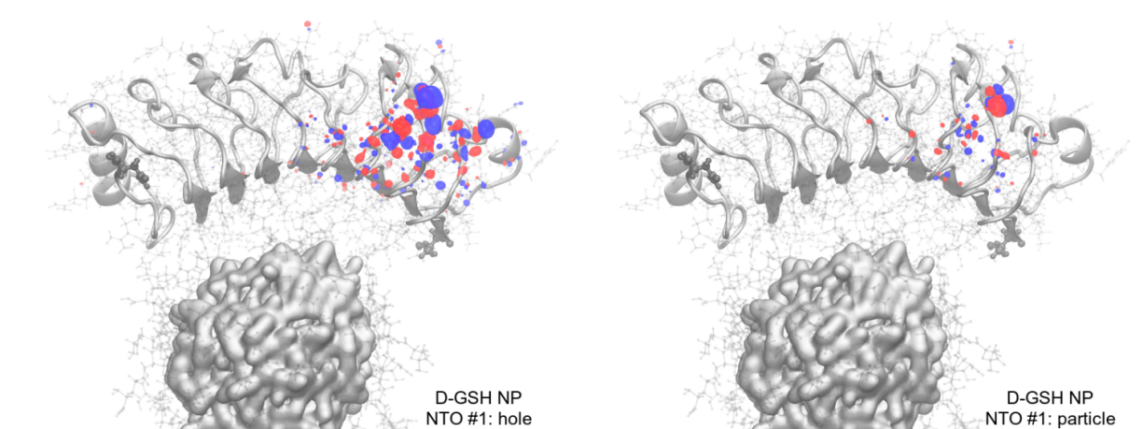

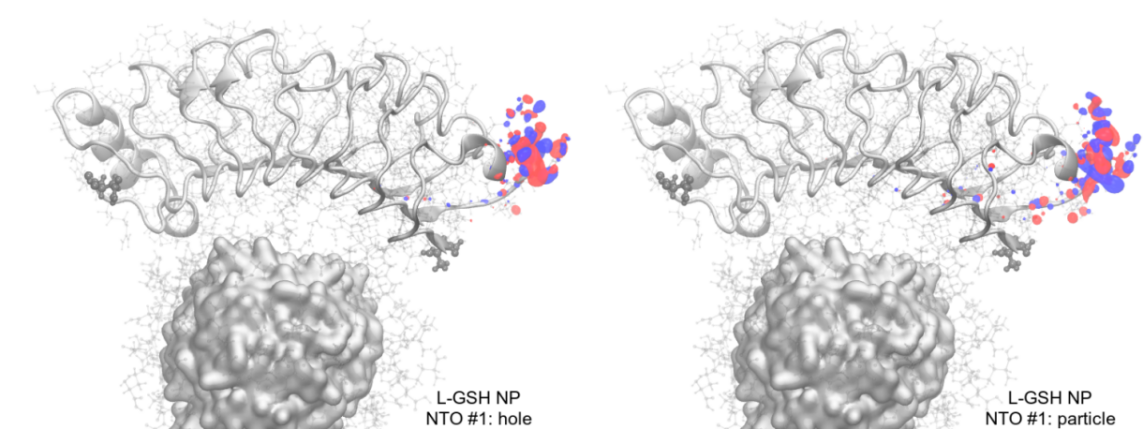

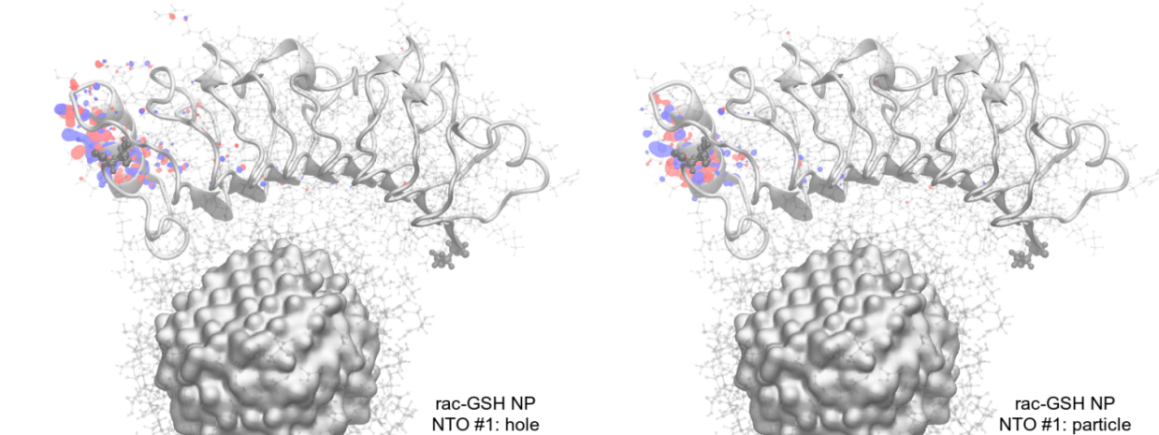

**Figure S37.** Natural Transition Orbitals (NTOs) for the lowest-energy transition obtained for *D*-GSH NP (top), *L*-GSH NP (center) and *rac*-GSH NP (bottom) complexes with a representative protein fragment. NP core is represented as a grey surface and GSH ligands are represented as grey molecular structures around it. Secondary structure of the protein fragment is represented by a grey cartoon with the residues also represented by grey atomic structures. Terminal residues (T128 and N331) are highlighted as the larger gray spheres. All NTOs were rendered considering an isovalue of 0.01 electron/Å³.

**Comments:** The protein fragment on its own is optically active only in the UV region (**Table S5**), as expected for any protein without chromophore prosthetic groups, while all complexes formed with *D*-GSH, *L*-GSH and *rac*-GSH NPs are active in the NIR region. We would expect the NPs to act as antennas, just like chromophore prosthetic groups, but inspection of the NTOs for the first 10 lowest-lying transitions shows that there is no charge transfer between NPs and proteins, all charge carriers are located exclusively on the protein (**Figure S37**). This finding is highly unexpected, since the protein molecular orbitals should never be active on this spectral window, as demonstrated by the calculation of the protein fragment without the NP (**Table S5**) and nonetheless the transition occurs in the NIR region if the NPs are present, even though the NPs are 2-3 nm away from the regions where NTOs were observed.

It is interesting to note that particle and hole were always very close to each other (**Figure S37**), indicating that these excitations are local polarizations rather than charge transfer processes. It also means that recombination is expected to take place readily. For *D*-GSH NP, charge carriers were located after the end of the alpha-helix (residues L136, T137) and along the first strand of the beta-sheet (residues P159 to E161) and also along the second strand of the beta-sheet (residues K183, R184). For *L*-GSH NP, transitions were located at the initial portion of the alpha-helix at the beginning of the protein fragment (V145 to L147) whereas for *rac*-GSH NP, transitions were located at different residues (between K307 and R316) of the alpha helix located at the end of the protein fragment. Besides being located at different positions along the LRR fragment, the NTOs differ in size, especially the particle NTO of *D*-GSH, which is highly localized on the terminal carboxylate of the E161 sidechain, being formed mostly by *2p* atomic orbitals of the C and the two O atoms. The degree of localization is expected to affect how the energy released upon nonradiative recombination takes place, and clearly the *D*-GSH NP will release the energy on a handful of atoms while the *L*-GSH and *rac*-GSH will spread the same amount of energy over many more atoms. The energy from the 980 nm photon used experimentally to illuminate the systems amounts to *ca.* 120 kJ/mol, which should be compared to the average thermal energy of *ca.* 4 kJ/mol per atom at 310 K. So, the photon energy split among the 3 atoms of the glutamate side chain in *D*-GSH NP means a tenfold increase in the kinetic energy of these atoms, while *L*-GSH and *rac*-GSH NPs will have the same amount of energy split over a few tens of atoms and this would increase only twofold the kinetic energy of these atoms. Thus, the differences observed for each enantiomer upon illumination are most likely related to the patterns of energy partition along the LRR fragment and not the actual amount of energy.

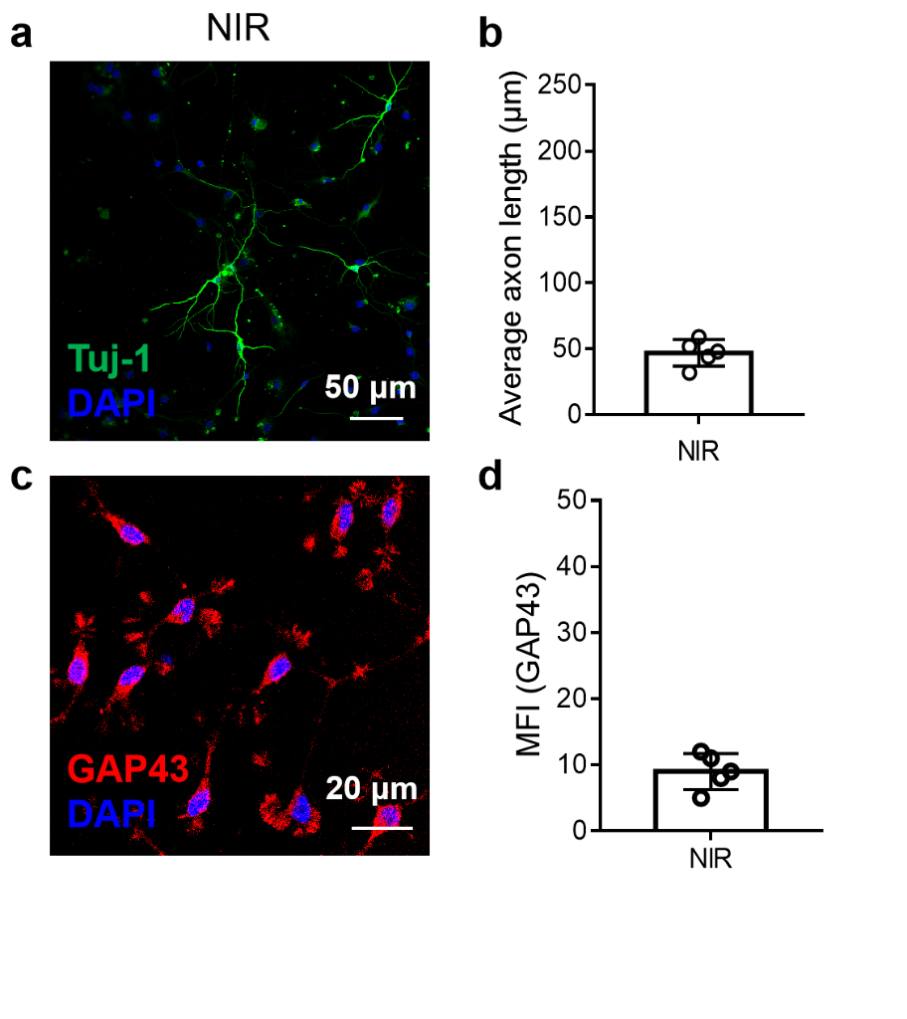

**Figure S38. (a)** Confocal images of hippocampal neurons extracted from AD mice after NIR illumination only (980 nm laser, 400 mW/cm^2^, no NPs). Tuj-1 (green) to label neurons, and DAPI (blue) to stain nuclei. **(b)** Quantification of the axonal length of hippocampal neurons extracted from AD mice after NIR illumination only (980 nm laser, 400 mW/cm^2^, no NPs) from **Figure** **S38a**. **(c)** Confocal images of hippocampal neurons extracted from AD mice only under NIR illumination (980 nm laser, 400 mW/cm^2^). GAP43 (red) to label axon, and DAPI (blue) to stain nuclei. **(d)** Mean fluorescence intensity of GAP43 in hippocampal neurons extracted from AD mice after NIR illumination only (980 nm laser, 400 mW/cm^2^, no NPs) from **Figure** **S38c**. About 10-50 cells from any region of each well were used for average axon length and fluorescence intensity statistics. Each experiment was repeated five times. Data are presented as mean ± s.d.

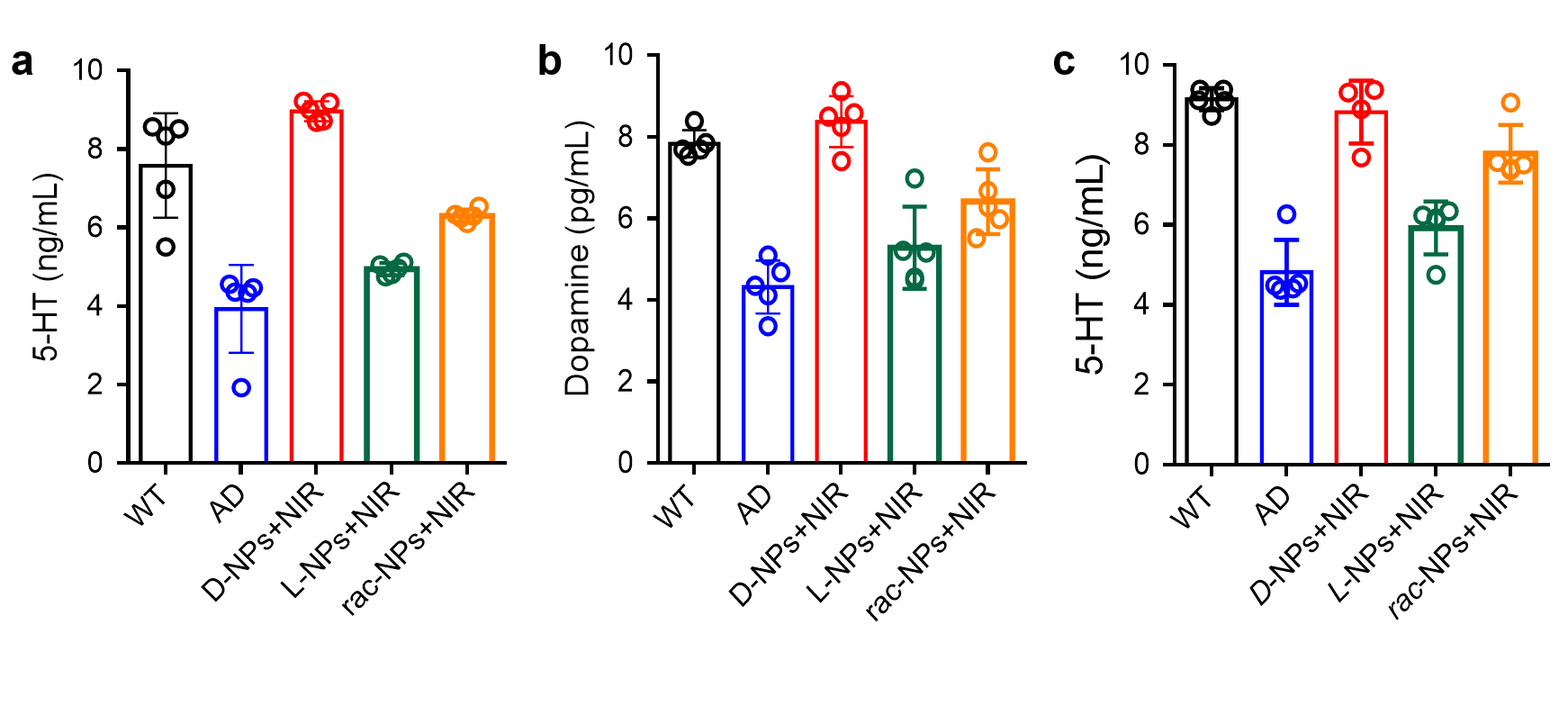

**Figure S39. (a, b)** The contents of 5-HT **(a)** and dopamine **(b)** detected by ELISA in hippocampal neurons extracted from WT or AD mice incubated with *D-*NPs, *L-*NPs, and *rac-*NPs under NIR illumination (980 nm laser, 400 mW/cm^2^). **(c)** The contents of 5-HT detected by liquid chromatography–tandem mass spectrometry (LC–MS/MS) in hippocampal neurons extracted from WT or AD mice incubated with *D-*NPs, *L-*NPs, and *rac-*NPs under NIR illumination (980 nm laser, 400 mW/cm^2^). Data are presented as mean ± s.d. (n = 5).

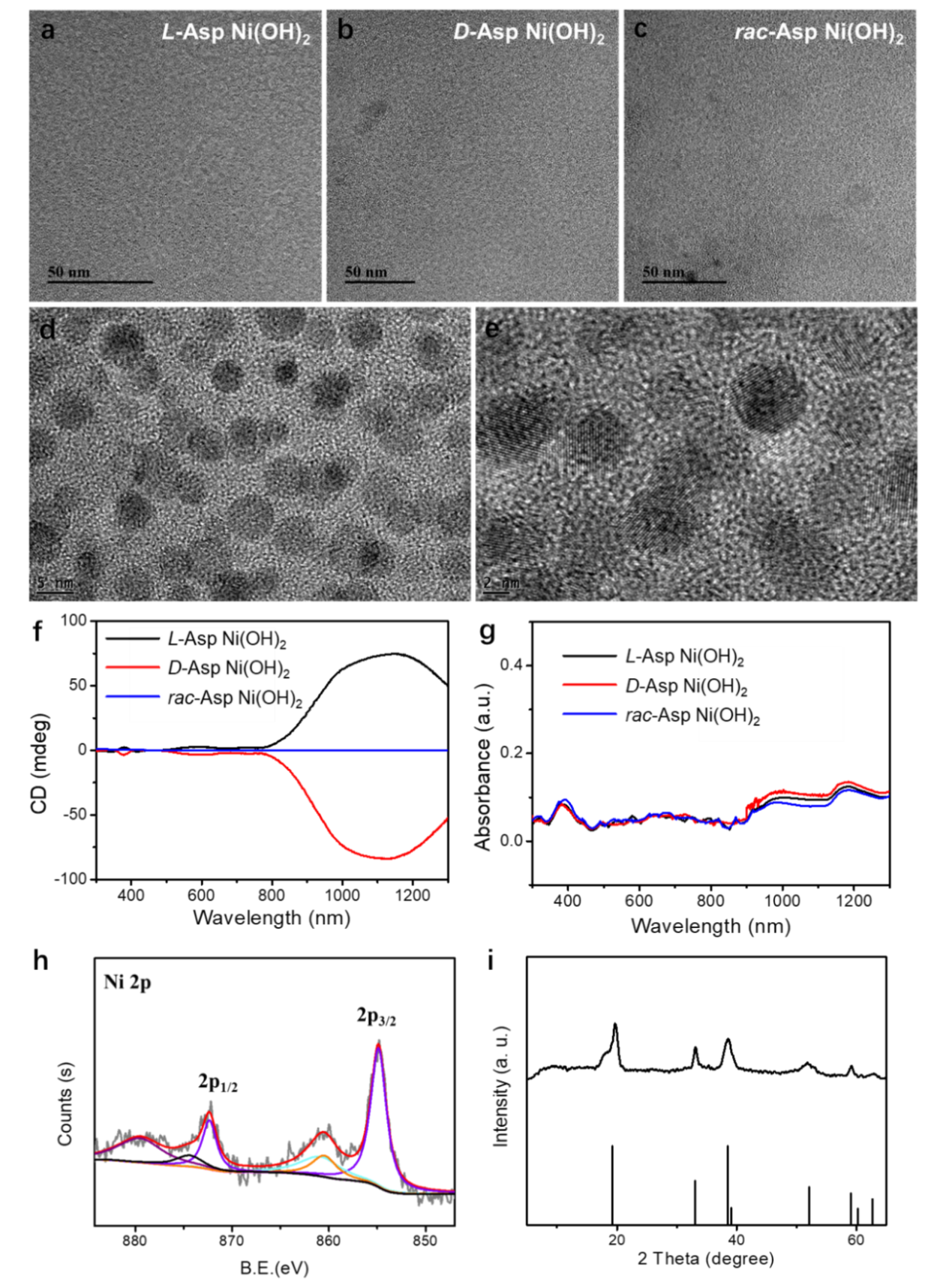

**Figure S40.** TEM images of **(a)** *L*-Asp-Ni(OH)_2_, **(b)** *D*-Asp-Ni(OH)_2_, and **(c)** *rac*-Asp-Ni(OH)_2_ NPs. **(d, e)** High resolution TEM image of *D*-Asp-Ni(OH)_2_ NPs. **(f)** CD and **(g)** UV-Vis spectra of chiral Asp-Ni(OH)_2_ NPs. **(h)** XPS spectra of chiral *D*-Asp-Ni(OH)_2_ NPs. **(i)** XRD spectra of chiral *D*-Asp-Ni(OH)_2_ NPs.

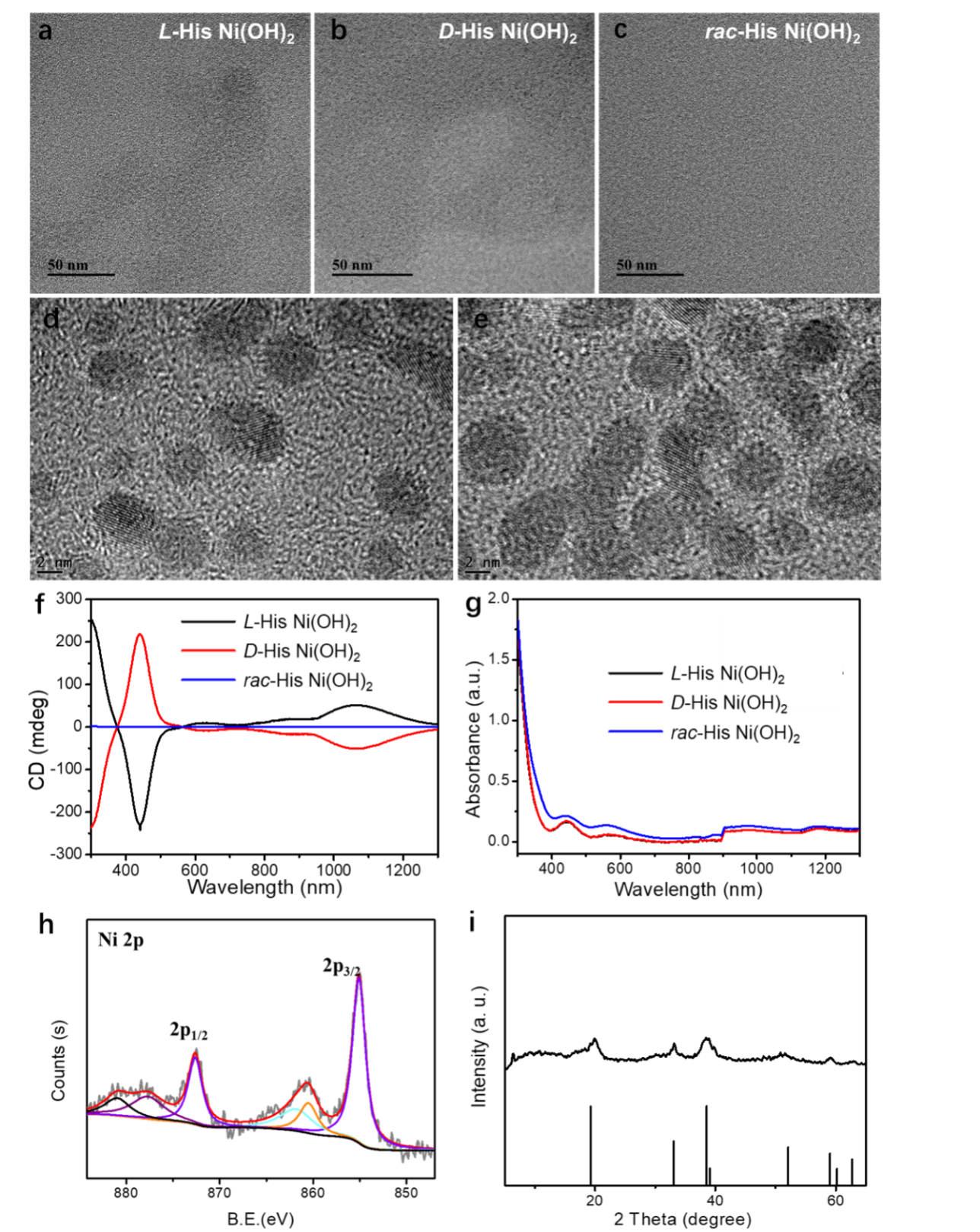

**Figure S41.** TEM images of **(a)** *L*-His-Ni(OH)_2_, **(b)** *D*-His-Ni(OH)_2_, and **(c)** *rac*-His-Ni(OH)_2_ NPs. **(d, e)** High resolution TEM image of *D*-His-Ni(OH)_2_ NPs. **(f)** CD and **(g)** UV-Vis spectra of chiral His-Ni(OH)_2_ NPs. **(h)** XPS spectra of chiral *D*-His-Ni(OH)_2_ NPs. **(i)** XRD spectra of chiral *D*-His-Ni(OH)_2_ NPs.

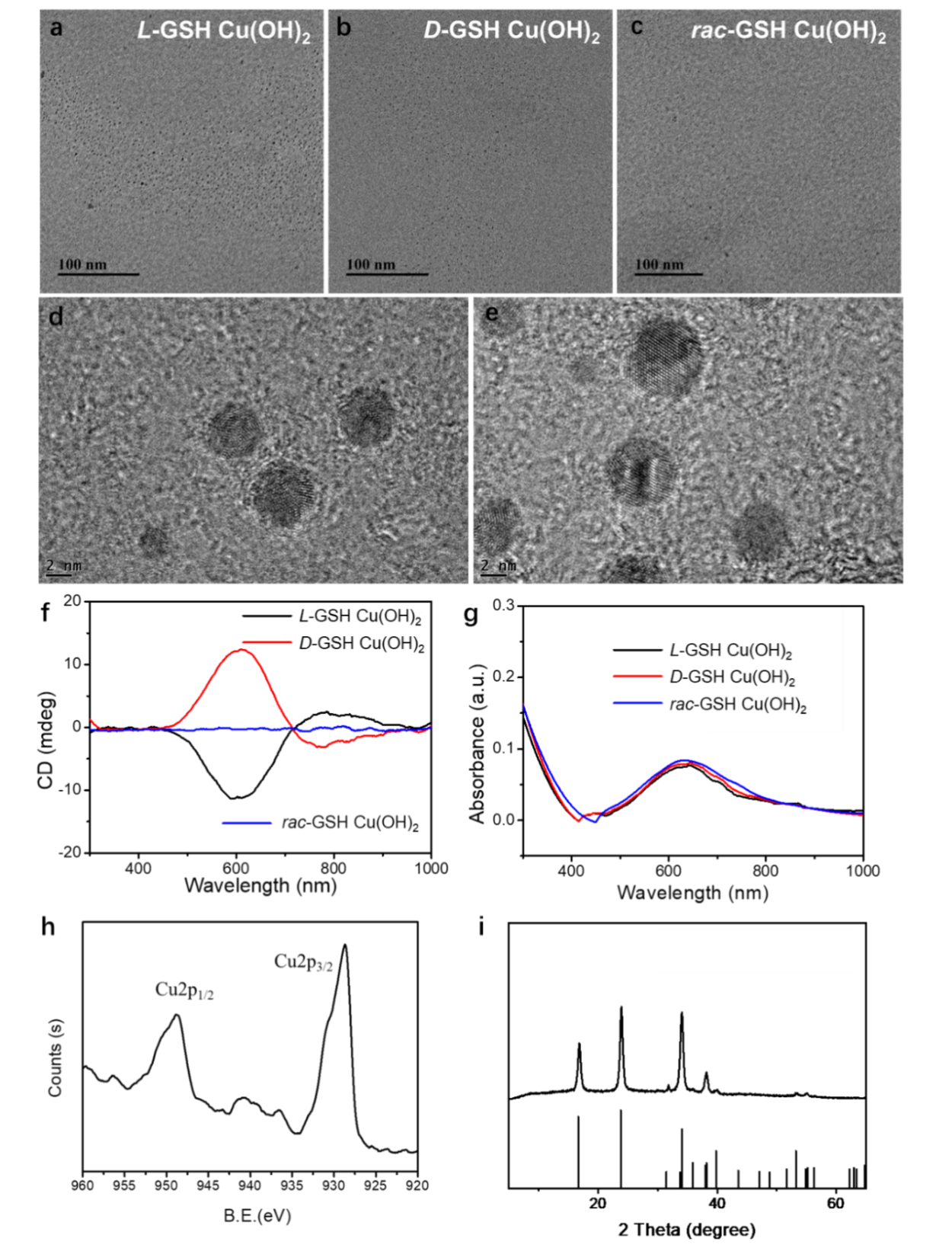

**Figure S42.** TEM images of **(a)** *L*-GSH-Cu(OH)_2_, **(b)** *D*-GSH-Cu(OH)_2_, and **(c)** *rac*-GSH-Cu(OH)_2_ NPs. **(d, e)** High resolution TEM image of *D*-GSH-Cu(OH)_2_ NPs. **(f)** CD and **(g)** UV-Vis spectra of chiral GSH-Cu(OH)_2_ NPs. **(h)** XPS spectra of chiral *D*-GSH-Cu(OH)_2_ NPs. **(i)** XRD spectra of chiral *D*-GSH-Cu(OH)_2_ NPs.

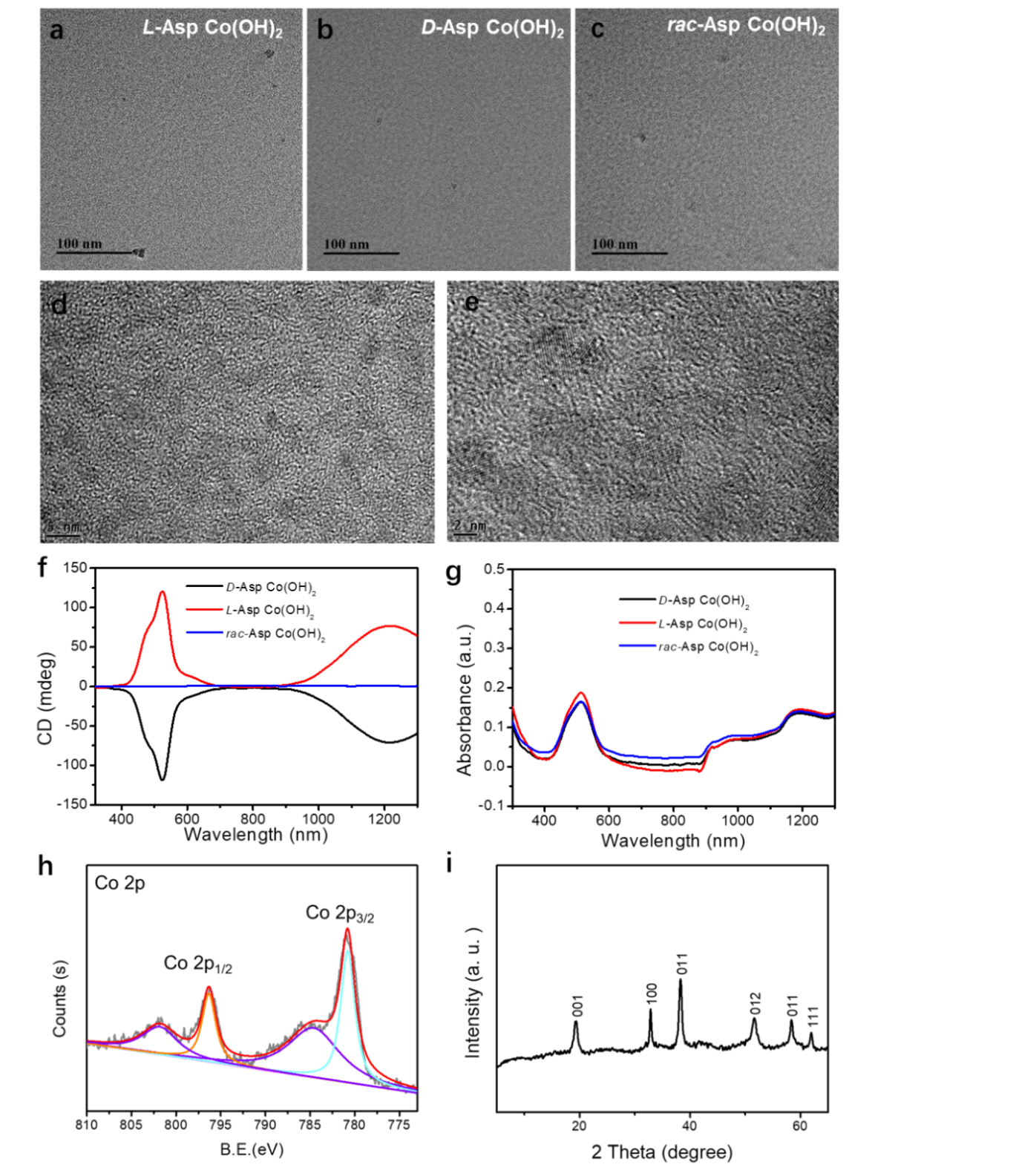

**Figure S43.** TEM images of **(a)** *L*-Asp-Co(OH)_2_, **(b)** *D*-Asp-Co(OH)_2_, and **(c)** *rac*-Asp-Co(OH)_2_ NPs. **(d, e)** High resolution TEM image of *D*-Asp-Co(OH)_2_ NPs. **(f)** CD and **(g)** UV-Vis spectra of chiral Asp-Co(OH)_2_ NPs. **(h)** XPS spectra of chiral *D*-Asp-Co(OH)_2_ NPs. **(i)** XRD spectra of chiral *D*-Asp-Co(OH)_2_ NPs.

**Figure S44. (a)** Confocal images of hippocampal neurons extracted from AD mice treated with Asp-Ni(OH)_2_ NPs and under NIR illumination (980 nm laser, 400 mW/cm^2^), or His-Ni(OH)_2_ NPs and under NIR illumination (980 nm laser, 400 mW/cm^2^), or GSH-Cu(OH)_2_ NPs and under 660 nm light illumination (660 nm laser, 400 mW/cm^2^), or Asp-Co(OH)_2_ NPs and under 532 nm light illumination (532 nm laser, 400 mW/cm^2^). Tuj-1 (green) to label neurons, and DAPI (blue) to stain nuclei. Scale bar, 50 μm. **(b)** Quantification of the axon length of hippocampal neurons (Tuj-1) extracted from AD mice with different treatments from **Figure S44a**. About 10-50 cells from any region of each well were used for average axon length statistics. Each experiment was repeated five times.

**Comments:** To test the specificity of GSH modified chiral Ni(OH)_2_ NPs on neuronal axon regeneration, other chiral hydroxides composed of different ligands or different elements, including Asp-Ni(OH)_2_ NPs, His-Ni(OH)_2_ NPs, GSH-Cu(OH)_2_ NPs, and Asp-Co(OH)_2_ NPs have been fabricated **(Figure S40-S43)**. These chiral NPs were incubated with neurons under light illumination, and there was no axon regeneration occurrence **(Figure S44)**.

**Figure S45.** Confocal imaging of hippocampus neurons incubated with *D*-Asp-Ni(OH)_2_ under NIR illumination (980 nm laser, 400 mW/cm^2^), *D*-His-Ni(OH)_2_ under NIR illumination (980 nm laser, 400 mW/cm^2^), *D*-GSH-Cu(OH)_2_ under 660 nm light illumination (660 nm laser, 400 mW/cm^2^), and *D*-Asp-Co(OH)_2_ under 532 nm light illumination (532 nm laser, 400 mW/cm^2^) for different times. Blue: DAPI, Red: *D*-/*L*-/*rac*-NPs-Cy5, Green: NGL-1-Cy3. Scale bar, 20 μm. About 10-50 cells from any region of each well were used for fluorescence intensity statistics. Each experiment was repeated five times.

**Figure S46. (a)** Confocal images of astrocytes, microglia, and hippocampal neurons incubated with *D*-NPs and under NIR illumination (980 nm laser, 400 mW/cm^2^) for different times or incubated with LPS as the positive group, detecting the ROS content using DCFH-DA oxidative stress indicator. **(b)** The DCFH-DA fluorescence intensity of astrocytes, microglia, and hippocampal neurons incubated with *D*-NPs and under NIR illumination for different times or incubated with LPS as the positive group. Scale bar: 50 μm, Data are presented as mean ± s.d. About 10-50 cells from any region of each well were used for fluorescence intensity statistics. Each experiment was repeated three times.

**Comments:** These results demonstrated that chiral *D*-NPs incubated with astrocytes, microglia, and hippocampal neurons cells do not produce any ROS under NIR illumination within 30min (980 nm laser, 400 mW/cm^2^).

**Figure S47. (a)** Confocal imaging of hippocampus neurons incubated with *D*-NPs under NIR illumination for different times. Blue: DAPI, Red: *D*-NPs-Cy5, Green: NetrinG1-Cy3. **(b)** Confocal images of hippocampus neurons after blocking NGL-1 by siRNA, and then incubated with *D*-NPs under NIR illumination. Blue: DAPI, Red: *D*-NPs-Cy5, Green: NGL-1-Cy3. Scale bar, 20 μm. **(c)** Confocal images of hippocampus neurons after blocking NetrinG1 by siRNA, and then incubated with *D*-NPs under NIR illumination. Blue: DAPI, Red: *D*-NPs-Cy5, Green: NGL-1-Cy3. **(d)** Mean fluorescence intensity of Cy5 for hippocampus neurons incubation with *D*-NPs under NIR light illumination. About 10-50 cells from any region of each well were used for fluorescence intensity statistics. Each experiment was repeated five times. Data are presented as mean ± s.d.

**Comments:** In **Figure S47a**, Fluorescence resonance energy transfer (FRET) experiment was designed to prove whether Netrin G1 has the same affinity with *D*-NPs as NGL-1. Here, two probes, *D*-NPs-Cy5 and NGL-1-Cy3, were prepared respectively. When *D*-NPs binds to Netrin G1, the fluorescence of Cy5 at 712 nm will be activated by the excitation of Cy3 at 550 nm because of the overlap between the emission wavelength of Cy3 at 561 nm and the excitation wavelength at 650 nm of Cy5. Based on this principle, it showed that the fluorescence intensity of Cy5 increased slightly as the incubation time extended, indicating lower bonding efficiency between *D*-NPs and Netrin G1.

As shown from **Figure S47b**, when we used siRNA to inhibit the expression of NGL-1, no *D*-NPs-Cy5 fluorescence appeared, indicating no *D*-NPs attached to the NGL-1. To be noticed, the FRET experiment showed that blocking Netrin G1 does not make too much difference on the binding between *D*-NP and NGL-1 (**Figure S47c**), supporting that *D*-NPs have a higher affinity for NGL-1 than for Netrin G1.

**Figure S48. (a)** Confocal images of hippocampus neurons pretreated with siIGF-1, and then incubated with *D*-NPs under NIR illumination. Tuj-1 (green) to label neurons, and DAPI (blue) to stain nuclei. **(b)** Confocal images of hippocampus neurons pretreated with siNGL-1, and then incubated with *D*-NPs under NIR illumination. **(c)** Confocal images of hippocampus neurons pretreated PD 98059 (ERK pathway inhibitor), and then incubated with *D*-NPs under NIR illumination. Green: IGF-1, Blue: DAPI. **(d)** Mean fluorescence intensity of IGF-1 in hippocampus neurons pretreated with PD 98059 (ERK pathway inhibitor), and then incubated with *D*-NPs under NIR illumination. About 10-50 cells from any region of each well were used for average axon length and fluorescence intensity statistics. Each experiment was repeated five times. **(e)** RT-PCR analysis for IGF-1 expression in hippocampus neurons pretreated with PD 98059 (ERK pathway inhibitor), and then incubated with *D*-NPs under NIR illumination. Data are presented as mean ± s.d.

**Figure S49. (a-c)** Confocal images of hippocampus neurons with *D*-NPs and NIR treatment; and pretreated with siNGL-2, or siNGL-3, or siNGL-1, siNGL-2, and siNGL-3, and then incubated with *D*-NPs under NIR illumination. Green: Tuj-1, Red: GAP43, Green: IGF-1, and Blue: DAPI. **(d)** Quantification the hippocampal neurons axon length with *D*-NPs and NIR treatment; and pretreated with siNGL-2, or siNGL-3, or siNGL-1, siNGL-2, and siNGL-3, and then incubated with *D*-NPs under NIR illumination from **Figure S49a**. **(e)** Mean fluorescence intensity of GAP43 in hippocampus neurons with *D*-NPs and NIR treatment; and pretreated with siNGL-2, or siNGL-3, or siNGL-1, siNGL-2, and siNGL-3, and then incubated with *D*-NPs under NIR illumination from **Figure S49b**. **(f)** Mean fluorescence intensity of IGF-1 in hippocampus neurons with *D*-NPs and NIR treatment; and pretreated with siNGL-2, or siNGL-3, or siNGL-1, siNGL-2, and siNGL-3, and then incubated with *D*-NPs under NIR illumination from **Figure S49c**. About 10-50 cells from any region of each well were used for average axon length and fluorescence intensity statistics. Each experiment was repeated five times. **(g)** RT-PCR analysis of IGF-1 expression in hippocampus neurons with *D*-NPs and NIR treatment; and pretreated with siNGL-2, or siNGL-3, or siNGL-1, siNGL-2, and siNGL-3, and then incubated with *D*-NPs under NIR illumination. Data are presented as mean ± s.d. (n = 5). **(h)** Western blot for GAP43, and IGF-1 expression in hippocampus neurons with *D*-NPs and NIR treatment; and pretreated with siNGL-1, or siNGL-2, or siNGL-3, or siNGL-1, siNGL-2, and siNGL-3, and then incubated with *D*-NPs under NIR illumination. Data are presented as mean ± s.d.

**Figure S50. (a)** 3D structures of NGL-2 and NGL-3 fragments. **(b)** Comparison of LRR sequences of NGL-1, NGL-2, and NGL-3.

**Comments:** As shown in **Figure S49**, the neurons from AD mice as the control group displayed short axon at about 50.2 ± 11.1 μm. We incubated the neurons with chiral *D*-NPs under NIR illumination and observed that the axon length displayed a marked increase to 207.2 ± 26.1 μm. We found that blocking NGL-2 and NGL-3 resulted in a much smaller effect on axon regeneration than blocking NGL-1. In addition, the expression of IGF-1 displayed a slight decrease after blocking NGL-2 and NGL-3, which was far less than that caused by blocking NGL-1. These results suggest that chiral nanoparticle-induced axon regeneration is mediated by NGL-1, but not NGL-2 and NGL-3. The reason is that the radius of curvature of the semispherical pocket in the leucine-rich repeat (LRR) segment in NGL-1, NGL-2, and NGL-3 are similar, but the differences are manifested in the sequences **(Figure S1 and Figure S50)**. Three LRR sequences have a difference of about 30% (*30-32*).

**Figure S51. (a)** ICP-MS analysis of the Ni element in the brain after intravenous injection with *D*-NPs (2 mg/kg, corresponding to Ni dose of 0.1mg/kg) using different methods, such as once through 90 days, and every five days until 60 days. **(b)** Photoacoustic images of *D*-NPs in brain for different times (70th, 80th, 90th day) after 60 days of intravenous injection, arrows represent *D*-NPs material signals. **(c)** ICP-MS analysis the Ni element in the brain after intravenous injection with *D*-NPs (2 mg/kg, corresponding to Ni dose of 0.1mg/kg) pretreated with different GSH transporter inhibitors. P-glycoprotein inhibitor: Zosuquidar; BCRP (breast cancer resistance protein) inhibitor: Ko143; MRP1 (multidrug resistance-associated proteins 1) inhibitor: MK571; MRP4 (multidrug resistance-associated proteins 4) inhibitor: Ceefourin 1; OAT (organic anion transport) inhibitor: Probenecid. **(d)** Photoacoustic images of kidney and liver tissue sections through intravenous injection with *D*-NPs (2 mg/kg, corresponding to a Ni dose of 0.1mg/kg) for different times. **(e-h)** ICP-MS analysis of the Ni element in the brain **(e)**, Blood **(f)**, Liver **(g)**, and kidney **(h)** after intravenous injection with *D*-NPs, *rac*-NPs, or *L*-NPs (2 mg/kg, corresponding to a Ni dose of 0.1mg/kg). Data are presented as mean ± s.d.

**Comments:** To illustrate the mechanism of chiral NPs traversing the BBB, they were pretreated with different GSH transporter inhibitors. ICP-MS results showed that when MRP1 and OAT were blocked, the content of *D*-NPs passing through the BBB and entering the brain is inhibited **(Figure S51c)**. This result indicated that the successful crossing of BBB by *D*-NP is mainly mediated by MRP1 and OAT. In addition, chiral NPs can enter the brain from the blood circulation, and then be excreted out of the body through the kidney and liver in the form of non-ionic nanoparticles **(Figure S51d-S51h)**. To improve the retention time of chiral NPs in the brain and the efficiency of axon regeneration, multiple intermittent intravenous injections (once every five days until 60 days) was used throughout the experiments **(Figure S51a)**.

**Figure S52. Toxicity analysis of chiral NPs and NiCl_2_. (a)** The cell viability of hippocampal neurons incubated with different concentrations of chiral *D*-NPs or NiCl_2_. **(b)** Representative histological analysis of different organs after tail vein injection of *D*-NPs, *L*-NPs, *rac*-NPs, or NiCl_2_, at the dose of 100 μg Ni/kg. **(c-f)** Serum biochemistry detection after tail vein injection of *D*-NPs, *L*-NPs, *rac*-NPs, or NiCl_2_, at the dose of 100 μg Ni/kg. ALT, alanine aminotransferase; AST, aspartate aminotransferase; BUN, blood urine nitrogen; CRE, creatinine. Control represents the mice without NPs injection. Data are presented as mean ± s.d. **(g)** Photograph showing embryotoxic and fetal malformations after tail vein injection with NiCl_2_ at the dose of 400 μg Ni/kg, or *D*-/*L*-/*rac*-NPs at the dose of 2000 μg Ni/kg. **(h)** Photograph showing skeletal abnormality of fetus (red arrow) after tail vein injection with NiCl_2_ at the dose of 400 μg Ni/kg, or *D*-NPs at the dose of 2000 μg Ni/kg.

**Comments:** When hippocampal neurons were incubated with chiral *D*-NPs or NiCl_2_ at the same Ni^2+^ concentration, the activity of neuronal cells treated with chiral NPs was higher than that of NiCl_2_ **(Figure S52a)**. *In vivo*, intravenous injection with chiral *D*-NPs (100 μg Ni/kg) was safe, as confirmed by histological analysis and serum biochemistry detection, but it caused obvious damage to liver and kidney tissue after injection with NiCl_2_ **(Figure S52b-S52f)**. Additionally, embryotoxic and fetal malformations data further illustrate that chiral NPs have no teratogenicity at a higher dose of 2000 μg Ni/kg, but a lower dose of NiCl_2_ (400 μg Ni/kg) caused embryonic agenesis **(Figure S52g, h)**. All above results showed that chiral Ni(OH)_2_ in the form of NPs avoid the biological toxicity caused by nickel ions.

**Figure S53.** A diagram illustrating the chiral NPs interaction with blood brain barrier and neurons under NIR illumination.

**Comments:** The chiral Ni(OH)_2_ NPs can effectively cross the blood-brain barrier through GSH transport, and then target the leucine-rich repeat (LRR) segment of the NGL-1 a transmembrane protein abundant in the dendrites of neurons (**Figure S53**). Under NIR illumination, it induced the conformational changes in NGL-1 and tightening of the NP-NGL-1 complex. Therefore, the downstream ERK and PI3K/Akt pathways were activated and induced the production of IGF-1, which thus promoted axon regeneration (**Figure 4m**).

**Figure S54. Uptake of *D*-NPs using the *in vitro* BBB model. (a)** Representative confocal images post treatment with two doses of 50 and 100 ng/mL *D*-NPs-Cy5 after 12 h on human brain endothelial hCMEC/D3 cells monolayer stained with DAPI representing nuclei (blue), claudin-5 antibody (CLD-5) (green), and zona-occludin antibody (ZO-1) (red). Scale bar 100 μm. **(b)** Representative confocal images post treatment with different GSH transporter inhibitor (MRP1, OAT, or MRP1 and OAT) for 24 h and then treatment with 100 ng/mL *D*-NPs-Cy5 for 12 h on human brain endothelial hCMEC/D3 cells monolayer stained with DAPI representing nuclei (blue), claudin-5 antibody (CLD-5) (green) and zona-occludin antibody (ZO-1) (red). Scale bar 100 μm.

**Comments:** We used human brain endothelial hCMEC/D3 cell lines to construct BBB model *in vitro*. The staining of claudin-5 antibody (CLD-5) (green) and zona-occludin antibody (ZO-1) (red) to label phenotypic presence of tight junctions. After establishing and validating the *in vitro* BBB model, *D*-NPs interaction with the BBB was studied. As shown from **Figure S54a**, a weaker fluorescent intensity present hCMEC/D3 cells monolayer when with 50 ng/mL of *D*-NPs treatment, which was enhanced at a higher dose of 100 ng/mL. This phenomenon showed an efficient uptake of *D*-NPs in vitro BBB model with a concentration dependent manner. However, the uptake was inhibited when treated with GSH transporter (MRP1 and OAT) inhibitors, demonstrating the crossing of BBB mediated by GSH transporters **(Figure S54b)**.

**Figure S55. (a)** Nissl staining of neuron cells in the brains (hippocampus) of WT and AD mice with different treatments. **(b, c)** Immunofluorescence of Aβ **(b)** and p-Tau **(c)** in the hippocampus of WT and AD mice with different treatments. **(d, e)** Representative immunostaining of hippocampal sections for Aβ **(d)** and p-Tau **(e)** protein aggregates of WT and AD mice with different treatments. **(f, g)** Mean fluorescence intensity of Aβ **(f)** and p-Tau **(g)** in the hippocampus of WT and AD mice with different treatments from **Figure S55b, c**. **(h, i)** Quantitative analysis of Aβ **(h)**, and p-Tau **(i)** load in the hippocampus of WT and AD mice with different treatments from **Figure S55d, e**. Data are presented as mean ± s.d. (n = 5).

**Figure S56. (a)** Confocal images of DRG neurons treated with *D*-NPs only, or only under NIR illumination (980 nm laser, 400 mW/cm^2^). Tuj-1 (green) to label neurons. Scale bar, 50 μm. **(b)** Quantification of the axonal length of DRG neurons treated with *D*-NPs only, or only under NIR illumination from **Figure S56a**. About 10-50 cells from any region of each well were used for average axon length. Each experiment was repeated five times. Data are presented as mean ± s.d.

**Figure S57. (a)** Global gene-expression pattern of transcription factors in DRG neurons treated with *D*-NPs and under NIR illumination, (n = 3). **(b)** RT-PCR analysis for IGF-1, AKT, PI3K, and ERK expression in DRG neurons treated with *D*-NPs and under NIR illumination. the DRG neurons without any treatment were used as the control group. Data are presented as mean ± s.d. (n = 5). *P < 0.05, **P < 0.01, ***P < 0.001.

**Comments:** The upregulation gene of DRG neurons treated with *D*-NPs and NIR were including IGF-1, AKT, PI3K, ERK, and neuron-associated markers is observed, which were consistent with the changes of gene expression in hippocampal neurons and also verified the developed pathway.

**Figure S58. (a)** Confocal images of DRG neurons pretreated with siNGL-1, and then incubated with *D*-NPs under NIR illumination. Green: IGF-1. Scale bar, 50 μm. **(b)** Mean fluorescence intensity of IGF-1 in DRG neurons pretreated with siNGL-1, and then incubated with *D*-NPs under NIR illumination from **Figure S58a**. About 10-50 cells from any region of each well were used for fluorescence intensity statistics. Each experiment was repeated five times. Data are presented as mean ± s.d.

**Figure S59.** **(a)** Representative footprints of healthy mice (Control), SCI mice, and SCI mice with *D*-NPs under NIR treatment (2 mg/kg, corresponding to Ni dose of 0.1 mg/kg), or intraperitoneal injection (10 mg/kg/day) of IGF-1. **(b)** Representative photographs of mice hind limb positioning when walking in different groups mentioned in **Figure S59a**. At least five animals per condition.

**Figure S60. (a)** Representative footprints of SCI mice spontaneous recovery for 60 days after injury. **(b, c)** Bar graphs representing **(b)** stride length in mm and **(c)** stride width in mm of mice mentioned spontaneous recovery for 60 days after injury. Data are presented as mean ± s.d. (n = 5). *P < 0.05, **P < 0.01, ***P < 0.001.

**Figure S61.** Magnetic resonance imaging of SCI mice with or without *D*-NPs and NIR treatment, or spontaneous recovery for 60 days after injury, the red circle represents the injured sites (n = 5).

**Figure S62.** Partial enlargement within the red lines of magnetic resonance imaging from **Figure S61.** The SCI mice with or without *D*-NPs and NIR treatment, or spontaneous recovery for 60 days after injury, the red circle represents the injured sites (n = 5).

**Figure S63. (a, b)** ICP-MS analysis of the Ni element in **(a)** breast and in **(b)** thyroid tissues after intravenous injection with *D*-NPs (2 mg/kg, corresponding to a Ni dose of 0.1mg/kg) at different time points. **(c-e)** Quantitative detection of IGF-1 concentration in breast **(c)**, thyroid **(d)**, and blood **(e)** from AD mice without treatment or with *D*-NPs and NIR treatment. **(f-h)** Detection of insulin **(f)**, blood sugar **(g)**, and hemoglobin A1c (HbA1c) **(h)** in AD mice without treatment or with D-NPs and NIR treatment. Data are presented as mean ± s.d. (n = 5).

**Comments:** Systemic administration of IGF-1 increases the risk of hypoglycemia^30^ as well as breast and thyroid cancers,^31^ limiting its potential as an AD treatment. When binding to NGL-1 on neurons, *D*-NPs do not accumulate in risk organs such as breast and thyroid **(Figure** **S63a, b)**. Similarly, breast and thyroid tissues showed no increase in IGF-1 concentration following treatment with *D*-NPs under NIR illumination. **(Figure S63c, d)**. Although IGF-1 levels in the blood stream showed a slight increase **(Figure S63e)**, they remained within the normal range, indicating low risk of hypoglycemia. Insulin, blood sugar, and HbA1c levels were also within normal ranges **(Figure S63f-63h)**.

**Table S1.** Average values of the Osipov-Pickup-Dunmur (OPD) chirality index calculated using the Ni and O coordinates of 15 structures of the cluster separated by 10 ns in the range between 60 and 200 ns of each trajectory.

| NPs | OPD value |
| --- | --- |
| *D*-GSH | -0.0156 ± 0.0119 |
| *L*-GSH | -0.0014 ± 0.0068 |
| *rac*-GSH | 0.0029 ± 0.0031 |
| No-GSH | -0.0005 ± 0.0007 |

**Note:** The differences between the averages for *rac*-GSH and *L*-GSH is small but it is statistically significant (p-value of 0.034 in a two sample T-test), while the difference between *D*-GSH and *L*-GSH is even more significant (p-value of 0.0004062).

**Table S2.** Protein residues for which individual hydrogen bonds with *D*-GSH-NP presented occupancy values (Occ) larger than 30%. The chemical group involved in the hydrogen bond as donor or acceptor are indicated between parentheses (in some cases the force field type is indicated for disambiguation) and for the protein it is also indicated whether the group belongs to the backbone or to the sidechain.

| Protein residue/  type | Total Occ (%) | Details | | |
| --- | --- | --- | --- | --- |
|  |  | Donor | Acceptor | Partial Occ (%) |
| ASP178/  Charged (-1) | 95.4 | DGSH (COOH from DGLY27)  DGSH (COOH from DGLY27) | ASP178 (OD1 from sidechain)  ASP178 (OD2 from sidechain) | 86.8 8.6 |
| ASP133/  Charged (-1) | 75.2 | DGSH (COOH from DGLY23)  DGSH (COOH from DGLY23) | ASP133 (OD1 from sidechain)  ASP133 (OD2 from sidechain) | 64.2  11.0 |
| GLU223/  Charged (-1) | 71.8 | DGSH (COOH from DGLY28)  DGSH (COOH from DGLY28)  Others | GLU223 (OE1 from sidechain)  GLU223 (OE2 from sidechain)  Others | 39.1  31.8  < 1.0 |
| GLU271/  Charged (-1) | 71.7 | DGSH (COOH from DGLY33)  DGSH (COOH from DGLY33) | GLU271 (OE1 from sidechain)  GLU271 (OE2 from sidechain) | 36.0  35.7 |
| GLU181/  Charged (-1) | 68.4 | DGSH (COOH from DGLU31)  DGSH (COOH from DGLU31)  Others | GLU181 (OE2 from sidechain)  GLU181 (OE1 from sidechain)  Others | 33.3  32.4  < 2.7 |
| GLU152/  Charged (-1) | 52.8 | DGSH (COOH from DGLU25)  DGSH (COOH from DGLU25)  Others | GLU152 (OE2 from sidechain)  GLU152 (OE1 from sidechain)  Others | 27.0  25.5  < 0.3 |
| THR324/  Polar | 37.0 | DGSH (NH2 from DGLU36)  Others | THR324 (O from backbone) Others | 35.3  < 1.8 |

**Table S3.** Protein residues for which individual hydrogen bonds with *L*-GSH-NP presented occupancy values (Occ) larger than 30%. The chemical group involved in the hydrogen bond as donor or acceptor are indicated between parentheses (in some cases the force field type is indicated for disambiguation) and for the protein it is also indicated whether the group belongs to the backbone or to the sidechain.

| Protein residue/ type | Total Occ (%) | Details | | |
| --- | --- | --- | --- | --- |
|  |  | Donor | Acceptor | Partial Occ (%) |
| ASN323/ polar | 79.3 | ASN323 (ND2 from sidechain)  ASN323 (ND2 from sidechain)  LGSH (N from LGLU1)  ASN323 (N from backbone)  ASN323 (ND2 from sidechain)  Others | LGSH (O from LCYS6)  LGSH (O from LCYS1)  ASN323 (OD1 from sidechain)  LGSH (N from LGLU1)  LGSH (O from LGLU1)  Others | 58.2  13.1  2.0  1.4  1.3  < 3.3 |
| GLU152/  charged (-1) | 76.3 | LGSH (COOH from LGLY18)  LGSH (COOH from LGLY18)  LGSH (COOH from LGLY15)  LGSH (COOH from LGLY15) | GLU152 (OE1 from sidechain)  GLU152 (OE2 from sidechain)  GLU152 (OE1 from sidechain)  GLU152 (OE2 from sidechain) | 33.3  27.5  8.2  7.3 |
| GLU181/ charged (-1) | 35.4 | LGSH (COOH from LGLY11)  LGSH (COOH from LGLY11) | GLU181 (OE2 from sidechain)  GLU181 (OE1 from sidechain) | 17.8  17.6 |
| SER322/ polar | 34.9 | LGSH (COOH from LGLU1)  LGSH (NH2 from LGLU1)  Others | SER322 (OG from sidechain)  SER322 (OG from sidechain)  Others | 30.6  2.5  < 1.8 |
| GLU271/ charged | 31.8 | LGSH (COOH from LGLU7)  LGSH (COOH from LGLU7)  Others | GLU271 (OE2 from sidechain)  GLU271 (OE1 from sidechain)  Others | 15.5  15.5  < 0.8 |
| ARG329/ charged (+1) | 31.7 | ARG329 (NH1 from sidechain)  ARG329 (NH2 from sidechain)  ARG329 (NH2 from sidechain)  ARG329 (NE from sidechain)  ARG329 (NH1 from sidechain)  Others | LGSH (COOH from LGLU4)  LGSH (NH2 from LGLU4)  LGSH (COOH from LGLU4)  LGSH (COOH from LGLU4)  LGSH (NH2 from LGLU4)  Others | 9,8  7.5  7.1  4.9  2.2  < 0.3 |
| GLN106/  polar | 30.0 | LGSH (COOH from LGLU40)  GLN106 (NE2 from sidechain)  LGSH (COOH from LGLU18)  GLN106 (NE2 from sidechain)  Others | GLN106 (OE1 from sidechain)  LGSH (O from LCYS18)  GLN106 (OE1 from sidechain)  LGSH (N2 from LGLU40)  Others | 17.6  4.4  2.5  2.0  < 3.5 |

**Table S4.** Protein residues for which individual hydrogen bonds with *rac*-GSH-NP presented occupancy values (Occ) larger than 30%. The chemical group involved in the hydrogen bond as donor or acceptor are indicated between parentheses (in some cases the force field type is indicated for disambiguation) and for the protein it is also indicated whether the group belongs to the backbone or to the sidechain.

| Protein residue/ type | Total Occ (%) | Details | | |
| --- | --- | --- | --- | --- |
|  |  | Donor | Acceptor | Partial Occ (%) |
| ARG64/ charged (+1) | 144.2 | ARG64 (NH2 from sidechain)  ARG64 (NH1 from sidechain)  Others | DGSH (NH2 from DGLU29)  DGSH (COOH from DGLU29)  Others | 81.4  60.2  < 2.6 |
| ARG156/ charged (+1) | 95.9 | ARG156 (NH1 from sidechain)  ARG156 (NH2 from sidechain)  ARG156(NH2 from sidechain)  ARG156(NH2 from sidechain)  Others | LGSH (NH2 from LGLU5)  LGSH (COOH from LGLY6)  LGSH (NH2 from LGLU5)  LGSH (COOH from LGLY6)  Others | 72.5  14.6  5.7  3.1  < 0.2 |
| GLU181/ charged (-1) | 89.0 | LGSH (COOH from LGLY6)  LGSH (COOH from LGLY6) | GLU181 (OE1 from sidechain)  GLU181 (OE2 from sidechain) | 55.1  33.9 |
| GLU130/ charged (-1) | 78.3 | DGSH (NH2 from DGLU27)  DGSH (NH2 from DGLU27)  DGSH (COOH from DGLU27) | GLU130 (OE1 from sidechain)  GLU130 (OE2 from sidechain)  GLU130 (OE2 from sidechain) | 42.5  33.8  2.0 |
| TRP154/ nonpolar | 65.0 | TRP154 (NE1 from sidechain)  Others | DGSH (COOH from DGLU27)  Others | 64.2  < 0.8 |
| GLU271/ charged (-1) | 57.5 | LGSH (COOH from LGLY15)  LGSH (COOH from LGLY15)  DGSH (COOH from DGLY35)  DGSH (COOH from DGLY35) | GLU271 (OE2 from sidechain)  GLU271 (OE1 from sidechain)  GLU271 (OE2 from sidechain)  GLU217 (OE1 from sidechain) | 29.0  25.3  1.7  1.5 |
| GLN106/ polar | 46.0 | GLN106 (NE2 from sidechain)  Others | LGSH (NH2 from LGLU11)  Others | 45.2  < 0.8 |
| GLU223/ charged (-1) | 38.3 | DGSH (COOH from DGLY35)  DGSH (COOH from DGLY35) | GLU223 (OE2 from sidechain)  GLU223 (OE1 from sidechain) | 20.4  17.9 |
| ASN273/ polar | 38.0 | ASN273 (ND2 from sidechain)  Others | LGSH (COOH from LGLY15)  Others | 37.6  < 0.4 |

**Notes:** Taking 30% total occupancy as the threshold for the analysis, we found that there are seven amino acid residues on *D*-GSH-NP (**Table S2**) and *L*-GSH-NP (**Table S3**) interacting with the protein, while there are nine amino acid residues on *rac*-GSH-NP (**Table S4**) involved in the same interactions. Most of these amino acids bear charged side-chains (ASP, GLU or ARG).

**Table S5.** Lowest energy transitions calculated for the NP/protein complexes and for the protein fragment from the *D*-GSH NP system.

| System | Wavelength (nm) | Energy (kJ/mol) | Oscillator strength  (L mol^-1^ cm^-1^) | Rotatory strength  (L mol^-1^ cm^-1^) |
| --- | --- | --- | --- | --- |
| *D*-GSH NP + protein | 1454.8 | 82.2 | 3661 | 2.690 |
| *L*-GSH NP + protein | 1015.2 | 117.8 | 9175 | 2.189 |
| *rac*-GSH NP + protein | 1175.6 | 101.8 | 8506 | -4.3040 |
| protein | 350.9 | 340.9 | 28497 | -0.7864 |

**Table S6**. Initial bond parameters for the atoms of the Ni(OH)_2_ NP.

| Atom i | Atom j | Bond length (nm) | Force constant (kJ/mol/nm²) |
| --- | --- | --- | --- |
| Ni | S | 0.1950 | 250000^a^ |
| Ni | O | 0.2055^b^ | 100000^c^ |
| O | H | 0.0980^b^ | 456000^d^ |

^a^ Initial value based on Fe-S from Charmm36 HEME group.

^b^ Initial crystallographic values; all bonds were set exactly to the same value.

^c^ Initial value that is expected to be smaller than the value for Fe-O from Charmm36, which yielded an excessively rigid NP.

^d^ Value for O-H bond from hydroxyl (Charmm36).

**Table S7**. Additional bonded parameters for the Ni atoms of the Ni(OH)_2_ NP obtained after initial relaxation.

| Atom i | Atom j | Bond length (nm) | Force constant (kJ/mol/nm²) |
| --- | --- | --- | --- |
| Ni | Ni | Distance after relaxation ^a^ | 50000^b^ |

^a^ Different bonding patterns were detected within layers and between layers, with Ni-Ni distances ranging from 0.41 to 0.56 nm.

^b^ The small force constant was used only to prevent the layer separation along the full MD in water and to enforce the chirality induced by the ligands.

**Table S8.** Quantitative RT-PCR primer sequence.

| Gene name | Sequence |
| --- | --- |
| *GAP43* | Forward: AAGGCAGGGGAAGATACCAC  Reverse: TTGTTCAATCTTTTGGTCCTCAT |
| *GFAP* | Forward: AAGCAGATGAAGCCACCCTG  Reverse: GTCTGCACG-GGAATGGTGAT |
| *AKT* | Forward: ATGTCCGAGATCCTACCCTACG  Reverse: AGCGAAGAAGGAGTTGGTGTC |
| *IGF-1* | Forward: GGCCGACTTCACTGTACAACCG  Reverse: GGTCACAAGCCAGTCCTCTTACTTC |
| *PI3K* | Forward: GGAACCAACTTCAGCAGCTC  Reverse: GAGCCTCTCTGTCCTGTTGG |
| *ERK* | Forward: CCAGGAAAGCATTACCTTGACC  Reverse: CCAGAGCCTGTTCAACTTCAATC |
| *Timp4* | Forward: TACACGCCATTTGACTCTT  Reverse: TGGTTCCTGGTCCCTACT |
| *GAPDH* | Forward: CTCGTCCCGTAGACAAAATG  Reverse: TGAGGTCAATGAAGGGGTCGT |
| *Steap4* | Forward: TGACTCTCCTTCCAGATCCCA  Reverse: TGCCCACACTAGGCTGACA |
| *Cxcl10* | Forward: CCAAGTGCTGCCGTCATTTTC |
|  | Reverse: GGCTCGCAGGGATGATTTCAA |
| *Fbln5* | Forward: CTGTGACCCAGGATATG AACTT |
|  | Reverse: TTGTAAATTGTAGCACGTCTGC |
| *Gbp2* | Forward: GGGGTCACTGTCTGACCACT |
|  | Reverse: ATGCATTCTGCACACAGAGG |
| *Serpina3n* | Forward: GGACATTGATGGTGCTGGTGAATTA |
|  | Reverse: CTCCTCTTGCCCGCGTAGAA |
| *Ggta1* | Forward: ACCGATTCTGCTGAAGACCT |
|  | Reverse: CAAACAGCAGAGCAACCGAG |

**References**

1.	M. J. Abraham *et al.*, GROMACS: High performance molecular simulations through multi-level parallelism from laptops to supercomputers. *SoftwareX* **1-2**, 19-25 (2015).

2.	D. Van Der Spoel *et al.*, GROMACS: fast, flexible, and free. *J. Comput. Chem.* **26**, 1701-1718 (2005).

3.	S. Páll, M. J. Abraham, C. Kutzner, B. Hess, E. Lindahl, Tackling Exascale Software Challenges in Molecular Dynamics Simulations with GROMACS. *ArXiv* **8759**, 3–27 (2015).

4.	T. Darden, D. York, L. Pedersen, Particle mesh Ewald: An N⋅log(N) method for Ewald sums in large systems. *J. Chem. Phys.* **98**, 10089-10092 (1993).

5.	U. Essmann *et al.*, A smooth particle mesh Ewald method. *J. Chem. Phys.* **103**, 8577-8593 (1995).

6.	B. Hess, H. Bekker, H. J. C. Berendsen, J. G. E. M. Fraaije, LINCS: A linear constraint solver for molecular simulations. *J. Comput. Chem.* **18**, 1463-1472 (1997).

7.	VMD: Visual Molecular Dynamics. *J. Mol. Graph.* **14**, 33-38 (1996).

8.	J. Huang, A. D. MacKerell, Jr., CHARMM36 all-atom additive protein force field: validation based on comparison to NMR data. *J. Comput. Chem.* **34**, 2135-2145 (2013).

9.	S. Grimme, C. Bannwarth, P. Shushkov, A Robust and Accurate Tight-Binding Quantum Chemical Method for Structures, Vibrational Frequencies, and Noncovalent Interactions of Large Molecular Systems Parametrized for All spd-Block Elements (Z = 1-86). *J. Chem. Theory Comput.* **13**, 1989-2009 (2017).

10.	C. Bannwarth *et al.*, Extended tight-binding quantum chemistry methods. *WIREs COMPUT. MOL. SCI.* **11**, e1493 (2020).

11.	K. Momma, F. Izumi, VESTA 3 for three-dimensional visualization of crystal, volumetric and morphology data. *J. Appl. Crystallogr.* **44**, 1272-1276 (2011).

12.	S. Grazulis *et al.*, Crystallography Open Database - an open-access collection of crystal structures. *J. Appl. Crystallogr.* **42**, 726-729 (2009).

13.	S. Grazulis *et al.*, Crystallography Open Database (COD): an open-access collection of crystal structures and platform for world-wide collaboration. *Nucleic Acids Res.* **40**, D420-427 (2012).

14.	V. Zoete, M. A. Cuendet, A. Grosdidier, O. Michielin, SwissParam: a fast force field generation tool for small organic molecules. *J. Comput. Chem.* **32**, 2359-2368 (2011).

15.	J. C. Lin, W.-H. Ho, A. Gurney, A. Rosenthal, The netrin-G1 ligand NGL-1 promotes the outgrowth of thalamocortical axons. *Nat. Neurosci.* **6**, 1270-1276 (2003).

16.	M. H. M. Olsson, C. R. Søndergaard, M. Rostkowski, J. H. Jensen, PROPKA3: Consistent Treatment of Internal and Surface Residues in Empirical pKa Predictions. *J. Chem. Theory Comput.* **7**, 525-537 (2011).

17.	F. Mariano Colombari, A. Lozada-Blanco, K. Bernardino, W. R. Gomes, A. Farias de Moura, Themis: A Software to Assess Association Free Energies via Direct Estimative of Partition Functions. *chemrxiv*, doi:10.26434/chemrxiv. 12925550.v12925552 (2020).

18.	H. J. C. Berendsen, J. P. M. Postma, W. F. van Gunsteren, A. DiNola, J. R. Haak, Molecular dynamics with coupling to an external bath. *J. Chem. Phys.* **81**, 3684-3690 (1984).

19.	M. Parrinello, A. Rahman, Polymorphic transitions in single crystals: A new molecular dynamics method. *J. Appl. Phys.* **52**, 7182-7190 (1981).

20.	S. Grimme, C. Bannwarth, Ultra-fast computation of electronic spectra for large systems by tight-binding based simplified Tamm-Dancoff approximation (sTDA-xTB). *J. Chem. Phys.* **145**, 054103 (2016).

21..	C. Bannwarth, S. Ehlert, S. Grimme, GFN2-xTB—An Accurate and Broadly Parametrized Self-Consistent Tight-Binding Quantum Chemical Method with Multipole Electrostatics and Density-Dependent Dispersion Contributions. *J. Chem. Theory Comput.* **15**, 1652-1671 (2019).

22.	G. J. Brewer, J. R. Torricelli, Isolation and culture of adult neurons and neurospheres. *Nat. Protoc.* **2**, 1490-1498 (2007).

23.	G. Kong *et al.*, AMPK controls the axonal regenerative ability of dorsal root ganglia sensory neurons after spinal cord injury. *Nat. Metab.* **2**, 918-933 (2020).

24.	Z. Ahmed *et al.*, Disinhibition of neurotrophin-induced dorsal root ganglion cell neurite outgrowth on CNS myelin by siRNA-mediated knockdown of NgR, p75NTR and Rho-A. *Mol. Cell. Neurosci.* **28**, 509-523 (2005).

25.	A. Kovac, Z. Somikova, N. Zilka, M. Novak, Liquid chromatography–tandem mass spectrometry method for determination of panel of neurotransmitters in cerebrospinal fluid from the rat model for tauopathy. *Talanta* **119**, 284-290 (2014).

26.	B. A. Cisterna *et al.*, Active acetylcholine receptors prevent the atrophy of skeletal muscles and favor reinnervation. *Nat. Commun.* **11**, 1073 (2020).

27.	Z. Álvarez *et al.*, Bioactive scaffolds with enhanced supramolecular motion promote recovery from spinal cord injury. *Science* **374**, 848-856 (2021).

28.	C. E. McNamee, S. Armini, S. Yamamoto, K. Higashitani, Determination of the binding of non-cross-linked and cross-linked gels to living cells by atomic force microscopy. *Langmuir* **25**, 6977-6984 (2009).

29.	J. K. Vasir, V. Labhasetwar, Quantification of the force of nanoparticle-cell membrane interactions and its influence on intracellular trafficking of nanoparticles. *Biomaterials* **29**, 4244-4252 (2008).

30	F. Fang, J. L. Goldstein, X. M. Shi, G. S. Liang, M. S. Brown, Unexpected role for IGF-1 in starvation: Maintenance of blood glucose. Proc. *Natl. Acad. Sci. U S A* **119**, e2208855119 (2022).

31	L. Manzella et al., Activation of the IGF Axis in Thyroid Cancer: Implications for Tumorigenesis and Treatment. *Int. J. Mol. Sci.* **20**, 3258 (2019).

32. https://www.uniprot.org/uniprotkb/Q8C031/entry#sequences.

33. https://www.uniprot.org/uniprotkb/Q99PH1/entry#sequences.

34. https://www.uniprot.org/uniprotkb/P0C192/entry#sequences.
